# Mendelian randomization analysis with pleiotropy-robust log-linear model for binary outcomes

**DOI:** 10.1101/2023.06.03.543587

**Authors:** Mintao Li, Tao Huang, Jinzhu Jia

**Affiliations:** Department of Biostatistics, School of Public Health, Peking University, Beijing 100191, China; Department of Epidemiology & Biostatistics, School of Public Health, Peking University, Beijing 100191, China; Key Laboratory of Molecular Cardiovascular Sciences (Peking University), Ministry of Education, Beijing 100191, China; Center for Statistical Science, Peking University, Beijing 100871, China

**Author notes:** Corresponding authors: Jinzhu Jia, Tao Huang.

**Keywords:** Mendelian randomization, pleiotropy, binary exposure, binary outcome, risk ratio

## Abstract

Mendelian randomization (MR) is a statistical technique that uses genetic variants as instrumental variables to infer causality between traits. In dealing with a binary outcome, there are two challenging barriers on the way toward a valid MR analysis, that is, the inconsistency of the traditional ratio estimator and the existence of horizontal pleiotropy. Recent MR methods mainly focus on handling pleiotropy with summary statistics. Many of them cannot be easily applied to one-sample MR. We propose two novel individual data-based methods, respectively named random-effects and fixed-effects MR-PROLLIM, to surmount both barriers. These two methods adopt risk ratio (RR) to define the causal effect for a continuous or binary exposure. The random-effects MR-PROLLIM models correlated pleiotropy, accounts for variant selection, and allows weaker instruments. The fixed-effects MR-PROLLIM can function with only a few selected variants. We demonstrate in this study that the random-effects MR-PROLLIM exhibits high statistical power while yielding fewer false-positive detections than its competitors. The fixed-effects MR-PROLLIM generally performs at an intermediate level between the classical median and mode estimators. In our UK Biobank data analyses, we also found (i) the MR ratio method tended to underestimate binary exposure effects to a large extent; (ii) about 26.5% of the trait pairs were detected to have significant correlated pleiotropy; (iii) the pleiotropy-sensitive method showed estimated relative biases ranging from -103.7% to 178.0% for inferred non-zero effects. MR-PROLLIM exhibits the potential to facilitate a more rigorous and robust MR analysis for binary outcomes.

## Introduction

Uncovering causal connections between traits is the primary objective of many biomedical and epidemiological studies. Different types of research can provide evidence with differential levels for causality (1). The randomized controlled trial (RCT) has long been regarded as the gold standard in causal inference. However, restricted by ethical issues and research costs, RCT is not always practicable. It is generally easier to conduct an observational study, but the results may be biased due to unmeasured confounders or reverse causality. The application of instrumental variables (IVs) has been shown in statistical theory and econometrical practice as a useful approach to surmount these shortcomings (2). One biomedical analog to IV analysis is the Mendelian randomization (MR), which treats genetic variants, typically single nucleotide polymorphisms (SNPs), as IVs for certain exposures and outcomes (3).

Currently, the commonly used statistical methods for IV or MR analysis originate from linear models, which assume linear associations among the SNP *G*, exposure *X*, and outcome *Y*, with the following three IV assumptions: (i) *G* is correlated with *X*; (ii) *G* is independent of the confounder *U*; and (iii) given *X* and *U, G* is independent of *Y* (i.e., the exclusion restriction assumption). **Fig. 1A** shows these IV assumptions, and if there is a set of control variables **C**, all these three assumptions should be based on holding **C** constant (4). The traditional ratio estimator (also known as the Wald estimator 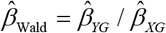, where 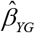 and 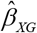 are the effect estimators of *G* on *Y* and *X*, respectively) works fine when both the exposure and outcome are continuous variables, and the linear associations hold.

**Fig. 1.**
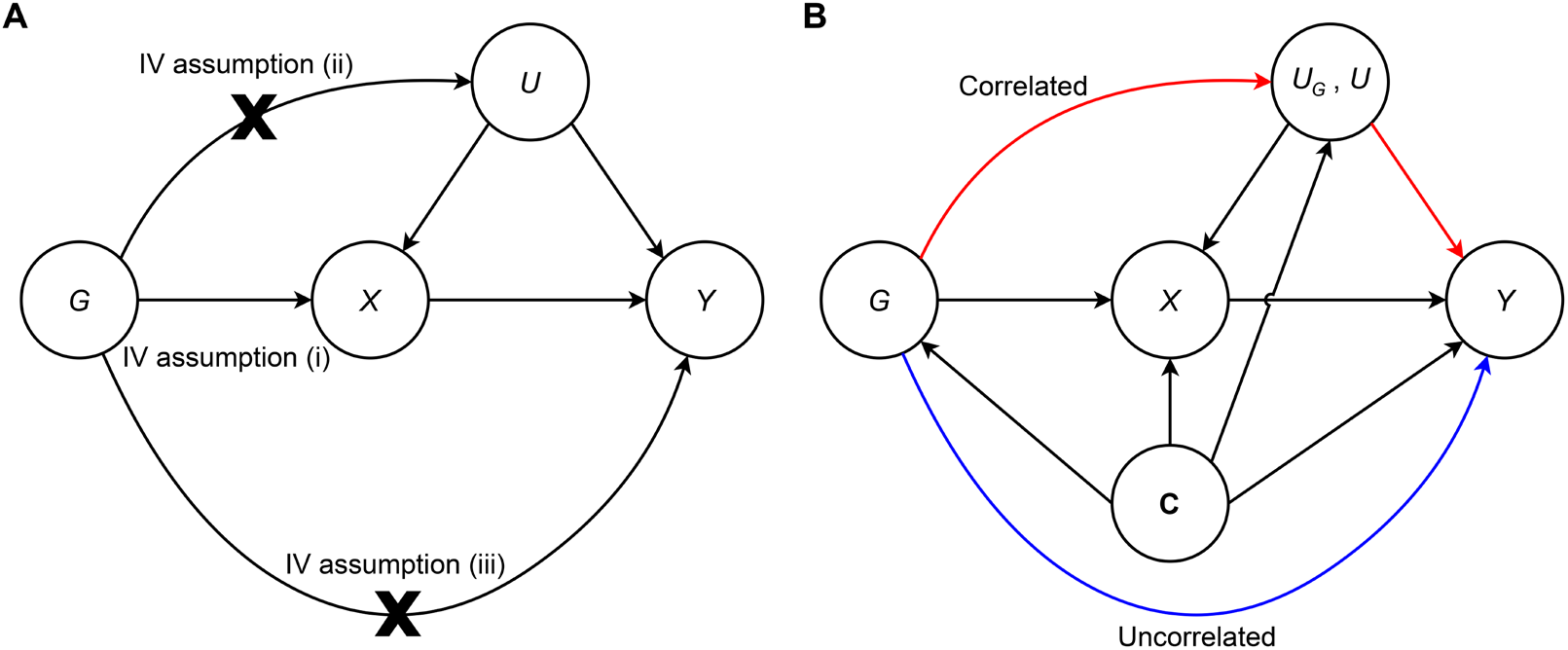
Directed acyclic graph illustrating the casual patterns assumed by traditional IV methods and MR-PROLLIM. (**A**) Causal pattern assumed by traditional IV methods. Edges marked with crosses indicate the corresponding associations should not exist. (**B**) Causal pattern that can be dealt with by MR-PROLLIM. The red and blue edges describe the correlated and uncorrelated horizontal pleiotropy, respectively. The confounder *U*_*G*_ is also a mediator that connects *G* with *X* and *Y*, while the confounder *U* is independent of *G*.

Inferring the causal effects on dichotomous outcomes is quite common in MR studies. Typically, a genome-wide association study (GWAS) uses a logistic regression to derive 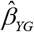. The MR ratio estimate 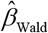 is thus widely interpreted as the causal logarithmic odds ratio (OR). However, such an interpretation is problematic because, in general, the ratio estimator is consistent for neither the conditional logarithmic OR defined by a logistic model nor the population-averaged one given by an ideal RCT (4). The resultant biases can be obvious, especially when the exposure is binary or the outcome is not rare (e.g., prevalence > 1%; see examples in **Supplementary Text S1** and **Fig. S1**). The underlying logic is mainly shaped by the non-collapsibility of OR, which means a marginal OR is not equal to the conditional one even in the absence of confounding effects (i.e., *X* is independent of *U*) (5). Besides, control variables, such as age and sex, are commonly used in MR analyses (inherited from GWAS). These variables bring extra difficulty in interpreting the MR results, as the value of OR can change obviously if other explanatory variables are controlled for (**Fig. S1A**). Fortunately, the ratio estimator is asymptotically correct on causal signs (i.e., sign consistency) when basic IV assumptions are satisfied. It can be used to test the existence and sign of a causal effect (6). However, obtaining a precise point estimate is also important, as it is the effect size that largely determines the value of a statistically significant finding.

Impeded by the non-collapsibility, consistent estimators of the conditional OR defined by a regular logistic model are difficult to construct in the IV estimation system (7). In contrast, another frequently used effect measure, the risk ratio (RR), is collapsible, easier to interpret, and can be consistently estimated using IVs. The idea of RR has been adopted in Poisson and Cox regressions, as well as the multiplicative structural mean model (8). Like a logistic model, a log-linear model can be defined as follows to incorporate RR:

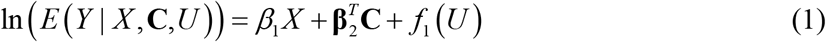

where *β*_1_ is interpreted as the logarithmic RR, and *f*_1_ is an arbitrary function that ensures *E*(*Y* | *X*, **C**,*U*) lies between 0 and 1.

Another challenge to valid MR analyses is the horizontal pleiotropy, which means an SNP may affect the outcome through pathways not mediated by the exposure (9). SNPs with horizontal pleiotropy violate the IV assumption (ii) or (iii) and will lead to biased effect estimators and even inverted casual signs (10). Some statistical methods, such as random-effects inverse-variance weighted (IVW) regression (11), MR-Egger regression (12), MR-PRESSO (10), MR-RAPS (13), CAUSE (14), and MRAID (15), have been proposed to deal with potential horizontal pleiotropy. Despite various model assumptions and statistical performances of these methods, none of them can be used to overcome the inconsistency brought by binary outcomes. Besides, these previous methods are all initially designed for summary statistic-based analyses, and many of them cannot be easily applied to one-sample MRs because of (i) failure to account for sample overlap and/or (ii) requiring a large number of SNPs (e.g., > 100,000) to be implemented. Though not as common as two-sample MRs, the one-sample design, which guarantees the homogeneity of populations (an issue needing consideration when two samples are obtained) and enables deeper investigations (e.g., sex-specific and region-specific analyses), can be seen not rarely in recent MR applications (16-18).

In this study, we propose MR-PROLLIM (Mendelian Randomization with Pleiotropy-RObust Log-LInear Model), which adopts RR as the effect measure and accounts for potential horizontal pleiotropy simultaneously. MR-PROLLIM requires individual-level data mainly measured in one sample. When the exposure is continuous, the data can be alternatively obtained from two independent samples. MR-PROLLIM provides two novel methods, respectively named random-effects and fixed-effects MR-PROLLIM, and incorporates several classical outlier-robust methods, including the median estimator (19), mode estimator (20), *Q* statistic-based outlier removal (9), and additive random-effects combination (21). MR-PROLLIM has the advantage of yielding effect estimates that have a clear estimation target and are robust to horizontal pleiotropy.

## Results

### Overview of MR-PROLLIM

MR-PROLLIM is designed to account for two types of horizontal pleiotropy (**Fig. 1B**). The first one is the uncorrelated pleiotropy, which means the size of pleiotropy is independent of the SNP-exposure effect. This may occur when an SNP affects *Y* through a pathway that is uncorrelated with *X* (blue edge). The second one is the correlated pleiotropy, which means the pleiotropic and SNP-exposure effects are correlated with each other and occurs when *G* affects an unmeasured confounder *U*_*G*_ (red edge). Such a correlation exists because two pathways point to *X*, thus cooperating to form the SNP-exposure effect, and one of them also determines the horizontal pleiotropy.

MR-PROLLIM is an integration of three distinct method series and adopts different principles to separate causality from the mixed effect (denoted as *m*) of pleiotropy and causality, including (i) random-effects MR-PROLLIM, which assumes correlated and uncorrelated pleiotropy to follow a flexible mixture distribution and requires multiple SNPs as samples, (ii) fixed-effects MR-PROLLIM, which treats horizontal pleiotropy as a fixed effect and estimates the causal and pleiotropic parameters simultaneously for each SNP under certain assumptions, and (iii) extensions of classical MR methods to log-linear models, which rely on initial estimations assuming no horizontal pleiotropy (see the schematic in **Fig. 2**). Among them, random-effects MR-PROLLIM models the SNP selection procedure and can thus incorporate more IVs to increase statistical power while reducing the selection bias. As the exposure of interest may be a continuous or categorical variable, all these methods currently have two subforms (one for continuous exposures and another for binary exposures). The difference lies in the SNP-exposure associations, for which the linear and log-linear models are used, respectively. Note that for a binary exposure, the connection between SNP-exposure and SNP-outcome effects is more complicated than that for a continuous exposure (**Materials and Methods**).

**Fig. 2.**
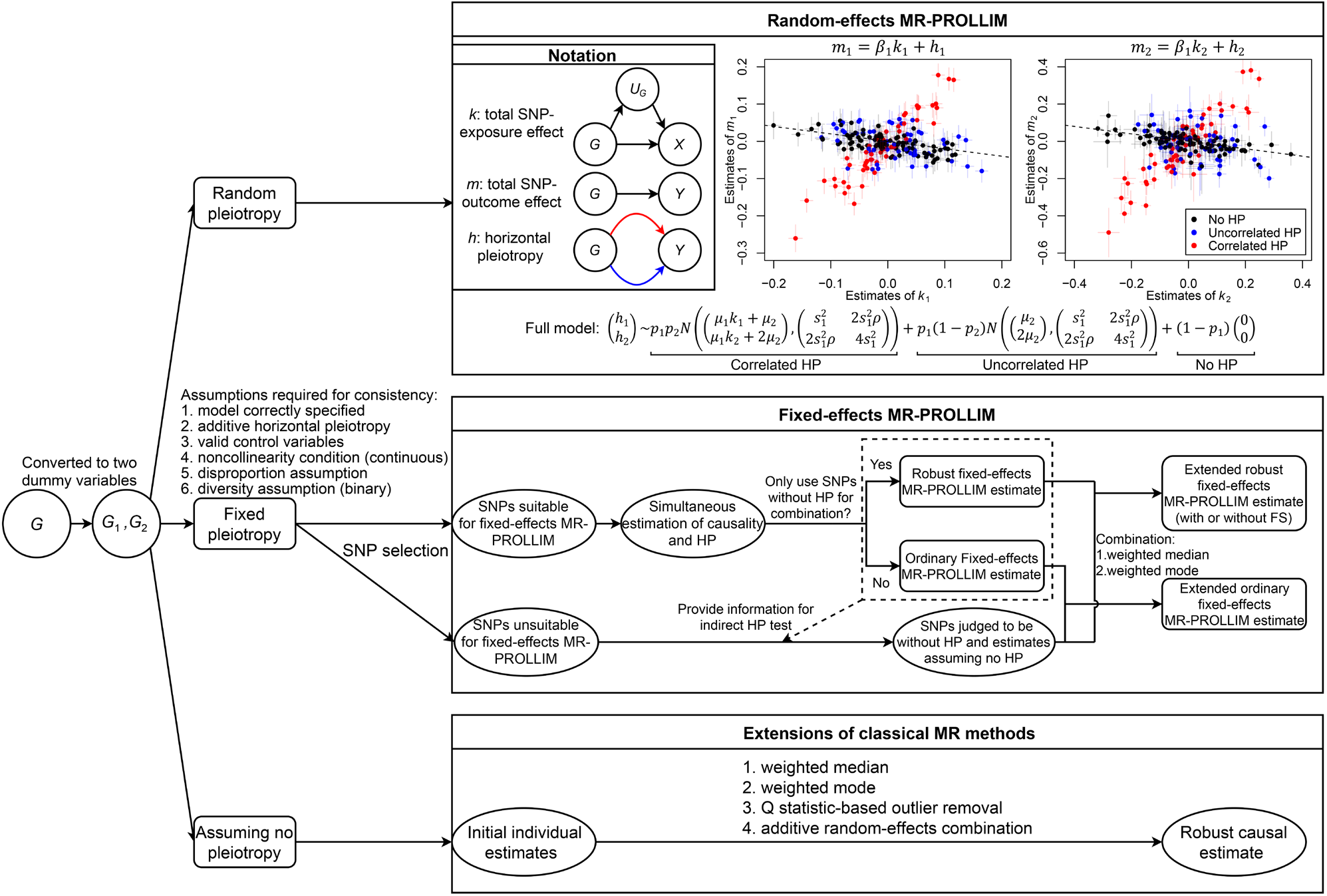
Schematic diagram of MR-PROLLIM. MR-PROLLIM allows SNPs detected in diploid organisms to exert non-additive effects on traits. Differentiated by initial assumptions on horizontal pleiotropy (HP), MR-PROLLIM consists of three distinct method series. Random-effects MR-PROLLIM assumes HP to follow the full model, where *p*_1_ ∈ (0,1) denotes the probability that the SNP has HP, *p*_2_ ∈ (0,1) denotes the probability of correlated HP, *µ*_1_ ≠ 0 is the coefficient connecting SNP-exposure effects and the mean values of correlated HP, and 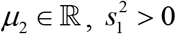, and *ρ* ∈[0,1] are additional parameters defining the bivariate normal distribution. A simulated case for a continuous exposure is presented in two scatter plots (not applicable to binary exposures). Crosses indicate 95% confidence intervals. Random-effects MR-PROLLIM estimates the causal effect (the slope of the dashed line) with the key information extracted from a group of homogenous SNPs (black points). Fixed-effects MR-PROLLIM first selects suitable SNPs and derives consistent estimates of causality and HP simultaneously under the perfectly additive horizontal pleiotropy (PAHP) assumption. It then uses the first-stage causal estimate to test for HP among unsuitable SNPs (second-stage indirect HP tests). Suitable SNPs, combined with unsuitable SNPs inferred to be without HP, are used to calculate the final estimate. The PAHP assumption can be relaxed by removing inferred pleiotropic SNPs in the first stage, which corresponds to the robust fixed-effects MR-PROLLIM. The robust version can be further classified according to with or without first-stage simplification (abbreviated as FS; i.e., whether or not to calculate the first-stage causal effect with estimates initially assuming no HP). The third type of MR-PROLLIM is established based on several classical outlier-robust methods, which accept individual effect estimates derived under the assumption of no HP.

In our proposal, MR-PROLLIM is constructed mainly based on frequentist inference and does not assign prior distributions to some key parameters. Thus, the full model of random-effects MR-PROLLIM may sometimes degenerate to other models depending on the estimated parameters and pre-specified model selection criteria, including the reduced model (*p*_2_ = 1), Egger model (*p*_1_ = 1, *p*_2_ = 0, and *µ*_1_ = 0 ; with a changeable sign before *µ*_2_), and intercept model (*p*_1_ = 1, *p*_2_ = 0, and *µ*_1_ = 0 ; with a fixed sign before *µ*_2_). More details are provided in

## Materials and Methods

### Performance of MR-PROLLIM in simulations

To investigate the statistical performances of the random-effects and fixed-effects MR-PROLLIM algorithms, we conducted a series of Monte Carlo simulations with individual-level data by resampling from the UK Biobank (UKB), which preserved the influences of linkage disequilibrium (LD) and SNP selection (**Materials and Methods**). Extensions of classical MR methods were used for comparisons. Across all simulated cases, we assumed there was a one-way or null causal effect of *X* on *Y*.

We first consider the MR analyses are conducted in the correct direction. As is shown in **Fig. 3**, the estimator given by random-effects MR-PROLLIM generally achieved the lowest median bias and median absolute error (MedAE) and yielded a coverage probability closest to the nominal level (95%). The classical weighted mode estimator also performed well in terms of median bias and coverage, but inefficiency appeared to be a weakness of this method. When the proportion of pleiotropic SNPs was not too high (e.g., *p*_1_ = 0.3 or 0.5), some other methods, including median-based ones and random-effects MR-PROLLIM, showed a higher statistical power. Another unique superiority of random-effects MR-PROLLIM was that the median bias did not increase obviously as *p*_1_ grew from 0.3 to 0.7, while other estimators were more sensitive to the proportion of SNPs having correlated pleiotropy.

**Fig. 3.**
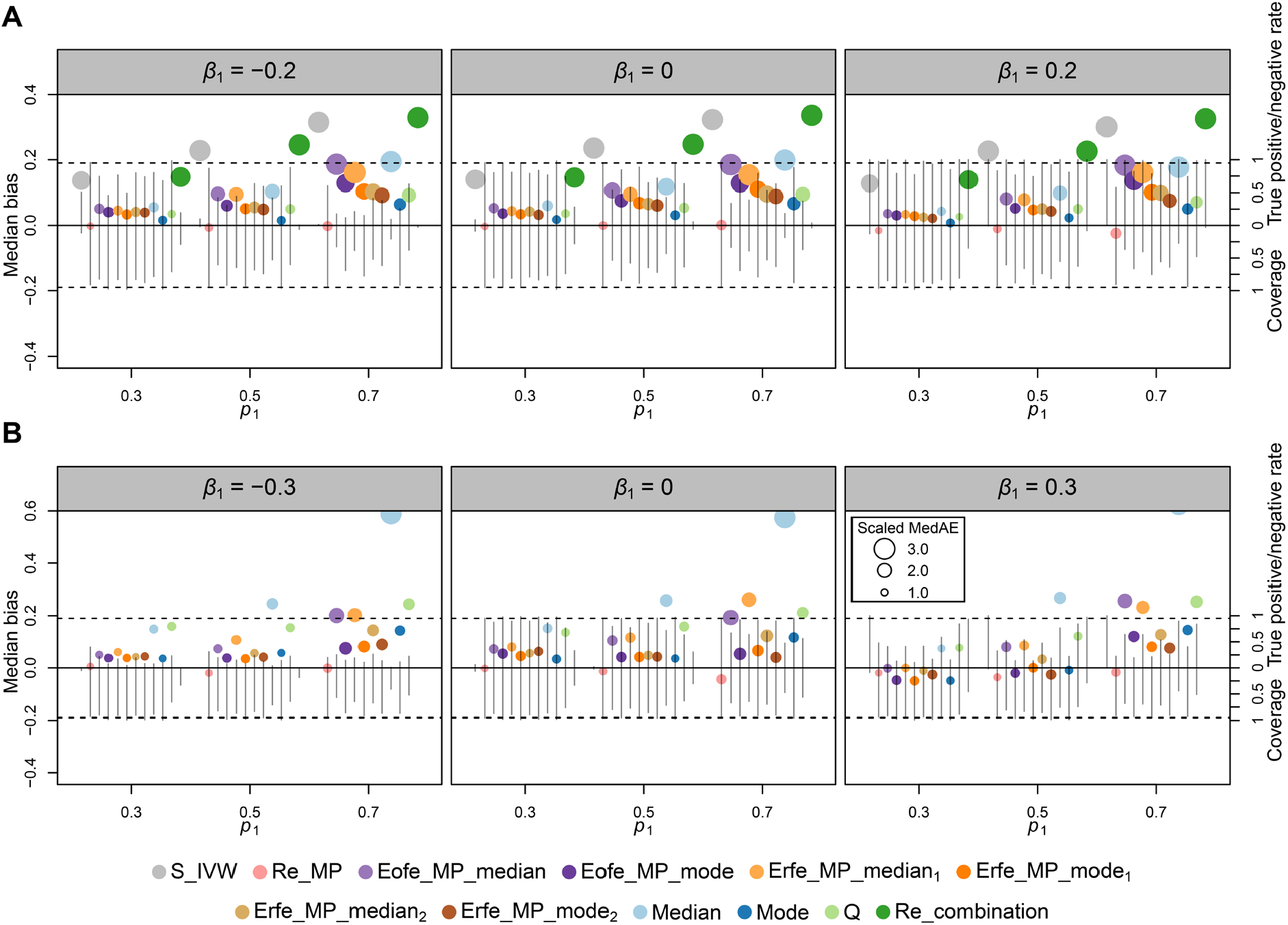
Statistical performances of different MR-PROLLIM methods in simulations with 200 effect SNPs. (**A**) Statistical performance of MR-PROLLIM for continuous exposures. (**B**) Statistical performance of MR-PROLLIM for binary exposures. The position of a colored point indicates the simulated median bias, and its size represents the scaled median absolute error (MedAE; the smallest value in each subfigure is set to 1.0, values ≥ the 75th percentile are set to 3.0, and other values are linearly transformed). The vertical line segments above the zero horizontal line indicate true positive rates for *β*_1_ ≠ 0 and true negative rates for *β*_1_ = 0. We define “true positive” as being statistically significant (unadjusted *P* < 0.05) with a correct sign of the point estimate. The vertical line segments below the zero line indicate the simulated coverage probabilities. The dashed lines mark 0.95 for the rates and coverages. We have removed outliers [point estimates ≥ (75th percentile + 3 × IQR) or ≤ (25th percentile − 3 × IQR); IQR: interquartile range] for each method to obtain more robust estimates of the evaluation indexes. For each parameter setting, 300 random simulations were conducted. The sample size was 40,000 for continuous exposures and 50,000 for binary exposures. The median biases show the same pattern across the simulations because we set the coefficient of correlated pleiotropy as a constant (*µ*_1_ = 1). Other fixed parameters included *p*_2_ = 0.5, *µ*2 = 0, and the expected heritability (**Materials and Methods**). S_IVW: simple IVW (fixed-effects IVW combination); Re_MP: random-effects MR-PROLLIM; Eofe_MP: extended ordinary fixed-effects MR-PROLLIM; Erfe_MP: extended robust fixed-effects MR-PROLLIM; Median: weighted median; Mode: weighted mode; Q: *Q* statistic-based outlier removal; Re_combination: additive random-effects combination. The subscript 1 indicates “without first-stage simplification”, and the subscript 2 indicates “with first-stage simplification”.

Regarding the fixed-effects MR-PROLLIM method series, the resultant estimators exhibited noticeable distortions in both point and interval estimations. This was principally because one of the core assumptions, the PAHP assumption (**Fig. 2** and **Materials and Methods**), was not satisfied in the simulations. Nevertheless, the extended robust fixed-effects MR-PROLLIM (Erfe_MP) median estimator (with FS; **Figs. 2** and **3**) showed an obvious reduction in median bias compared with the classical median estimator and gained a higher power than the classical mode estimator. The extended ordinary fixed-effects MR-PROLLIM (Eofe_MP) and Erfe_MP (without FS) also performed better than the classical median method in terms of bias, especially when the exposure is dichotomous.

In the above simulations, we used a bivariate normal distribution to generate random SNP-exposure effects for 200 SNPs. To obtain a more comprehensive view on MR-PROLLIM, we conducted another two series of simulations, where 1,000 SNPs were assigned with non-zero but smaller effects (Case A), and 20 SNPs showed non-zero effects with 10 of them being prominently strong (Case B; **Materials and Methods**). We observed in Case A a similar pattern of evaluation indexes as in **Fig. 3**. The random-effects MR-PROLLIM remained high-powered and robust to bias, while some other methods (e.g., median-based estimators) showed larger biases than the case of 200 SNPs (**Fig. S2**). We consider Case B may confer relative advantages on fixed-effects MR-PROLLIM, as the random-effects models usually require more SNPs to obtain precise estimates. Our simulation results supported this consideration. Notably, even without PAHP, the Erfe_MP median estimator (with FS) achieved fine performances in terms of median bias, coverage, and statistical power simultaneously when *p*_1_ was relatively small (e.g., *p*_1_ = 0.3). The random-effects MR-PROLLIM estimators showed biased coverage probabilities with insufficient SNPs (**Fig. S3**).

### Applications to inferring causality on cardiometabolic diseases

We applied MR-PROLLIM to inferring causal associations of certain living habits, physiological indexes, and blood biomarkers with cardiometabolic diseases among approximately 340 thousand UKB individuals. SNPs for MR-PROLLIM were obtained by first collecting candidates according to published GWAS summary data and then selecting based on Wald *P* values (joint test of *k*_1_ = *k*_2_ = 0 ; **Fig. 2**) calculated with individual-level data. Following GWAS conventions, we added the first ten genetic principal components, sex, and baseline age as covariates and excluded close relatives according to kinship estimates (**Materials and Methods**).

A total of 83 directed trait pairs (12 exposures and 7 outcomes) were analyzed with MR-PROLLIM. CAUSE, a Bayesian MR method, which models both correlated and uncorrelated pleiotropy (14), together with several other methods, was also adopted for summary statistic-based MR analyses. Under careful consideration of proofs from previous literature and MR results in this study, we further classified these trait pairs into three causal categories (considered causal, non-causal, and inconclusive; **Materials and Methods**). The corresponding results are partially summarized in **Table 1**, with more details provided in **Tables S1** and **S2**.

**Table 1.**
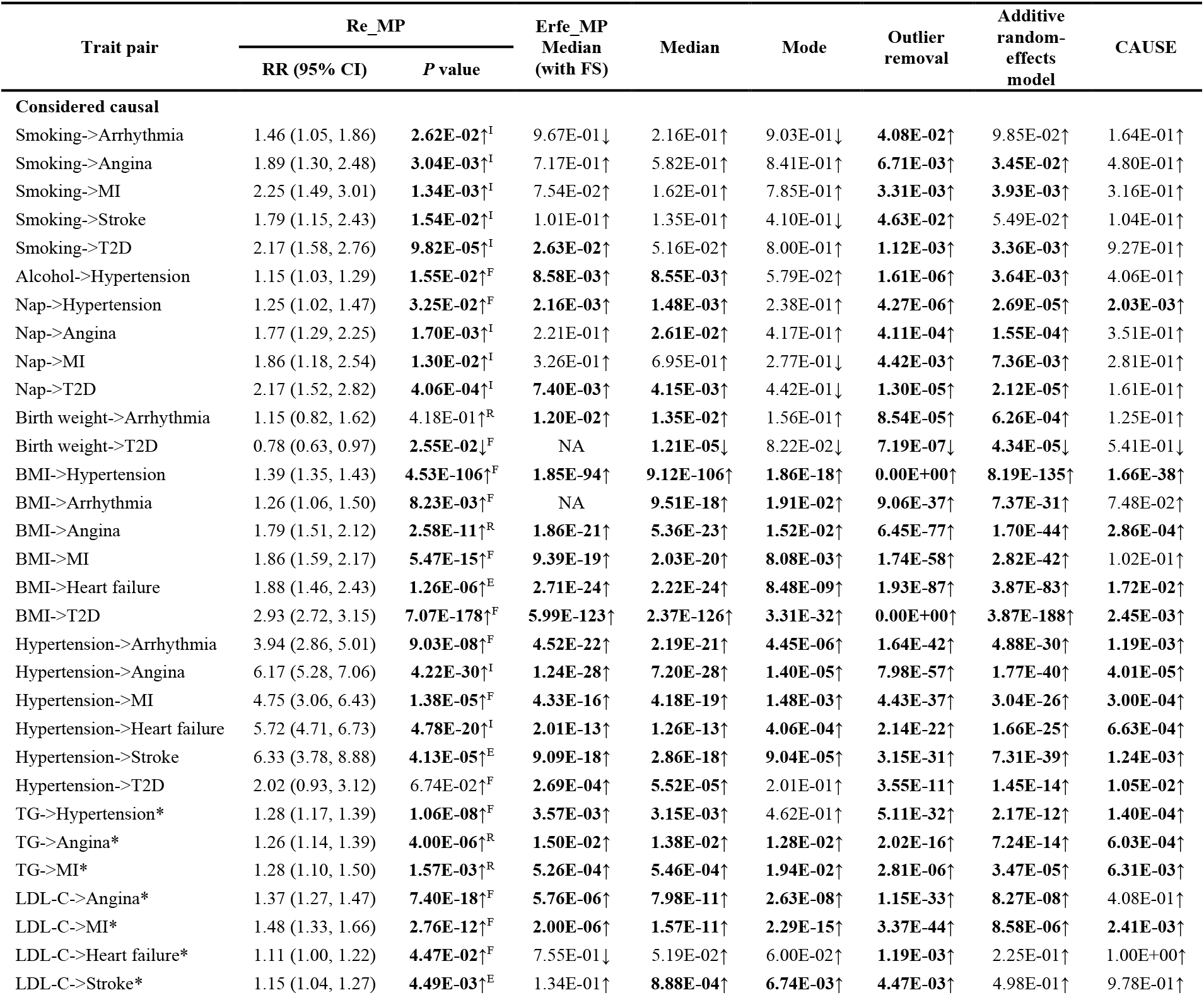
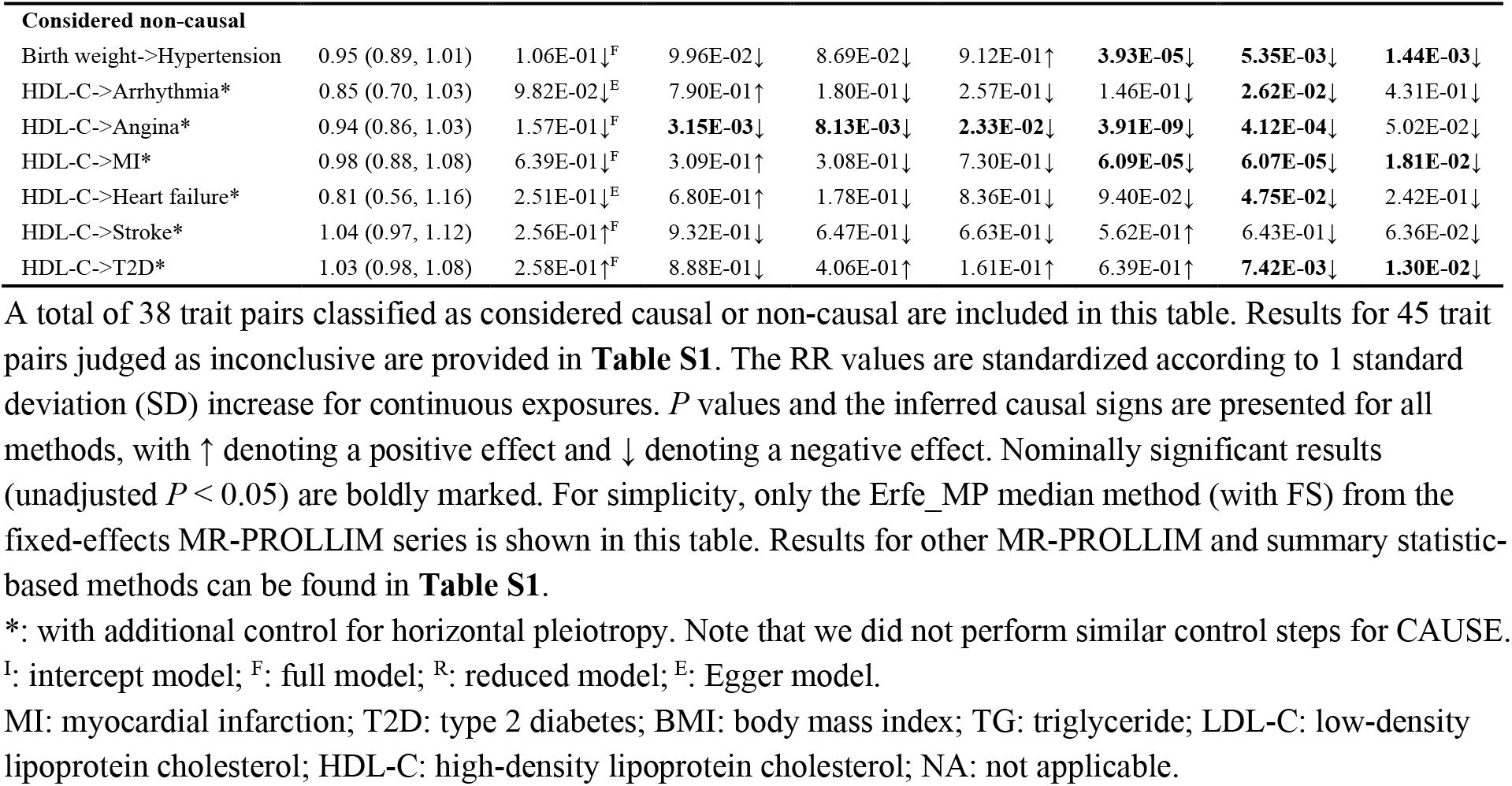
MR results of real data analyses using MR-PROLLIM and CAUSE.

We consider there are causal connections from smoking initiation, daytime napping, BMI, and high blood pressure to many of the cardiometabolic diseases, as they are generally supported by previous studies and our MR results given by certain methods (**Table S1**). The concentrations of TG, LDL-C, and HDL-C are closely related due to complicated interactions and/or shared mechanisms of lipid regulations. Previous studies have reported various loci affecting multiple lipids (22), and these loci may cause substantial biases if they are directly used as IVs. We therefore added the other two lipids as control variables when analyzing the effects of LDL-C and HDL-C. Elevated blood TG has been indicated in prior studies to influence LDL-C and HDL-C through cholesteryl ester transfer protein (CETP), which promotes cholesteryl ester and TG transfers among lipoproteins (23-25). To avoid collider bias and potential distortion of the total TG effect due to downstream blocking-up, we adopted another strategy: SNPs still showing significant effects on LDL-C or HDL-C given the TG concentration were removed from the final analysis (**Supplementary Text S2** and **Fig. S4)**.

With these additional adjustments for horizontal pleiotropy, our MR results consistently supported TG as a risk factor for hypertension, angina, and MI. However, both the methods we used and previous MR studies failed to provide accordant proofs regarding T2D. For example, one MR study reported a neutral effect of TG on T2D (26), while another one observed a likely hazardous effect (27). The true association between these two traits is unclear. But the random-effects MR-PROLLIM, which output an insignificant result (*P* = 0.11), estimated 14.8% of the SNPs might have correlated pleiotropy with 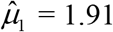. This indicates MR estimates will be biased toward larger values if no adjustment is made. In line with previous RCTs and MR studies (28), our results given by most methods identified LDL-C as a risk factor for cardiovascular diseases. However, some methods, such as fixed-effects MR-PROLLIM, weighted median (with individual-level data), and outlier removal, suggested a protective effect of LDL-C on T2D, while the random-effects MR-PROLLIM and CAUSE yielded insignificant results. Whether there exists a causal connection between these two traits is still controversial (**Table S1**) (14). HDL-C was traditionally regarded as a protective factor for a wide range of cardiometabolic diseases, yet the concept has been challenged recently due to the null effects indicated by RCTs of HDL-C-increasing drugs and MR analyses using carefully selected SNPs (29-31). We therefore classified most of the HDL-C-associated trait pairs into the considered non-causal group. If controlling for the other two lipids was not implemented, we found the majority of methods, including random-effects MR-PROLLIM, the weighted mode, and CAUSE, would produce significantly negative effect estimates for angina or MI (**Table S2**). These results indicated that there might be severe correlated pleiotropy or the distribution of pleiotropy might deviate substantially from the model assumptions. Nevertheless, with such adjustments, random-effects MR-PROLLIM obtained insignificant results, while the other methods still suggested protective effects. We further classified the SNPs according to the estimates of random-effects MR-PROLLIM (**Fig. S5**). The resultant classification plots identified a certain number of likely pleiotropic SNPs that may bias the MR results to negative values. Although we classified most HDL-C-related trait pairs as non-causal, in accordance with previous pharmacological and/or genetic evidence, we do not negate the potential role of HDL in these diseases, as HDL consists of various lipidic and proteinaceous components, and HDL-C may be a non-functional measure of HDL (32).

Previous meta-analyses of cohort studies have identified low birth weight as a risk factor for adulthood high blood pressure, coronary heart disease, and T2D (33-35). However, a recent genetic study by Warrington *et al*. found maternal birth weight-lowering SNPs (including maternal blood pressure-increasing SNPs, which might lead to pregnancy with hypertension and thus a lower birth weight) did not increase offspring blood pressure when offspring SNPs were adjusted for (36). This phenomenon indicated that the significant MR results using offspring SNPs might be substantially attributable to the effects of inherited hypertension predisposition SNPs. We thus consider there is no causality between birth weight and adulthood blood pressure. This concept is also supported by another MR study (37) and some of the methods we used, including both random-effects and fixed-effects MR-PROLLIM. For other outcomes, our results did not provide consistent evidence supporting a causal link between birth weight and MI, heart failure, or stroke (**Table S1**). Interestingly, we did find, with most methods, a protective effect of lower birth weight on arrhythmia but a hazardous effect on T2D, in line with previous MR studies (38, 39). The opposite effects reflect the complexity of long-term influences of birth weight and call for a circumspect pregnancy management to promote normal-weight newborns, thus balancing the disease risks.

Besides, we also investigated the potential effects of three widely consumed beverages (coffee, tea, and alcoholic drinks) on the cardiometabolic diseases. Previous observational studies have found moderate consumptions of coffee and tea are both associated with lower risks of many diseases, with heavy consumers still showing decreased or similar risks compared to non-consumers (40, 41). Alcohol drinkers were also reported to be less susceptible to coronary heart diseases than non-drinkers in a previous meta-analysis (42). However, except for certain trait pairs, neither MR-PROLLIM nor the summary statistic-based methods yielded consistent proofs in support of such protective effects. In contrast, our MR results suggested a hazardous effect of coffee consumption on T2D (**Table S1**). Whether it corresponds to causality requires further confirmation, as discordant MR results were reported previously (43, 44). Further research may pay more attention to different coffee and tea types, as the composition of bioactive substances may differ across types, thus leading to inconsistent effects (44). In regard to alcohol consumption, there is emerging evidence (including our results) sustaining its risk-increasing effect on hypertension (45, 46). We therefore classified this trait pair as considered causal. Despite hypertension being a strong risk factor for MI, the causal connection between alcohol and MI is considered inconclusive due to conflicting results from MR analyses and lack of support from RCTs (47, 48).

### Performance of MR-PROLLIM in real data applications

As is shown in **Fig. 4**, among the 31 considered causal pairs, random-effects MR-PROLLIM yielded nominally significant results for 93.5% of them, while the Erfe_MP median (with FS), weighted median, and weighted mode methods identified 72.4%, 77.4%, and 51.6%, respectively. This pattern remained robust when we additionally included the inconclusive pairs. The higher statistical power of random-effects MR-PROLLIM is within anticipation owing to its parametric modelling, compatibility with more SNPs, and usage of intercept models. However, the intercept model and Egger model do not allow correlated pleiotropy. We thus conducted *post hoc* diagnoses for trait pairs with these two models. We found 54.8% of the degenerations to an intercept or Egger model were attributable to lack of significant heterogeneity among individual effect estimators, which suggested the SNPs might have no pleiotropic effects. Also, sensitivity analyses did not detect substantial changes in results that seemed to be affected by correlated pleiotropy or outlier SNPs (**Supplementary Text S3, Figs. S6** and **S7**, and **Table S3**).

**Fig. 4.**
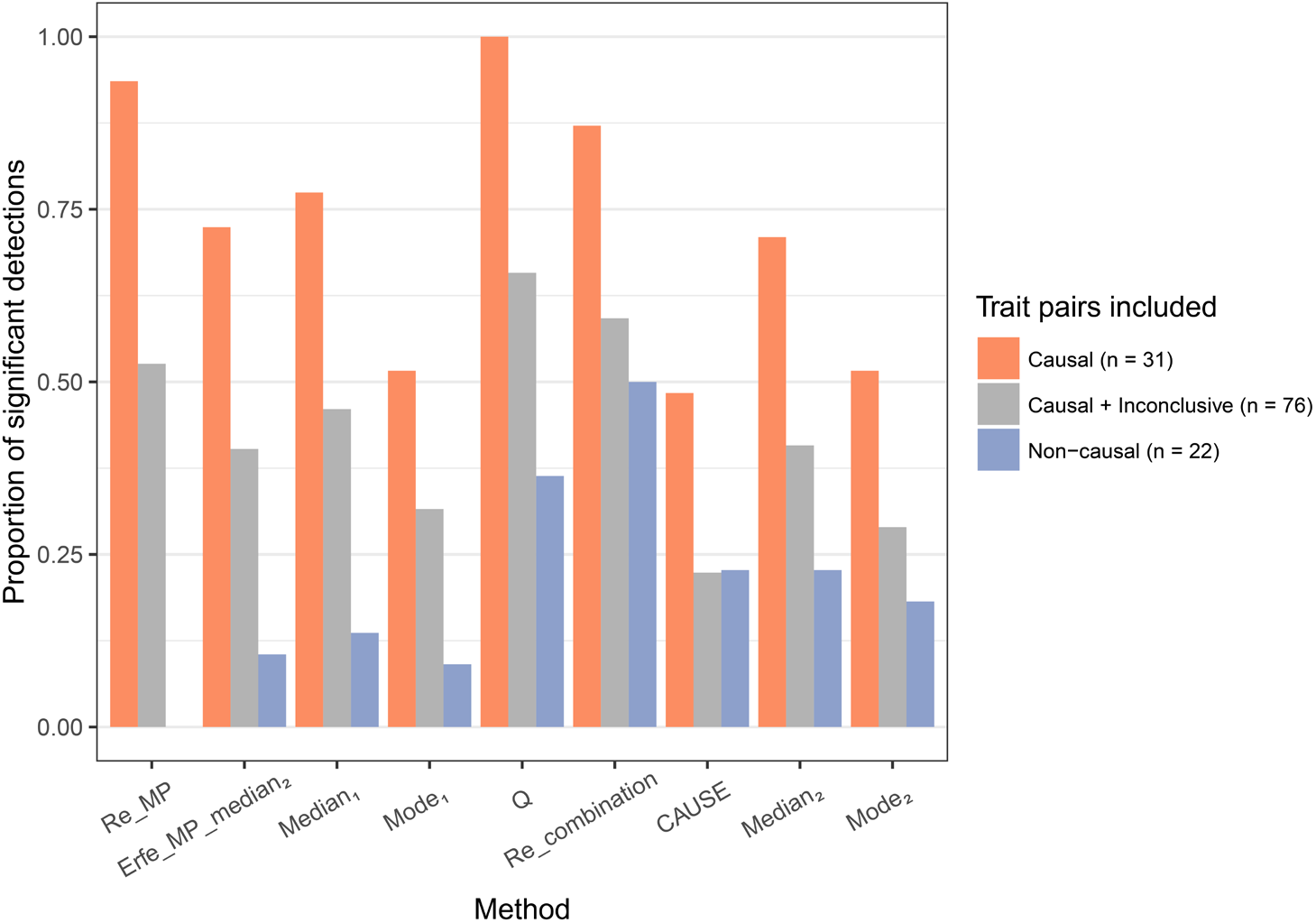
Proportions of significant detections by methods and included trait pairs. Trait pairs with birth weight as the outcome are also included in the non-causal group (see **Reverse MR analyses with MR-PROLLIM**). The classical weighted median and mode methods using individual-level data (subscript 1) and summary data (subscript 2) exhibit similar proportions of false-positive detections if controlling for the other two lipids is not conducted for HDL-C (**Table S2**).

Regarding the considered non-causal trait pairs with HDL-C or birth weight as the exposure, only random-effects MR-PROLLIM produced a throughout insignificant detection. The Erfe_MP median (with FS), weighted median, and weighted mode methods all detected one false-positive signal, while other methods detected more signals. Especially, the simple IVW and additive random-effects combination seemed severely misleading when used for HDL-C (**Table S1**).

According to random-effects MR-PROLLIM results, we further classified the 83 trait pairs into two subgroups [i.e., statistically significant or insignificant; false discovery rate (FDR)-adjusted *P* threshold = 0.1]. Among the 39 statistically significant trait pairs, the simple IVW was estimated to show relative biases ranging from -103.7% to 178.0% (median absolute value, 22.3%). And among the 44 statistically insignificant trait pairs, the simple IVW had a nominally significant detection rate of 45.5% (potential false-positive detections; unadjusted *P* < 0.05; **Tables S2** and **S4**). As an additional function, the reduced model of random-effects MR-PROLLIM can be used to test the existence of correlated pleiotropy (i.e., whether *µ*_1_ equals 0). A total of 22 (26.5%) trait pairs were inferred to have nominally significant correlated pleiotropy (**Table S2**). Among them, the simple IVW showed larger relative biases (median absolute value, 29.2% vs. 17.9%; *P* = 0.03) and a higher nominally significant detection rate (81.8% vs. 33.3%; *P* = 0.01) compared to those with insignificant or null correlated pleiotropy (**Table S4**). These results suggest that it is important to control for pleiotropy in MR analyses and that 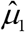 given by the reduced model can be taken as an indicator of correlated pleiotropy, which is known to be bias-introducing. But note that the correlated pleiotropy test may produce false-negative detections.

It is worth mentioning that fixed-effects MR-PROLLIM can yield a series of different estimators (**Fig. 2**). We mainly focus on the Erfe_MP median estimator (with FS) because it performed at an intermediate level between the classical median and mode estimators in both simulations and real data applications (**Fig. 3** and **Table S5**).

### Comparison of MR-PROLLIM with the MR ratio method for binary exposures

As we have shown in **Supplementary Text S1** and **Fig. S1C**, when the exposure is binary and the causality exists, the MR ratio estimate can deviate obviously from the conditional and population-averaged logarithmic ORs and appears quite small. Although we cannot know the real effect sizes, we can partially examine this through comparing the simple IVW results given by MR-PROLLIM and the ratio method among UKB trait pairs with outcomes that are not too common. The comparison results are provided in **Table S6**. We found the ratio estimates were, on average, 0.265 (range, 0.045 to 0.419) times the MR-PROLLIM estimates. If we wanted to select nominally significant trait pairs with RR (or OR) > 2 or < 0.5 for further investigations, MR-PROLLIM would identify 8 (28.6%) pairs, while the ratio method would identify 0 (even hypertension, a well-established strong risk factor for cardiovascular diseases, would be missed). These empirical results indicate that the MR ratio estimate cannot be directly interpreted as the causal logarithmic OR for the occurrence of the binary exposure. Otherwise, it may mislead the importance assessment of a causal effect.

Regarding the interpretation of the ratio estimate for a binary exposure, some researchers have suggested only treating it as a statistic for testing whether there is causality or, under further assumptions, interpreting it as the logarithmic multiplicative change in the odds of the outcome per 2.72-fold multiplicative increase in the odds of the exposure (49). But in general, this interpretation is also problematic because it cannot be easily extended to other circumstances. For example, if policy makers are interested in the effect of increasing the exposure prevalence to 100% (or close to 100%), they need to multiply the ratio estimate by positive infinity (or a very large number), which will result in an irrational effect size. On the contrary, the results provided by MR-PROLLIM do not suffer from this problem.

### Reverse MR analyses with MR-PROLLIM

Avoiding false-positive results induced by reverse causality is another challenge for MR analysis. The difficulty again arises from horizontal pleiotropy. As is shown in **Fig. 5A**, three sets of SNPs will be selected if the sample size is sufficiently large and the causality exists. SNPs from set 3 are valid IVs, but those from sets 1 and 2 are SNPs that exhibit correlated pleiotropy, thus invalidating a wide range of InSIDE assumption-dependent MR methods, such as the MR-Egger regression (11). When there is no effect of *X* on *Y*, a proportion of SNPs from set 2 may also induce correlated pleiotropy bias if no adjustment is made (**Fig. 5B**).

**Fig. 5.**
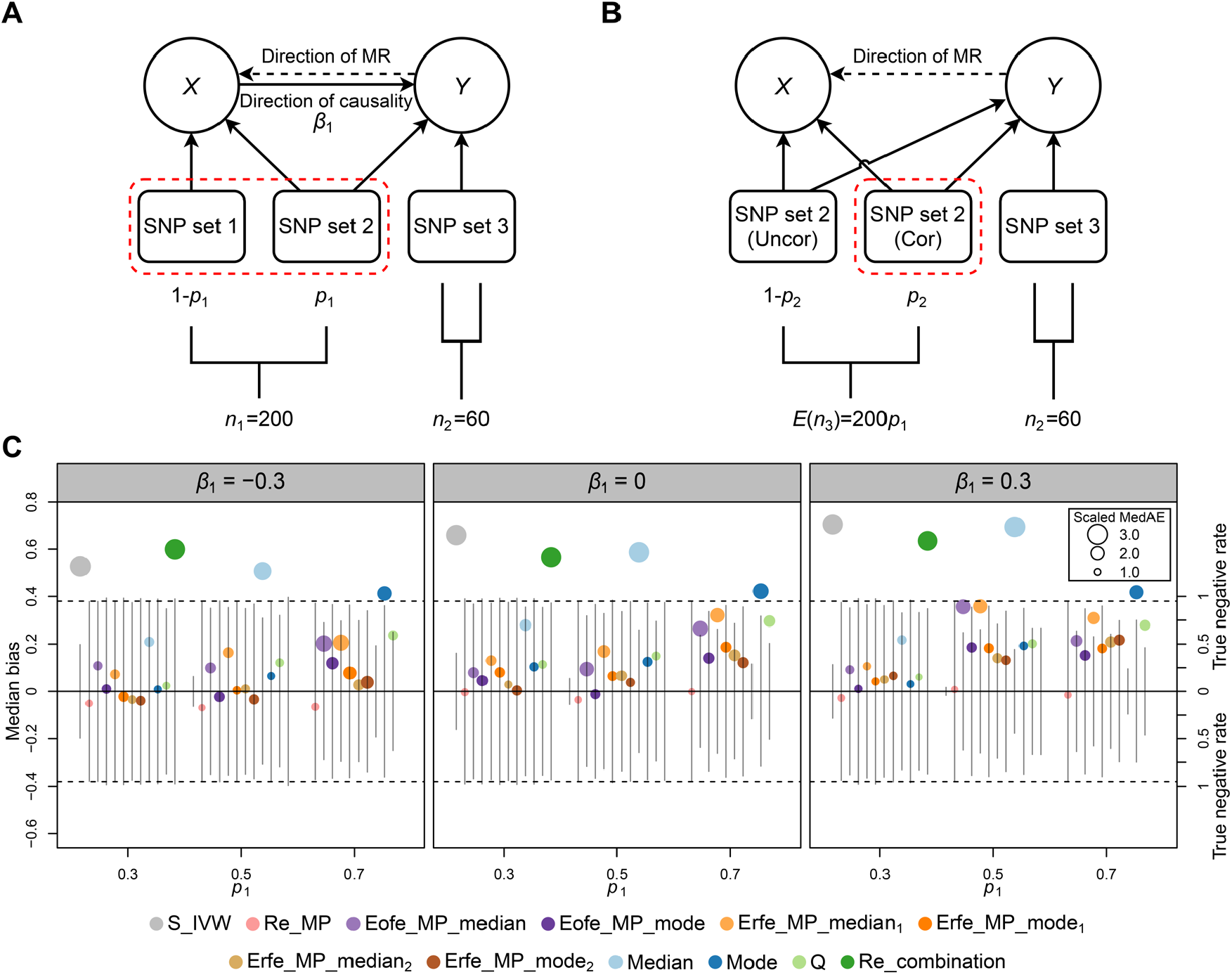
Directed acyclic graphs and simulation results of reverse MR analyses. (**A**) Directed acyclic graph illustrating involved SNPs if causality exists. (**B**) Directed acyclic graph illustrating involved SNPs if there is no causality. The red dashed curves circle the SNPs that are known to show correlated horizontal pleiotropy. The parameters *n*_1_, *n*_2_, and *n*_3_ denote the corresponding SNP numbers we set for simulations. According to which type of pleiotropy the SNPs have, the SNP set 2 in (**B**) can be further classified, with “Uncor” meaning “uncorrelated” and “Cor” meaning “correlated”. (**C**) Performances of different MR-PROLLIM methods in simulated reverse MR analyses. The position and size of a colored point indicate the median bias and scaled MedAE, respectively. Both the vertical line segments above and below the horizontal zero line mark the simulated true negative rates. See the caption of **Fig. 3** for more details. We implemented a similar procedure as for **Fig. 3** to generate individual-level data (sample size = 100,000) with 200 *X*-associated SNPs and additional 60 valid instrumental SNPs for *Y*. The parameter *p*_2_ was fixed to 0.5, and 300 replicates were conducted for each parameter setting.

To study the robustness of MR-PROLLIM in dealing with reverse causality, we performed simulations by running reverse MR analyses where both the exposure and outcome were binary variables. As is shown in **Fig. 5C**, only random-effects MR-PROLLIM yielded true negative rates conforming well to the nominal level (i.e., 0.95) across all cases we simulated. Note that a reverse MR will lead to a certain deviation from the full model described in **Fig. 2**, but it did not seem to exert too much impact. Except for random-effects MR-PROLLIM, the series of fixed-effects MR-PROLLIM estimators generally showed lower median biases than other methods. The Erfe_MP median estimator (with FS), as expected, obtained a clear improvement in true negative rate compared with the classical median estimator, but it did not reach the level achieved by the classical mode method.

Some trait pairs analyzed in this study may have bidirectional causality. However, it is chronologically implausible for an adulthood condition to have an effect on birth weight. We therefore performed real data MR analyses with birth weight as the outcome to further examine the robustness of different methods. The results are provided in **Table S7**. When all considered non-causal trait pairs were taken into account, the random-effects MR-PROLLIM, classical mode (within MR-PROLLIM framework), and Erfe_MP median (with FS) methods achieved the top three performances (**Fig. 4**).

## Discussion

We have introduced MR-PROLLIM, a collection of three different pleiotropy-robust method series designed for binary outcomes. One novel method, namely the random-effects MR-PROLLIM, which models both correlated and uncorrelated pleiotropy, exhibits superiorities in bias, statistical power, and control of false-positive detections over its competitors according to the results from simulations and real data applications. The core model of random-effects MR-PROLLIM is analogous to that of CAUSE, a summary statistic-based method that is rooted in double linear models (14). Although these two methods have different applicable scopes, we found CAUSE seemed to have a statistical power comparable with the weighted mode estimator, while our method showed a higher power. Another analog is MRAID, which parameterizes pleiotropy similarly, allows correlated SNPs, and uses summary-level data to construct maximum likelihood-based estimators (15). However, like CAUSE, this method assigns an informative prior to the probability of correlated pleiotropy. Parameters defining this prior distribution may influence the final results to a relatively large extent in some cases. Besides, MRAID currently does not account for sample overlap and is therefore not as suitable as CAUSE for our case, where only UKB GWAS data were used.

The second novel method we proposed, namely the Erfe_MP median estimator (with FS), reaches a new balance point in dealing with the trade-off between bias and statistical power. It seems to be less biased than the classical median estimator and higher-powered than the mode estimator, thus adding to the method library as a new alternative for robust MR analyses. Also, if only a few strong SNPs are available, this method may be a better choice than random-effects models, especially when one wishes to detect small exposure effects (**Fig. S3**). A previous semi-parametric method called MR-GENIUS also handles invalid IVs within a fixed-effects framework (50). However, when used for RR estimations, this method appears sensitive to model misspecifications. It may produce estimators converging in probability to non-zero values or fail to output a result if there is no causality, no pleiotropy, but a deviation from the assumed multiplicative model (e.g., changed to a logistic model).

In addition to handling pleiotropy, MR-PROLLIM largely avoids the difficulty in result interpretations, especially for binary exposures. In practice, MR-PROLLIM may provide more useful information to health consultants about the effect of (say) changing a certain behavior, to policy makers about the population benefit of an intervention, or to other fields where interpretable and confounder-robust effect sizes are highly valued.

We also note that the two novel methods have several limitations. First, MR-PROLLIM requires individual-level data mainly measured in one sample, which raises the barrier for conducting an MR analysis. Nevertheless, such MR studies also have comparative advantages over summary statistic-based ones, including no asymptotic distortion in interval estimations caused by sample overlap (a common problem of methods assuming two independent samples), no worry about the comparability of populations, and more flexibility (e.g., add additional control variables and conduct subgroup analyses). Second, the log-linear model cannot guarantee the probabilities to be less than 1 when control variables are included, and MR-PROLLIM for binary exposures may yield positive infinities as estimates if the uncertainty is large. Similar issues also exist in other RR-targeted methods (8, 50). Notwithstanding, we have proposed several solutions to handle these abnormal cases (**Materials and Methods**), and we did not get stuck in our real data applications. Third, the Erfe_MP median method may sometimes fail to provide a result due to lack of suitable SNPs. Users may choose to fill the gap with the classical median estimate. Fourth, users will need to dichotomize the traits if they want to apply MR-PROLLIM to continuous outcomes. There are not always well-established thresholds for such dichotomizations. Fifth, like many other MR methods, MR-PROLLIM focuses on approximately independent SNPs. Further research may pay attention to incorporating the information provided by correlated SNPs within similar estimation frameworks.

## Materials and Methods

### Notations and basic settings for MR-PROLLIM

MR-PROLLIM adopts structural equations to model the underlying data-generating process and defines causal effects as the model parameters that are conditional on measured and unmeasured covariates. Let *Y* denote a binary outcome, *X* denote a continuous or binary exposure, *U* denote unmeasured confounders, and **C**, an *L*-dimensional column vector, denote a set of control variables. MR-PROLLIM uses the log-linear model, such as **Eq. 1**, and the parameter *β*_1_, which can be interpreted as logarithmic RR, to describe causality between the exposure and outcome.

Although some of the proposed methods in MR-PROLLIM support additional IV types (e.g., binary variants), MR-PROLLIM focuses on biallelic SNPs detected in diploid organisms. A generic one of these SNPs is denoted as *G* and coded with 0, 1, and 2. For many complex traits, not all SNPs exert a perfectly additive effect (**Supplementary Text S4** and **Table S8**). We therefore convert *G* into two dummy variables, *G*_1_ and *G*_2_, which is compatible with the commonly assumed additive effect and reduces the impacts of potential model misspecifications. As we will show later, MR-PROLLIM needs to run log-linear regressions of *X* and *Y* on *G*_1_, *G*_2_, and **C**. If there is no control variable, the dummy variable transformation will result in a saturated model and thus surmount an innate limitation of the log-linear regression (i.e., it cannot guarantee the predicted probability lies between 0 and 1). We also propose several ways that can handle abnormal probabilities if control variables are included (i.e., modify control variables or use the Poisson regressions; **Supplementary Text S5**). It is worth mentioning that treating *G* as a categorical variable with three levels will lead to two effect estimates, and the precision of these estimates depends largely on the sample size of the reference level. MR-PROLLIM will, by default, recode each input SNP with the reference zero denoting two copies of the major allele and mark the SNP name for distinguishment.

A complex trait can be genetically determined by multiple SNPs, and some of these SNPs may be correlated with each other due to LD. MR-PROLLIM assumes the existence of a representative SNP in each LD block across the genome. The marginal effect of a representative SNP is the mixture of all SNP effects in that LD block, and the portions that cannot be represented will be merged into the error term. In this case, we can model SNP-trait associations using only independent representative SNPs, calculate individual MR estimators, and eventually combine them together like what is done in the traditional IVW method.

According to the directed acyclic graph in **Fig. 1B**, the basic MR-PROLLIM assumptions are as follows:

i. *G* has a non-zero effect on *X*.
ii. Given **C**, *G* is independent of *U*.
iii. **C** is a set of valid control variables.

The first two are just ordinary IV assumptions. Note that *G* may be correlated with *U*_*G*_ even when **C** is held constant, but we assume the effects on *X* and *Y* mediated by *U*_*G*_ can be re-expressed with the dummy variables *G*_1_ and *G*_2_. Basic assumption (iii), combined with (ii), is required for consistent estimators of the total SNP effects on *X* and *Y*. The detailed expressions of (iii) depend on whether *X* is continuous or binary, which we will demonstrate later in this study.

### Random-effects MR-PROLLIM with a continuous exposure

Taking horizontal pleiotropy into account, we model the causality between *X* and *Y* as follows:

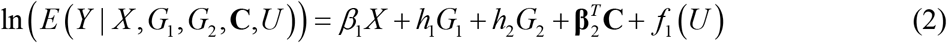

where *β*_1_ is the causal parameter, *h*_1_ and *h*_2_ denote horizontal pleiotropy, and *f*_1_(*U*) is an arbitrary function of *U*, which may include a non-zero intercept and should keep the probability less than 1. Note that if *X* is > 0, replacing *X* with ln(*X*) will lead to another meaningful interpretation of *β*_1_. That is the percentage increase in *P*(*Y* = 1) caused by a 1% increase in *X* given other covariates. For simplicity, **Eq. 2** uses a generic representative SNP to describe the pleiotropy, with other SNPs merged into *f*_1_(*U*). We assume **Eq. 2** holds for all representative SNPs and *n* randomly collected individuals.

We first consider there is a continuous exposure and suppose the following SNP-exposure association:

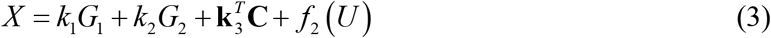

To ensure the consistency of the least squares estimators of *k*_1_ and *k*_2_, the control variables additionally need to satisfy 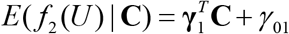, where **γ** is an arbitrary *L*-dimensional column vector, and *γ*_01_ is an arbitrary intercept. We call this the valid control variable assumption (a). Note that *f*_2_ (*U*) does not have to follow a normal distribution if the sample size is large. Substituting **Eq. 3** into **Eq. 2** and integrating out *U*, we can get:

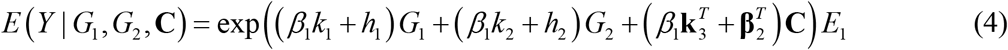

where *E*_1_ = *E*(exp(*β*_1_ *f*_2_ (*U*) + *f*_1_ (*U*)) | **C**). Similarly, we assume the valid control variable assumption (b) holds: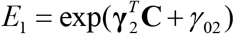. In this case, a log-linear regression of *Y* on *G*_1_, *G*_2_, and **C** with maximum likelihood estimation (MLE) can be used to derive consistent estimators of *m*_1_ = *β*_1_*k*_1_ + *h*_1_ and *m*_2_ = *β*_1_*k*_2_ + *h*_2_.

Inspired by CAUSE (14), we assume the parameters of horizontal pleiotropy, *h*_1_ and *h*_2_, to follow a mixture distribution:

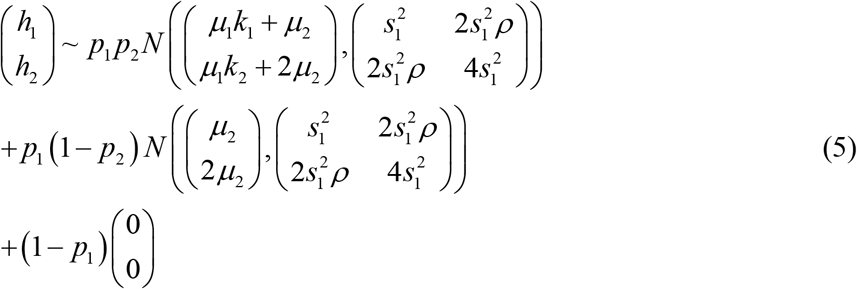

where *p*_1_ ∈ (0,1) denotes the probability that the SNP has horizontal pleiotropy, *p*_2_ ∈ (0,1) denotes the probability of correlated pleiotropy, *µ*_1_ ≠ 0 is the coefficient connecting SNP-exposure effects and the mean values of correlated pleiotropy, 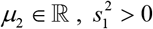, and *ρ* ∈ [0,1] are additional parameters defining the bivariate normal distribution, and (0, 0)^*T*^ represents the bivariate Dirac *δ* distribution. **Formula 5** consists of three different components, which correspond to correlated pleiotropy, uncorrelated pleiotropy, and no pleiotropy, respectively. The parameter *µ*_2_ is included in case the horizontal pleiotropy is unbalanced. As many SNPs exhibit approximately additive effects on traits, for both types of pleiotropy, we assume the variance of *h*_2_ to be four times that of *h*_1_. However, restricted by the innate limitation of the log-linear model, **Formula 5** cannot perfectly hold. Otherwise, there will be a non-zero probability that *E*(*Y* | *X, G*_1_, *G*_2_, **C**,*U*) in **Eq. 2** is > 1. Nevertheless, considering the SNP effects are typically small (i.e.,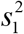 is small), we take **Formula 5** as an approximation to the real distribution of horizontal pleiotropy.

For now, we can get four consistent estimators (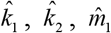, and 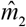). We model their joint distribution as follows:

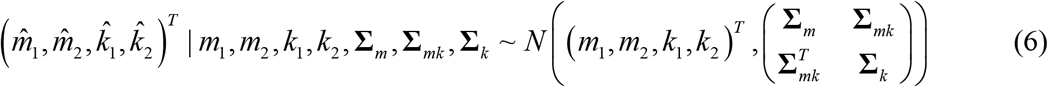

where **Σ**_*m*_, **Σ**_*mk*_, and **Σ**_*k*_ are corresponding covariance components that can be estimated using the “sandwich” method (51). We assume the covariance matrix is measured without error and can be directly replaced by its estimator. According to **Formula 6**, the following conditional distribution holds:

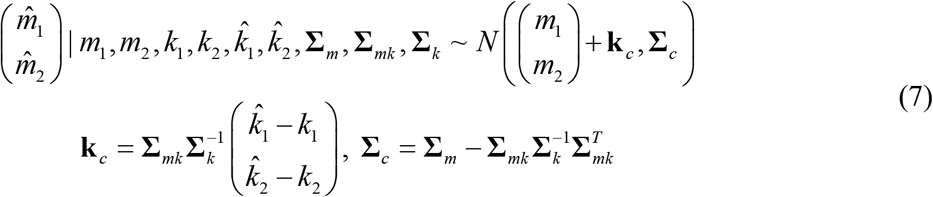

Substituting *m*_1_ = *β*_1_*k*_1_ + *h*_1_, *m*_2_ = *β k* + *h*_2_, and **Formula 5** into **7**, we can get:

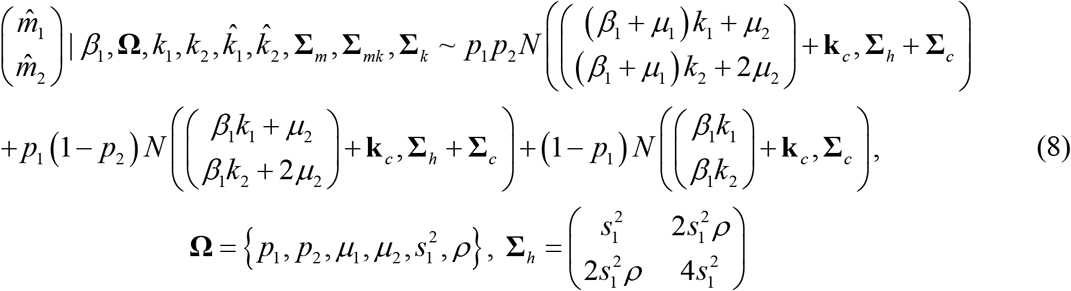

The parameters *β*_1_ and **Ω** are what MR-PROLLIM is going to estimate, but the SNP-exposure effects *k*_1_ and *k*_2_ are unknown. Some MR methods, such as the MR-Egger regression and others adopting the “first-order” variance estimates, suppose the NOME (NO Measurement Error) assumption holds (11, 52). In this case, 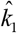 and 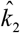 will be exactly their true values, with **k**_*c*_ being (0, 0)^*T*^ and **Σ**_**c**_ being **Σ**_**m**_. Then, the likelihood function can be written as follows:

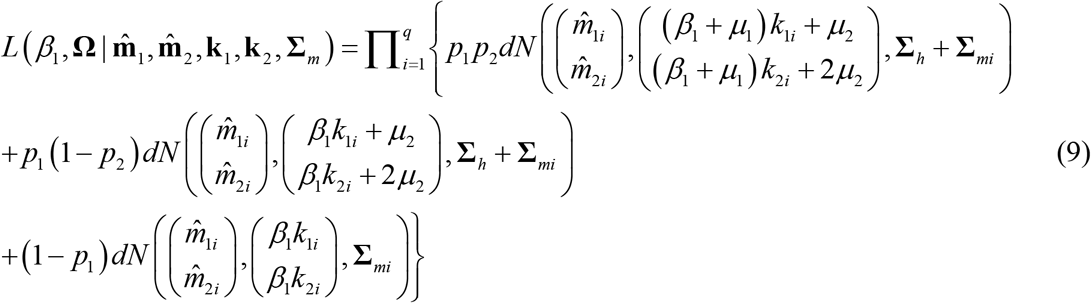

where 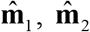, **k**_1_, **k**_2_, and **Σ**_*m*_ are lists of length *q*, containing the sample values for *q* SNPs, and *dN* denotes the density function of a bivariate normal distribution. The maximum likelihood estimators can be derived accordingly.

Ignoring the measurement errors of 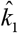 and 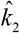 will introduce bias to the causal estimator. Through assigning a prior distribution to *k*_1_ and *k*_2_, we can integrate them out from **Formula 8** using the posterior density 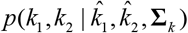 under the assumption that the influence of (*k*_1_, *k*_2_) on (**Σ**_*m*_, **Σ**_*mk*_) is negligible. If the prior distribution is flat relative to the measurement error, the posterior distribution of *k*_1_ and *k*_2_ will approximate 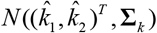, which means the results will be insensitive to the form of prior distribution under this condition. Nevertheless, we model the prior distribution as follows:

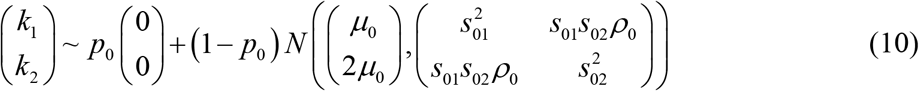

We estimate the above parameters using MLE while accounting for SNP selections according to Wald *P* values (**Supplementary Text S6**). We then adopt a sampling method to approach the likelihood function marginal on *k*_1_ and *k*_2_ and carry out MLE to derive estimators of *β*_1_ and **Ω** without the NOME assumption (**Supplementary Text S7**).

It is possible that the true values of *p*_1_ and *p*_2_ may be very close (or equal) to 0 or 1, approximately violating the assumptions of **Formula 5**. If *p*_1_ = 1 (i.e., all SNPs have horizontal pleiotropy), it can be seen from **Formula 8** that *β*_1_ is interchangeable with (*β*_1_ + *µ*_1_). If *p*_1_ = 0 (i.e., all SNPs are valid IVs), the parameters are also unidentifiable, and MLE based on **Formula 5** may yield estimates with an extremely small *ŝ*_1_. Therefore, if the initial 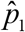 is close to 1 (e.g., > 0.999) or 0 (e.g., < 0.001; this condition is hardly met as it corresponds to a local maximum rather than the global one; **Supplementary Text S7**), or *ŝ*_1_ is too small (e.g., < 0.01 × the median value of 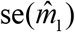; se denotes “standard error”), MR-PROLLIM will automatically turn to another two models, namely the Egger model and the intercept model. Both the models set *p*_1_ = 1, *p*_2_ = 0, and *µ*_1_ = 0, but allow an arbitrary intercept *µ*_2_. Following the MR-Egger method, which aligns the SNP-exposure effects to positive values (12), we let the signs before *µ* and 2*µ* vary with the signs of 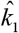 and 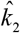 (e.g., signs before *µ*_2_ and 2*µ*_2_ are pluses if 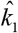 and 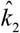 are positive, respectively) for the Egger model. On the contrary, we define the intercept model as having fixed signs before *µ*_2_ and 2*µ*_2_. These two models reflect different beliefs about the distribution of horizontal pleiotropy. MR-PROLLIM will, by default, select the one with a higher maximum likelihood. In the case that *p*_1_ equals 0, both the models can produce valid causal estimates. The intercept model may be a better choice, as its estimator generally has a smaller standard error. But in the case that *p*_1_ equals 1, selecting from them should be taken as a compromise because neither of these models allows correlated pleiotropy. We have proposed several *post hoc* analyses, which may help reduce the bias caused by correlated pleiotropy when the intercept or Egger model is selected (**Supplementary Text S3**). Furthermore, if *p*_2_ is estimated to be close to 1 or 0, MR-PROLLIM will automatically choose to fit the reduced model, which sets *p*_2_ = 1 and keeps *µ*_1_ as a free parameter. This model is compatible with both cases (i.e., *p*_2_ = 0 or 1). Note that with a finite number of SNPs, the above model selection (or degeneration) procedure will not always result in an appropriate model. See **Supplementary Text S8** and **Fig. S8** for detections of potential abnormalities in model fitting and corresponding treatments.

### Random-effects MR-PROLLIM with a binary exposure

In order to define correlated pleiotropy similarly, we first need to parameterize the association between SNPs and the binary exposure. It is quite difficult to incorporate the commonly used logistic model due to the non-collapsibility of OR. We thus propose the double log-linear model. The corresponding SNP-exposure association is as follows:

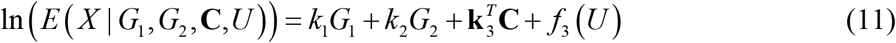

where *k*_1_, *k*_2_, and 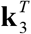 are now interpreted as logarithmic RRs rather than mean differences in **Eq. 3**. According to **Eqs. 2** and **11**, the following equations hold:

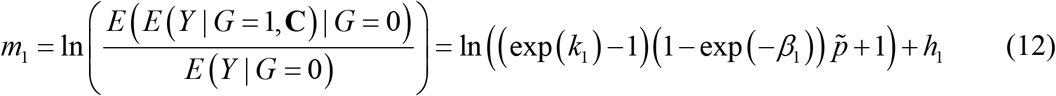

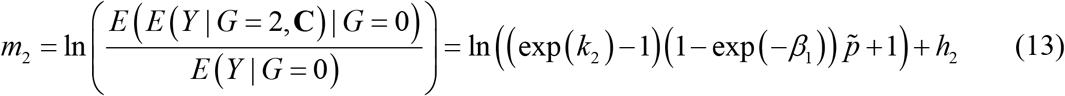

where 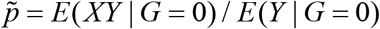. See **Supplementary Text S9** for details. The parameters *k*_1_ and *k*_2_ can be consistently estimated under the valid control variable assumption (c): 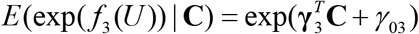. The parameter 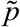 can be consistently estimated as a ratio of two moment estimates. The two iterative expectations, *E*(*E*(*Y* | *G* = 1, **C**) | *G* = 0) and *E*(*E*(*Y* | *G* = 2, **C**) | *G* = 0), can be estimated empirically, without further parametric assumptions on **C** (**Supplementary Text S10**), thus yielding consistent estimators of *m*_1_ and *m*_2_.

Similar to the procedure of dealing with a continuous exposure, MR-PROLLIM implements MLE to estimate *β*_1_ based on **Eqs. 12** and **13, Formula 5**, and the incorporation of a prior distribution for 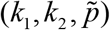 (see **Supplementary Text S6** and **S7**). Note that MR-PROLLIM first needs to estimate (1− exp(−*β*_1_)) and then transforms the corresponding point and interval estimates to those of *β*_1_. A major limitation of this model is that when the causal effect is strongly positive and the uncertainty is relatively large, the original point estimate or upper bound of the confidence interval may be > 1, thus leading to infinities after transformation. These cases can be generally avoided by coding the binary exposure inversely (i.e., changing the reference group), in which case the causal parameter *β*_1_ will shift to −*β*_1_, and the resultant estimates can be shifted backward manually. Different ways of coding *X* correspond to different model beliefs about the SNP-exposure association, as the resultant 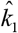 and 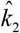 are not just added with minus signs. Nevertheless, we found in our real data analyses that recoding *X* did not seem to affect the significance judgments substantially (**Table S2**).

### Fixed-effects MR-PROLLIM with a continuous exposure

Random-effects MR-PROLLIM requires a relatively large SNP number and the existence of valid instrumental SNPs (*p*_1_ ≠ 1). However, in MR practice, there are not always enough SNPs for a reliable causal estimate under the random-effects model. We propose fixed-effects MR-PROLLIM, which, in theory, works fine with a small number of suitable SNPs. The key model is as follows:

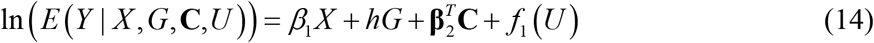

**Eq. 14** is a restricted version of **Eq. 2**. It assumes perfectly additive horizontal pleiotropy (PAHP). The causal parameter *β*_1_ and pleiotropic parameter *h* can be simultaneously identified for each SNP under further assumptions.

We first consider the case for a continuous exposure, where **Eq. 3** is supposed to hold. Under the valid control variable assumption (b), we can get from **Eqs. 3** and **14** that:

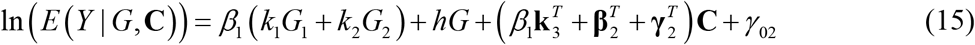

The term 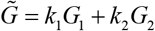 can be consistently estimated if the valid control variable assumption (a) holds. Therefore, if 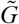 is not perfectly collinear with *G*, which we call the noncollinearity condition, both *β*_1_ and *h* can be consistently estimated by a log-linear regression of *Y* on 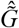, *G*, and **C**. The measurement error of 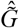 can be accounted for by integrating the estimating equations for *k*_1_ and *k*_2_ with those for the log-linear model and implementing the “sandwich” method. For each SNP that satisfies the noncollinearity condition, the causal estimate and standard error can be calculated accordingly. We then combine all individual estimators using the simple IVW method to define the ordinary fixed-effects MR-PROLLIM estimator. An outlier removal procedure based on *Q* statistics is also carried out by default to handle potential heterogeneity (see **Extensions of classical MR methods to log-linear models** below). The PAHP assumption is the key to identifiability. We therefore conducted several empirical analyses to partially examine the rationality of this assumption (**Supplementary Text S11**). In brief, we did not find a significant connection between the SNP-exposure effect pattern and the pleiotropic effect pattern among several non-causal trait pairs (**Fig. S9** and **Table S9**). These results suggest PAHP may approximately hold for most SNPs that are selected to satisfy the noncollinearity condition, as additive SNP effects appear common (**Table S8**). However, the above null connection may not be correct for all trait pairs of interest, and the ordinary fixed-effects MR-PROLLIM still suffers from the risk of model misspecifications.

Considering the properties that **Eq. 15** holds for SNPs without horizontal pleiotropy and that only if *h*_2_*k*_1_ equals *h*_1_*k*_2_ can 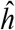 derived according to **Eq. 15** converge in probability to zero (**Supplementary Text S12**), we propose the robust fixed-effects MR-PROLLIM. Suppose **Eqs. 2** and **3**, valid control variable assumptions (a) and (b), and the noncollinearity condition hold for all SNPs, the disproportion assumption (i.e., *h*_2_*k*_1_ ≠ *h*_1_*k*_2_) additionally holds for pleiotropic SNPs, and there is at least one valid instrumental SNP. Then, a consistent causal estimator can be derived by removing SNPs that exhibit significant “pleiotropic effects” measured by 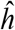 before the IVW combination. We use 0.05 as the default unadjusted *P* threshold for the SNP removal. The disproportion assumption is the key to consistency, and we do not consider it as a strong assumption (**Supplementary Text S11**).

Fixed-effects MR-PROLLIM requires an additional SNP selection for the noncollinearity condition (*P* value for 2*k*_1_ ≠ *k*_2_ should be low). The default threshold used in this study is *P* < 0.1 with a Bonferroni correction. We call the qualifiers suitable SNPs. There may be a group of significant and independent SNPs that fail to pass this selection (unsuitable SNPs), leading to information loss for the causal inference. The ordinary and robust fixed-effects MR-PROLLIM estimators are both consistent for the casual parameter under certain assumptions and can thus be used to judge whether an unsuitable SNP has horizontal pleiotropy. This is done by running a log-linear regression of *Y* on *G*_1_, *G*_2_, and **C** according to **Eq. 4** with *β*_1_ substituted by a fixed-effects MR-PROLLIM estimate. The measurement error can be accounted for in the same way mentioned previously. We adopt the Wald *P* value (joint test of *h*_1_ = *h*_2_ = 0 ; FDR-corrected *P* > 0.1 is the default) to select potential valid SNPs from the unsuitable group and calculate individual effect estimators through log-linear regressions of *Y* on 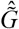 and **C** assuming no horizontal pleiotropy. These individual estimators, together with the original fixed-effects MR-PROLLIM estimator, can be combined together using the weighted median or mode method. We call the final combined estimator the extended ordinary (robust) fixed-effects MR-PROLLIM estimator (**Fig. 2**).

For the extended robust fixed-effects MR-PROLLIM, the first-stage estimator (incorporating the unsuitable SNPs is the second stage) can be alternatively calculated with individual effect estimates initially assuming no horizontal pleiotropy (i.e., the first-stage simplification; abbreviated as FS), which is expected to reduce the standard error of the first-stage estimator, thus leading to a stronger second-stage SNP filtration. Accordingly, the Erfe_MP median method (with FS), on which we mainly focus, can be viewed as a modified version of the classical weighted median method. It splits all SNPs into two groups, prunes the suitable SNPs based on the disproportion assumption and heterogeneity test, refines the unsuitable SNPs with the first-stage information, and robustly combines all information together. Regarding the control of false-positive detections, the first-stage direct pleiotropy test remains valid for suitable instrumental SNPs even if our models are erroneously specified (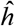 converges in probability to zero when no pleiotropy exists; **Supplementary Text S12**). It will, even under model misspecifications, remove pleiotropic SNPs with a higher probability, given that all pleiotropic SNPs violating the disproportion assumption is a zero-probability event.

### Fixed-effects MR-PROLLIM with a binary exposure

For a binary exposure, the individual estimators whose combination defines the fixed-effects MR-PROLLIM can be derived by solving **Eqs. 12** and **13** under the PAHP assumption (*h* = *h*_1_ = *h*_2_ / 2). With the elimination of *h*, we can get:

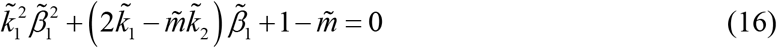

where 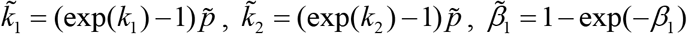, and 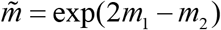. Things get complicated as **Eq. 16** (with 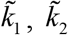, and 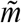 substituted by their consistent estimates) may yield two different roots if 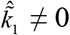, which means the causal parameter cannot be fully identified by this method with only one SNP. Nevertheless, in MR settings, there are generally a number of SNPs that exhibit different effects on the exposure (SNP-exposure effect) and outcome (horizontal pleiotropy). This diversity will lead to heterogeneity among the “wrong” roots given by **Eq. 16**, making it possible to identify the causal logarithmic RR. We call this the diversity assumption. Suppose the double log-linear model defined by **Eqs. 2** and **11**, the valid control variable assumption (c), the PAHP assumption, and *k*_1_ ≠ 0 hold for all suitable SNPs and at least two suitable SNPs satisfy the diversity assumption. Then, a consistent causal estimator can be derived by carrying out an additional root selection according to Cochran’s *Q* statistic before the IVW combination (**Supplementary Text S13**).

The robust fixed-effects MR-PROLLIM and the extended versions for binary exposures are defined similarly. When the PAHP assumption does not hold, the estimators of *h* (two possible roots) converge in probability to non-zero values if and only if (exp(*k*_1_) −1)(exp(*m*_2_) −1) ≠ (exp(*k*_2_) −1)(exp(*m*_1_) −1), which is the disproportion assumption for binary exposures (**Supplementary Text S14**). The robust fixed-effects MR-PROLLIM estimator is thus derived by removing SNPs with both roots indicating significant “horizontal pleiotropy” prior to the root selection procedure.

### Extensions of classical MR methods to log-linear models

MR-PROLLIM also incorporates the weighted median (19), weighted mode (20), *Q* statistic-based outlier removal (9), and additive random-effects combination (21), which have been proposed previously to deal with potential horizontal pleiotropy or heterogeneity among individual effect estimators. All these methods require initial estimations under the assumption of no horizontal pleiotropy.

For a continuous exposure, MR-PROLLIM runs a log-linear regression of *Y* on 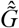 and **C** according to **Eq. 15** with *h* held as zero for each SNP to prepare the initial data. The weighted median and mode estimators are well-known for their robustness to extreme values. MR-PROLLIM follows the original studies proposing these two methods to set the weights (i.e., the inverse-variance weights) and other default parameters (19, 20). The standard errors are calculated based on parametric bootstrap procedures.

There are different versions of the *Q* statistic-based outlier removal that can be used for MR. By default, MR-PROLLIM adopts the stepwise procedure to remove the SNP with the largest *Q* decrease each time until no significant heterogeneity is detected or only two SNPs remain, of which the latter case will result in a warning message. Then, the simple IVW method will be used to combine the information of the remaining SNPs.

The last method (i.e., the additive random-effects combination) assumes the following model:

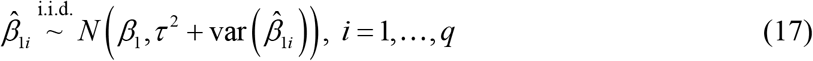

where *τ*^2^ is the between-SNP variance, measuring the impact of horizontal pleiotropy on each individual effect estimator. By default, MR-PROLLIM implements MLE to derive the estimators of *β* and *τ*^2^.

For a binary exposure, **Eqs. 12** and **13** with *h*_1_ and *h*_2_ held as zeros can be used to construct the estimator of 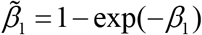. The transformed causal parameter is overidentified by two univariate equations. We thus adopt the optimal linear combination of these two estimates as follows:

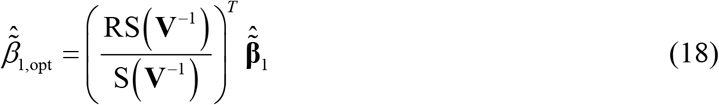

where **V** is the covariance matrix of 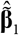, and RS(**V**^−1^) and S(**V**^−1^) denote the row sum and sum of the inverse of **V**, respectively. It is worth mentioning that there exists a generally very weak correlation among the above linear combination estimators derived using different independent SNPs. By default, MR-PROLLIM additionally estimates the covariance matrix of 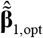 to adjust for this correlation (**Supplementary Text S15** and **Table S10**).

### SNP selection for MR-PROLLIM

MR-PROLLIM assumes a linear or log-linear association between SNPs and the exposure. It is generally required to derive Wald *P* values (joint test of *k*_1_ = *k*_2_ = 0) for all candidate SNPs prior to further steps. But it is also time-consuming to conduct a *de novo* GWAS, especially when the sample size is large. Therefore, we recommend users select a set of significant and independent SNPs as candidates first and then calculate Wald *P* values with built-in functions of MR-PROLLIM. Candidate SNPs can be obtained according to external datasets of GWAS summary statistics or by performing a *de novo* GWAS with fast algorithms, such as SAIGE (53) and fastGWA-GLMM (54), under their own model assumptions. In both cases, we recommend adopting a laxer cutoff Wald *P* value for candidates than that for MR-PROLLIM. We have described in detail how to set *P* thresholds while taking several other factors (e.g., inflated type Ι error due to multiple testing) into account in **Supplementary Text S16** and **Table S11**.

### Compatibility with two-sample MR and case-control sampling

When the exposure is continuous, it is practicable to run MR-PROLLIM with individual-level data from two independent datasets. This can be done because the required variables *k*_1_, *k*_2_, *m*_1_, and *m*_2_ for random-effects MR-PROLLIM and the regressor 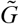 for fixed-effects MR-PROLLIM (and other methods) can be consistently estimated with the separation of datasets. The standard error and covariance matrix calculations need to be modified in two-sample MR settings.

However, when the exposure is binary, it is not feasible to incorporate dataset separation without further assumptions because MR-PROLLIM assuming the double log-linear model requires 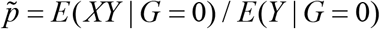 to be consistently estimated, which is done with the simultaneous measurement of *X* and *Y*.

The mean difference and RR value are sensitive to sample selections on response variables (e.g., *X* for the SNP-exposure regression and *Y* for the SNP-outcome regression). If the prevalence of the selected trait is already known, a set of weights can be calculated and assigned to all individuals to recover the cross-sectional structure, thus avoiding the bias caused by case-control sampling.

### Simulation procedure

To investigate the statistical performances of MR-PROLLIM, we conducted a series of simulations based on resampling from the UKB. Briefly, we created a genotype database of 570,600 SNPs following a realistic LD pattern, simulated control variables, and selected SNPs according to pre-specified Wald *P* thresholds and LD clumping procedures. More details are described in **Supplementary Text S17**. We also conducted a series of simplified simulations to imitate the cases where only a few strong SNPs were eventually used (i.e., Case B; **Supplementary Text S18**).

### Real data analysis

We selected 12 exposures (coffee consumption, tea consumption, smoking, alcohol consumption, daytime napping, birth weight, body mass index, blood pressure, triglyceride, low-density lipoprotein cholesterol, high-density lipoprotein cholesterol, and blood urate) and 7 cardiometabolic outcomes (hypertension, cardiac arrhythmia, angina, myocardial infarction, heart failure, stroke, and type 2 diabetes) for real data MR analyses. We conducted these analyses among UKB samples. The UKB project is a prospective cohort study that recruited half a million adults aged between 40 and 69 in the United Kingdom. Details about this study have been described previously (55, 56). The North West Multi-centre Research Ethics Committee approved the UKB study, and all participants provided written informed consent. The current study was conducted with the UKB Resource under Application Number 44430. This study was also approved by the Ethical Committee of Peking University (Beijing, China). See **Supplementary Text S19** and **Tables S12** and **S13** for details about our real data analyses.

After formal MR analyses, we further classified the trait pairs into three causal categories based on both the results in this study and proofs from prior literature. In general, a trait pair was assigned to the “considered causal” category if (i) previous RCTs and MR studies or (ii) previous MR studies and most of the pleiotropy-robust methods we used provided consistent (i.e., consistency in statistical significance and effect sign) results in support of a causal effect. A trait pair was classified as “inconclusive” if (i) there was insufficient prior knowledge from RCTs and MR studies or (ii) conflict results were presented in previous studies and/or the current study. The “considered non-causal” category included (i) most trait pairs with HDL-C as the exposure (30, 31, 57), (ii) birth weight and adulthood hypertension (36), and (iii) trait pairs with birth weight as the outcome because there was strong evidence repudiating causal links from the above exposures to the outcomes.

## Acknowledgments

This study was supported by the National Natural Science Foundation of China (grant number 82173499) and High-performance Computing Platform of Peking University. This study was conducted using the UK Biobank Resource under Application Number 44430.

## Author Contributions

M.L. and J.J. conceived and designed the methods. M.L. developed the R package with the assistance from J.J. M.L. conducted the simulations and real data analyses under the supervision of J.J. and T.H. T.H. played an important role in obtaining the UK Biobank individual-level data and interpreting the study results. All authors contributed to writing this manuscript. All authors reviewed and approved the final manuscript.

## Data and Code Availability

The UK Biobank individual-level data are available upon application to UK Biobank (https://www.ukbiobank.ac.uk). The UK Biobank GWAS summary data are publicly available at http://www.nealelab.is/uk-biobank. The GWAS summary data of alcohol consumption are publicly available at https://conservancy.umn.edu/handle/11299/201564. The MR-PROLLIM R package and R codes we used can be accessed at https://github.com/limintao-pku/MR-PROLLIM.

## References

1. B. Djulbegovic, G. H. Guyatt, Progress in evidence-based medicine: A quarter century on. Lancet 390, 415–423 (2017).

2. M. Baiocchi, J. Cheng, D. S. Small, Instrumental variable methods for causal inference. Stat. Med. 33, 2297–2340 (2014).

3. G. Davey Smith, G. Hemani, Mendelian randomization: Genetic anchors for causal inference in epidemiological studies. Hum. Mol. Genet. 23, R89–R98 (2014).

4. P. S. Clarke, F. Windmeijer, Instrumental variable estimators for binary outcomes. J. Am. Stat. Assoc. 107, 1638–1652 (2012).

5. M. Pang, J. S. Kaufman, R. W. Platt, Studying noncollapsibility of the odds ratio with marginal structural and logistic regression models. Stat. Methods Med. Res. 25, 1925–1937 (2016).

6. S. Burgess, D. S. Small, S. G. Thompson, A review of instrumental variable estimators for Mendelian randomization. Stat. Methods Med. Res. 26, 2333–2355 (2017).

7. S. Vansteelandt, J. Bowden, M. Babanezhad, E. Goetghebeur, On instrumental variables estimation of causal odds ratios. Stat. Sci. 26, 403–422 (2011).

8. M. A. Hernán, J. M. Robins, Instruments for causal inference: An epidemiologists dream? Epidemiology 17, 360–372 (2006).

9. G. Hemani, J. Bowden, G. Davey Smith, Evaluating the potential role of pleiotropy in Mendelian randomization studies. Hum. Mol. Genet. 27, R195–R208 (2018).

10. M. Verbanck, C. Chen, B. Neale, R. Do, Detection of widespread horizontal pleiotropy in causal relationships inferred from Mendelian randomization between complex traits and diseases. Nat. Genet. 50, 693–698 (2018).

11. J. Bowden et al., A framework for the investigation of pleiotropy in two-sample summary data Mendelian randomization. Stat. Med. 36, 1783–1802 (2017).

12. S. Burgess, S. G. Thompson, Interpreting findings from Mendelian randomization using the MR-Egger method. Eur. J. Epidemiol. 32, 377–389 (2017).

13. Q. Zhao, J. Wang, G. Hemani, J. Bowden, D. S. Small, Statistical inference in two-sample summary-data Mendelian randomization using robust adjusted profile score. Ann. Stat. 48, 1742–1769 (2020).

14. J. Morrison, N. Knoblauch, J. H. Marcus, M. Stephens, X. He, Mendelian randomization accounting for correlated and uncorrelated pleiotropic effects using genome-wide summary statistics. Nat. Genet. 52, 740–747 (2020).

15. Z. Yuan et al., Likelihood-based Mendelian randomization analysis with automated instrument selection and horizontal pleiotropic modeling. Sci. Adv. 8, eabl5744 (2022).

16. J. OLoughlin et al., Mendelian randomisation study of body composition and depression in people of East Asian ancestry highlights potential setting-specific causality. BMC Med. 21, 37 (2023).

17. T. Nakao et al., Mendelian randomization supports bidirectional causality between telomere length and clonal hematopoiesis of indeterminate potential. Sci. Adv. 8, eabl6579 (2022).

18. M. C. Magnus, M. C. Borges, A. Fraser, D. A. Lawlor, Identifying potential causal effects of age at menopause: A Mendelian randomization phenome-wide association study. Eur. J. Epidemiol. 37, 971–982 (2022).

19. J. Bowden, G. Davey Smith, P. C. Haycock, S. Burgess, Consistent estimation in Mendelian randomization with some invalid instruments using a weighted median estimator. Genet. Epidemiol. 40, 304–314 (2016).

20. F. P. Hartwig, G. Davey Smith, J. Bowden, Robust inference in summary data Mendelian randomization via the zero modal pleiotropy assumption. Int. J. Epidemiol. 46, 1985–1998 (2017).

21. W. Viechtbauer, Bias and efficiency of meta-analytic variance estimators in the random-effects model. J. Educ. Behav. Stat. 30, 261–293 (2005).

22. H. Tada et al., Multiple associated variants increase the heritability explained for plasma lipids and coronary artery disease. Circ.-Cardiovasc. Genet. 7, 583–587 (2014).

23. P. J. Barter et al., Cholesteryl ester transfer protein: A novel target for raising HDL and inhibiting atherosclerosis. Arterioscler. Thromb. Vasc. Biol. 23, 160–167 (2003).

24. A. D. Mooradian, Dyslipidemia in type 2 diabetes mellitus. Nat. Rev. Endocrinol. 5, 150–159 (2009).

25. J. W. Skeggs, R. E. Morton, LDL and HDL enriched in triglyceride promote abnormal cholesterol transport. J. Lipid Res. 43, 1264–1274 (2002).

26. N. M. De Silva et al., Mendelian randomization studies do not support a role for raised circulating triglyceride levels influencing type 2 diabetes, glucose levels, or insulin resistance. Diabetes 60, 1008–1018 (2011).

27. J. Liu et al., A Mendelian randomization study of metabolite profiles, fasting glucose, and type 2 diabetes. Diabetes 66, 2915–2926 (2017).

28. B. A. Ference et al., Low-density lipoproteins cause atherosclerotic cardiovascular disease. 1. Evidence from genetic, epidemiologic, and clinical studies. A consensus statement from the European Atherosclerosis Society Consensus Panel. Eur. Heart J. 38, 2459–2472 (2017).

29. G. G. Schwartz et al., Effects of dalcetrapib in patients with a recent acute coronary syndrome. N. Engl. J. Med. 367, 2089–2099 (2012).

30. B. F. Voight et al., Plasma HDL cholesterol and risk of myocardial infarction: A Mendelian randomisation study. Lancet 380, 572–580 (2012).

31. C. L. Haase, A. Tybjærg-Hansen, B. G. Nordestgaard, R. Frikke-Schmidt, HDL cholesterol and risk of type 2 diabetes: A Mendelian randomization study. Diabetes 64, 3328–3333 (2015).

32. A. von Eckardstein, B. G. Nordestgaard, A. T. Remaley, A. L. Catapano, High-density lipoprotein revisited: Biological functions and clinical relevance. Eur. Heart J. 44, 1394–1407 (2022).

33. M. Mu et al., Birth weight and subsequent blood pressure: A meta-analysis. Arch. Cardiovasc. Dis. 105, 99–113 (2012).

34. S. Wang et al., Birth weight and risk of coronary heart disease in adults: A meta-analysis of prospective cohort studies. J. Dev. Orig. Health Dis. 5, 408–419 (2014).

35. D. Mi, H. Fang, Y. Zhao, L. Zhong, Birth weight and type 2 diabetes: A meta-analysis. Exp. Ther. Med. 14, 5313–5320 (2017).

36. N. M. Warrington et al., Maternal and fetal genetic effects on birth weight and their relevance to cardio-metabolic risk factors. Nat. Genet. 51, 804–814 (2019).

37. P. Zeng, X. Zhou, Causal association between birth weight and adult diseases: Evidence from a Mendelian randomization analysis. Front. Genet. 10, 618 (2019).

38. R. L. Kember et al., Genetically determined birthweight associates with atrial fibrillation: A Mendelian randomization study. Circ.-Genom. Precis. Med. 13, e002553 (2020).

39. BIRTH-GENE (BIG) Study Working Group, Association of birth weight with type 2 diabetes and glycemic traits: A Mendelian randomization study. JAMA Netw. Open 2, e1910915 (2019).

40. M. Ding, S. N. Bhupathiraju, A. Satija, R. M. van Dam, F. B. Hu, Long-term coffee consumption and risk of cardiovascular disease: A systematic review and a dose-response meta-analysis of prospective cohort studies. Circulation 129, 643–659 (2014).

41. M. Inoue-Choi et al., Tea consumption and all-cause and cause-specific mortality in the UK Biobank: A prospective cohort study. Ann. Intern. Med. 175, 1201–1211 (2022).

42. P. E. Ronksley, S. E. Brien, B. J. Turner, K. J. Mukamal, W. A. Ghali, Association of alcohol consumption with selected cardiovascular disease outcomes: A systematic review and meta-analysis. BMJ 342, d671 (2011).

43. A. T. Nordestgaard, M. Thomsen, B. G. Nordestgaard, Coffee intake and risk of obesity, metabolic syndrome and type 2 diabetes: A Mendelian randomization study. Int. J. Epidemiol. 44, 551–565 (2015).

44. X. Wang, J. Jia, T. Huang, Coffee types and type 2 diabetes mellitus: Large-scale cross-phenotype association study and Mendelian randomization analysis. Front. Endocrinol. 13, 818831 (2022).

45. K. J. Biddinger et al., Association of habitual alcohol intake with risk of cardiovascular disease. JAMA Netw. Open 5, e223849 (2022).

46. S. van Oort, J. W. J. Beulens, A. J. van Ballegooijen, D. E. Grobbee, S. C. Larsson, Association of cardiovascular risk factors and lifestyle behaviors with hypertension: A Mendelian randomization study. Hypertension 76, 1971–1979 (2020).

47. S. C. Larsson, S. Burgess, A. M. Mason, K. Michaëlsson, Alcohol consumption and cardiovascular disease: A Mendelian randomization study. Circ.-Genom. Precis. Med. 13, e002814 (2020).

48. T. Hisamatsu et al., Alcohol consumption and subclinical and clinical coronary heart disease: A Mendelian randomization analysis. Eur. J. Prev. Cardiol. 29, 2006–2014 (2022).

49. S. Burgess, J. A. Labrecque, Mendelian randomization with a binary exposure variable: Interpretation and presentation of causal estimates. Eur. J. Epidemiol. 33, 947–952 (2018).

50. E. T. Tchetgen, B. Sun, S. Walter, The GENIUS approach to robust Mendelian randomization inference. Stat. Sci. 36, 443–464 (2021).

51. D. A. Freedman, On the so-called “Huber Sandwich Estimator” and “Robust Standard Errors”. Am. Stat. 60, 299–302 (2006).

52. J. Bowden et al., Improving the accuracy of two-sample summary-data Mendelian randomization: Moving beyond the NOME assumption. Int. J. Epidemiol. 48, 728–742 (2019).

53. W. Zhou et al., Efficiently controlling for case-control imbalance and sample relatedness in large-scale genetic association studies. Nat. Genet. 50, 1335–1341 (2018).

54. L. Jiang, Z. Zheng, H. Fang, J. Yang, A generalized linear mixed model association tool for biobank-scale data. Nat. Genet. 53, 1616–1621 (2021).

55. C. Sudlow et al., UK Biobank: An open access resource for identifying the causes of a wide range of complex diseases of middle and old age. PLoS Med. 12, e1001779 (2015).

56. C. Bycroft et al., The UK Biobank resource with deep phenotyping and genomic data. Nature 562, 203–209 (2018).

57. D. Keene, C. Price, M. J. Shun-Shin, D. P. Francis, Effect on cardiovascular risk of high density lipoprotein targeted drug treatments niacin, fibrates, and CETP inhibitors: Meta-analysis of randomised controlled trials including 17,411 patients. BMJ 349, g4379 (2014).

