## Supplementary Materials for "Mendelian randomization analysis with pleiotropy-robust log-linear model for binary outcomes"

**This PDF file includes:**

Supplementary Text S1 to S19
Figs. S1 to S9
Tables S3 to S6, S8 to S11, and S13
Supplementary References

**Other supplementary materials for this manuscript include the following:**

Tables S1, S2, S7, and S12

### Supplementary Text

#### Section S1. Examples illustrating the non-collapsibility of OR in MR analyses

We present in this section the details of the four examples (Examples 1 to 4; **Fig. S1**) that we simulated to illustrate the non-collapsibility of OR and its influence on MR results. If the outcome is rare, a logistic regression will approximate a log-linear regression, which does not suffer from the non-collapsibility. We demonstrate in Examples 2 and 3 that outcome prevalence being less than 10% (1-3) may not be a good indicator of low bias in MR settings, where the influence of covariates (measured and unmeasured) cannot be neglected.

The data-generating process for Example 1 is as follows:

$$X \sim \text{Bernoulli}(0.5), C_1 \sim N(0,1), C_2 \sim \text{Bernoulli}(0.5),$$

$$\text{logit}(P_Y) = \ln\left(\frac{P_Y}{1-P_Y}\right) = X + C_1 + C_2 - 1,$$

where $X$, $C_1$, and $C_2$ are mutually independent.

The data-generating process for Example 2 (continuous exposure) is as follows:

$$C, U_1, U_2 \overset{\text{i.i.d.}}{\sim} N(0,1),$$

$$X = 0.2G + 0.5C + 1.2U_1,$$

$$\text{logit}(P_Y) = X + 2C + 0.3U_1 + 1.5U_2 - l,$$

where $C$ is an observed control variable (e.g., sex and age; they are commonly included in GWAS), $G$ denotes an instrumental SNP with a minor allele frequency of 0.3 (following Hardy-Weinberg's equilibrium), and $l$ is used to adjust the outcome prevalence. We calculate the population-averaged logarithmic OR as:

$$\ln\left(\text{odds}\left(M(P_{Y,X})\right)/\text{odds}\left(M(P_{Y,X-d})\right)\right)/d,$$

where $\text{odds}(x) = x/(1-x)$, $M$ denotes a simple average across the sample, and $d$ is set to 0.01. This value can be viewed as the one obtained by an ideal RCT that reduces the exposure levels of all individuals in the treatment group by $d$ (4). It is considered to have significance in the field of public health.

The data-generating process for Example 3 (binary exposure) is as follows:

$$C, U_1, U_2 \overset{\text{i.i.d.}}{\sim} N(0,1),$$

$$\text{logit}(P_X) = 0.5G + 0.8C + 1.5U_1 - l_1,$$

$$\text{logit}(P_Y) = -X + 2C + 0.5U_1 + 1.5U_2 - l_2,$$

where $G$ denotes an instrumental SNP with a minor allele frequency of 0.3. Considering that if both $X$ and $Y$ are rare, an MR analysis may be underpowered. We thus set the exposure prevalence to about 30% to illustrate the biases. The MR ratio estimate is calculated using two logistic regressions. The population-averaged logarithmic OR is calculated among individuals with $X = 0$ as:

$$\ln\left(\text{odds}\left(M(P_{Y,X+1})\right)/\text{odds}\left(M(P_{Y,X})\right)\right).$$

Note that even if the SNP-exposure and exposure-outcome associations follow log-linear models (i.e., the non-collapsibility is not a problem), the causal effect cannot be expressed as a ratio of two SNP effects (see the double log-linear model in **Materials and Methods**).

The data-generating process for Example 4 is as follows:

$$U_1, U_2 \overset{\text{i.i.d.}}{\sim} N(0,0.3^2),$$

$$X = G_1 - G_2 + U_1,$$

$$\text{logit}(P_Y) = X + U_1 + U_2 + 1,$$

where $G_1$ and $G_2$ are two strong instrumental SNPs with minor allele frequencies of 0.3 and 0.5, respectively. This example is considered unrealistic because the SNP effects are typically small in MR settings. We use it to demonstrate that the non-collapsibility of OR may, in theory, lead to significant heterogeneity among individual effect estimators even if all SNPs are valid IVs.

### Section S2. Adding control variables in MR-PROLLIM

We roughly classify the control variables that may be included in MR-PROLLIM into three types based on inclusion purposes. The first type contains variables measuring population structure, such as self-reported races, inferred ancestries, and genetic principal components. Population structure is a potential confounder between SNPs and traits. It is generally recommended to include this type of variables, especially when the samples are obtained from a genetically heterogeneous population. The second type corresponds to variables that affect the exposure and/or outcome but do not affect the genotypes (e.g., age). These variables are not confounders. Adding them as covariates may reduce the standard errors of the SNP-trait effect estimators and thus contribute to a more precise MR result. The third type, which should be treated with caution, consists of additional control variables for horizontal pleiotropy.

Users may consider the third-type control variables (denoted as $C$) if there is evidence that the SNPs may act through them to affect $Y$ (i.e., horizontal pleiotropy). According to the causality connecting $X$ and $C$, we further classify these variables into three subtypes (Subtype A, B, and C; **Fig. S4**).

For Subtype A, if there is no effect of $U$ on $C$ (red edge in **Fig. S4A**) or the direction is inverted, the horizontal pleiotropy can be adjusted for by including $C$ as a control variable for both the SNP-exposure and SNP-outcome regressions. However, if the red edge does exist, such regressions will suffer from collider biases, which arise because $G$ is correlated with $U$ when $C$ is held constant. Nevertheless, there are several reasons supporting this additional control variable, even in the presence of collider biases:

i. The control variable $C$ is a downstream effector of $U$. If $C$ is included in the causal system, the direct effect of $U$ on $Y$ may be attenuated, resulting in a small bias of the final MR estimate.
ii. If the collider bias is relatively small for the SNP-exposure regression, holding $C$ constant can contribute to the removal or down-weighting of the SNPs that have small direct effects on $X$ but large effects on $C$. These SNPs are expected to cause substantial biases due to horizontal pleiotropy if no adjustment is made.
iii. Most importantly, the collider bias only influences SNPs having effects on $C$ (valid instrumental SNPs will not be affected). The collider effect behaves like a pleiotropic effect and may be smaller according to (i) and (ii), which may thus improve the pleiotropy-robust estimates given by MR-PROLLIM.

It is worth mentioning that users may alternatively choose to remove the SNPs which exhibit significant marginal effects on $C$. But this strategy may suffer from false-negative detections and loss of information due to SNP removal.

For Subtype B, even if the red edge (**Fig. S4B**) does not exist, directly controlling for $C$ will lead to collider biases, as both $U$ and $G$ have pathways pointing to $C$. An alternative strategy is to remove the SNPs that still exhibit significant effects on $C$ when $X$ is held constant. The conditional effects can be estimated based on linear regressions (continuous $C$) or logistic (or log-linear) regressions (binary $C$). If the red edge does exist, these regressions will also suffer from collider biases. If such biases are relatively small, this strategy is more likely to rule out SNPs having strong direct effects on $C$, which may result in a residual pleiotropic pattern that can be more easily handled by MR-PROLLIM.

If there is no causal connection between $X$ and $C$ (Subtype C; **Fig. S4C**), strategies for Subtype A (i.e., controlling for $C$ and SNP removal based on marginal estimates) can be adopted for this subtype as well.

With regard to blood lipids, we assume the TG concentration may causally influence LDL-C and HDL-C, and the levels of LDL-C and HDL-C are correlated with each other mainly due to shared regulatory mechanisms. When analyzing the effects of TG on diseases, we removed TG-related SNPs that showed significant conditional effects on the other two lipids (at least one of the FDR-adjusted $P$ values < 0.05). Although in the main text, we majorly discussed the results with additional adjustments for horizontal pleiotropy based on the strategies mentioned above, we found only the trait pairs with HDL-C as the exposure exhibited obvious changes (**Table S2**).

**Section S3. *Post hoc* analyses for random-effects MR-PROLLIM with the intercept or Egger model**

The intercept or Egger model will be selected if $\hat{p}_1$ is very close to 1 or 0, or $\hat{s}_1$ is too small (**Materials and Methods**). We consider three cases that may result in these models: (a) $p_1 = 0$ (i.e., no SNPs have pleiotropic effects); (b) $p_1 = 1$ and $p_2 = 0$ (i.e., all SNPs have uncorrelated pleiotropy); (c) $p_1 = 1$ and $p_2 \neq 0$ (i.e., all SNPs have pleiotropic effects, and some SNPs have correlated pleiotropy). We mainly focus on case (c) because it will introduce bias to the casual estimator derived under an intercept or Egger model. We propose a procedure of *post hoc* analyses to detect potential correlated pleiotropy and to reduce its influence (see **Fig. S6**).

Heterogeneity tests can be conducted based on $Q$ statistics within the third MR-PROLLIM framework (**Fig. 2** and **Materials and Methods**). An insignificant result indicates all SNPs may be valid IVs. In this case, users may accept the current model or turn to the intercept model for higher statistical power if the Egger model does not suggest a non-zero $\mu_2$. We do not recommend directly setting $p_1 = 0$ because the intercept and Egger models may capture small pleiotropic effects that are undetectable to heterogeneity tests. On the contrary, if significant heterogeneity is detected, we recommend using the classification plots (**Fig. S5**) for further diagnoses. Under the full model of random-effects MR-PROLLIM, there should be three groups of points: (a) points that are densely distributed around the zero horizontal line (most confidence intervals should cover the zero line), (b) points that are distributed around the blue horizontal dashed line (the variance is expected to be larger than that of black points), and (c) points that are distributed around the red dashed line (the variance is also expected to be larger, and the slope of the line should deviate from zero). If there is correlated pleiotropy, the classification plots of an intercept or Egger model are expected to show two groups of points (typically distributed in an X-shaped pattern), though the causal estimate is biased. In the absence of valid instrumental SNPs, MR-PROLLIM cannot distinguish causality and correlated pleiotropy without further assumptions. But if we have a prior belief that the majority of SNPs only have uncorrelated pleiotropy (5), we can identify the causal effect by removing the minority SNPs or fitting the weak correlation model ($p_1 = 1$, $p_2 < 0.5$, and $\mu_1$ is kept as a free parameter). Note that we recommend avoiding the weak correlation model when the SNP separation is not obvious because it may overfit the data and give misleading results. We also recommend users avoid directly conducting these two sensitivity analyses if the proportion of the minority seems too large (i.e., $p_2$ is close to 0.5), in which case professional knowledge about the SNPs may be required.

In our real data applications, we identified that 19 (45.2%) trait pairs with intercept or Egger models showed significant heterogeneity (unadjusted $P > 0.05$; **Table S1**). We then checked the classification plots and found eight trait pairs were likely to be affected by outlier SNPs or correlated pleiotropy. We further conducted sensitivity analyses for these trait pairs. The results are provided in **Table S3**. Classification plots for three trait pairs with relatively clear abnormalities are given in **Fig. S7**.

**Section S4. Detection of non-additive SNP effects**

For a given complex trait, an associated SNP may have a non-additive effect, which means the effect of two alternative alleles is not exactly twice the effect of only one allele (i.e., $k_2 \neq 2k_1$). We conducted empirical analyses among 12 traits of interest using the UKB data to investigate the proportions of such SNPs. For each trait, we first adopted linear (continuous trait) or log-linear (binary trait) regressions to collect a group of significant (Wald $P$ value < 5E-5 for alcohol and < 1E-5 for other traits) and approximately independent (LD $r^2 < 0.1$) SNPs from the pre-selected candidates (selected based on GWAS summary data; **Materials and Methods**). We next tested the null hypothesis, $k_2 = 2k_1$, for each of them according to $Z$ statistic calculated as follows:

$$Z = \frac{2\hat{k}_1 - \hat{k}_2}{\sqrt{4\text{var}(\hat{k}_1) + \text{var}(\hat{k}_2) - 4\text{cov}(\hat{k}_1, \hat{k}_2)}},$$

where the variances and covariance were directly replaced by their estimates. We defined "nominally non-additive" as having an unadjusted $P$ value of < 0.05.

If the null hypothesis is true, we will have a probability of 0.05 to obtain a false-positive detection. For each SNP, we expect the following equations hold:

$$P_{\text{positive}} = P_{\text{power}} P_{\text{non-additive}} + 0.05(1 - P_{\text{non-additive}}),$$

$$P_{\text{non-additive}} = \frac{P_{\text{positive}} - 0.05}{P_{\text{power}} - 0.05},$$

where $P_{\text{positive}}$ denotes the probability of a positive (i.e., nominally significant) detection, $P_{\text{power}}$ denotes the statistical power, and $P_{\text{non-additive}}$ is the probability that the SNP has a non-additive effect. We can assume $P_{\text{power}} = 1$ and use the empirical estimate of $P_{\text{positive}}$ to calculate $\hat{P}_{\text{non-additive}}$. Note that such an estimate is conservative as the statistical power cannot always reach a level close to 1. The corresponding results are provided in **Table S8**. In summary, we did find evidence that there were a proportion of SNPs exhibiting non-additive effects, but the proportion seemed not so high, especially for continuous traits.

**Section S5. Avoiding abnormal probabilities in log-linear regressions**

In MR-PROLLIM practice, the log-linear regression may sometimes produce predicted probabilities that are > 1, thus invalidating the MLE procedure. This case may occur, especially when control variables that have strong effects on the exposure or outcome are included. Occurrence of such abnormal probabilities does not necessarily mean the log-linear model is erroneous. It instead indicates that the strong control variables may be put into the regression in improper forms. We list five ways that can be used to avoid abnormal probabilities here:

i. Delete unnecessary strong control variables. A control variable is defined as unnecessary if its removal does not invalidate the assumptions of MR-PROLLIM. Some postzygotic variables, such as age, can be taken as unnecessary because they cannot affect the allele frequencies.
ii. Discretize continuous control variables and replace them with corresponding dummy variables.
iii. Change the forms of control variables. For example, using $\ln(\text{age})$ instead of age will lead to another meaningful interpretation of the regression coefficient and may also avoid abnormal probabilities.
iv. Modify the extreme values of control variables. Sometimes the extreme values (or outlier values) are responsible for abnormal probabilities. Optionally, MR-PROLLIM will force the values of each continuous control variable outside a certain range (e.g., the 2.5–97.5th percentiles) to be the boundary values. This means MR-PROLLIM assumes the "outlier" values on each side to have the same effect as the boundary value.
v. Use Poisson regressions instead of log-linear regressions. The Poisson regression does not model probabilities that should be less than 1 and is therefore less sensitive to extreme values of control variables. If the log-linear model is correct, both regressions can produce consistent effect estimators. The "sandwich" method can be used to construct consistent variance estimators for Poisson regressions (6). But note that Poisson estimators are generally less efficient than log-linear estimators and will thus result in a certain (typically small) loss of statistical power.

**Section S6. Estimation of the prior distribution**

It is required in random-effects MR-PROLLIM without the NOME assumption to estimate the prior distribution of SNP-exposure effects. We assume the prior distribution has the following form:

$$\begin{pmatrix} k_1 \\ k_2 \end{pmatrix} \sim p_0 \begin{pmatrix} 0 \\ 0 \end{pmatrix} + (1 - p_0) N\left( \begin{pmatrix} \mu_0 \\ 2\mu_0 \end{pmatrix}, \begin{pmatrix} s_{01}^2 & s_{01}s_{02}\rho_0 \\ s_{01}s_{02}\rho_0 & s_{02}^2 \end{pmatrix} \right) \quad \text{(S1)}$$

where $p_0$, $\mu_0$, $s_{01}$, $s_{02}$, and $\rho_0$ are five parameters we are interested in. In MR practice, we usually have a selected group of significant and independent SNPs, for which the estimates of $k_1$ and $k_2$ and the corresponding covariance matrix can be calculated. Owing to the SNP selection by Wald $P$ values, the distribution of $k_1$ and $k_2$ for the observed SNPs generally has a volcano-like density surface (the surface is concave around the zero point). To account for this, we derive the conditional density for $\hat{k}_1$ and $\hat{k}_2$ under one-step SNP section as follows:

$$p(\hat{k}_1, \hat{k}_2 | \mathbf{\Sigma}_k, \boldsymbol{O}) = p_0' dN\left( \begin{pmatrix} \hat{k}_1 \\ \hat{k}_2 \end{pmatrix}, \begin{pmatrix} 0 \\ 0 \end{pmatrix}, \mathbf{\Sigma}_k \right) / p_c + (1 - p_0') dN\left( \begin{pmatrix} \hat{k}_1 \\ \hat{k}_2 \end{pmatrix}, \begin{pmatrix} \mu_0 \\ 2\mu_0 \end{pmatrix}, \mathbf{\Sigma}_p + \mathbf{\Sigma}_k \right) / p_{\boldsymbol{O}},$$

$$p_0' = \frac{p_0 p_c}{p_0 p_c + (1 - p_0) p_{\boldsymbol{O}}},$$

$$p_{\boldsymbol{O}} = 1 - \int_{\theta=0}^{\theta=2\pi} \int_{l=0}^{l=l_\theta} dN\left( \begin{pmatrix} l\cos(\theta) \\ l\sin(\theta) \end{pmatrix}, \begin{pmatrix} \mu_0 \\ 2\mu_0 \end{pmatrix}, \mathbf{\Sigma}_p + \mathbf{\Sigma}_k \right) l dl d\theta,$$

$$l_\theta = \sqrt{\frac{\text{qchisq}(1 - p_c, 2)}{\mathbf{\Sigma}_k^{-1}(1,1)\cos^2(\theta) + \mathbf{\Sigma}_k^{-1}(2,2)\sin^2(\theta) + 2\mathbf{\Sigma}_k^{-1}(1,2)\sin(\theta)\cos(\theta)}},$$

where $\boldsymbol{O}$ means "observed", $dN$ is the density function of a bivariate normal distribution, $\mathbf{\Sigma}_k$ is the covariance matrix for $\hat{k}_1$ and $\hat{k}_2$, $p_c$ is the cutoff Wald $P$ value, $\mathbf{\Sigma}_p = \begin{pmatrix} s_{01}^2 & s_{01}s_{02}\rho_0 \\ s_{01}s_{02}\rho_0 & s_{02}^2 \end{pmatrix}$, and qchisq$(\cdot, 2)$ denotes the quantile function of a chi-square distribution with 2 degrees of freedom. MLE can thus be implemented according to the above equations. Sometimes, $p_{\boldsymbol{O}}$ may be very small if a strict $p_c$ has been used. In this case, the error generated by computing $p_{\boldsymbol{O}}$ numerically will exert a great impact on the resultant estimates. To lower such errors, we adopt Farebrother's algorithm (7) to calculate $p_{\boldsymbol{O}}$ using the function "pmvnEll" in R package "shotGroups" (https://CRAN.R-project.org/package=shotGroups; version 0.8.1).

For random-effects MR-PROLLIM with a binary exposure, we additionally need to account for the measurement error of $\hat{\tilde{p}}$ (see **Materials and Methods, Random-effects MR-PROLLIM with a binary exposure**). It is worth mentioning that the SNPs are not directly selected according to $\hat{\tilde{p}}$ (unlike $\hat{k}_1$ and $\hat{k}_2$, $\hat{\tilde{p}}$ is less likely to have a large bias induced by SNP selection) and $\tilde{p}$ itself is a probability that should not be zero (the absolute value of the relative bias is upper bounded). We therefore consider the measurement error of $\hat{\tilde{p}}$ will exert less influence on the causal estimates when the exposure $X$ and outcome $Y$ are not both rare. The parameter $\tilde{p}_i = E(XY|G_i = 0)/E(Y|G_i = 0)$ for the $i$th SNP is partially determined by $\mathbf{k}_{1,-i}$ and $\mathbf{k}_{2,-i}$, where $-i$ denotes "excluding the $i$th SNP". This correlation pattern is complex, and the prior distribution of $\tilde{p}$ is thus hard to model. Considering that if the measurement error is small relative to the prior variance of $\tilde{p}$, the posterior distribution of $\tilde{p}$ given $\hat{\tilde{p}}$ is approximately a normal distribution, we set the prior distribution as:

$$\tilde{p}_i \sim U\left(\max\left(\hat{\tilde{p}}_i - \text{qnorm}(0.5 + p_{\text{cover}}/2)\text{se}_i, 0\right), \min\left(\hat{\tilde{p}}_i + \text{qnorm}(0.5 + p_{\text{cover}}/2)\text{se}_i, 1\right)\right), \text{(S2)}$$

where $U$ denotes the uniform distribution, $\text{qnorm}(\cdot)$ is the quantile function of a standard normal distribution, $p_{\text{cover}}$ is the coverage probability (0.9999 is used as the default), and $\text{se}_i$ is the standard error of $\hat{\tilde{p}}_i$.

**Section S7. Sampling method for MR-PROLLIM without the NOME assumption**

The marginal density $p(Y)$ evaluated at $y_0$ can be approached in probability by $\lim_{n\to\infty} \frac{1}{n}\sum_{i=1}^{n} p(Y = y_0|X = x_i)$, where $\{x_1, \ldots, x_n\}$ is a random sample for $X$. This idea is adopted in random-effects MR-PROLLIM to account for measurement errors of $\hat{k}_1$ and $\hat{k}_2$ (plus $\hat{\tilde{p}}$ for binary exposures) through integrating out the true variables.

We first need to draw random samples according to the posterior density $p\left(k_1, k_2|\hat{k}_1, \hat{k}_2, \mathbf{\Sigma}_k, \boldsymbol{O}\right)$. To do this, we rewrite the density as follows:

$$p\left(k_1, k_2|\hat{k}_1, \hat{k}_2, \mathbf{\Sigma}_k, \boldsymbol{O}\right) = p\left(k_1, k_2|\hat{k}_1, \hat{k}_2, \mathbf{\Sigma}_k\right) = \frac{p(\hat{k}_1, \hat{k}_2|k_1, k_2, \mathbf{\Sigma}_k)p(k_1, k_2|\mathbf{\Sigma}_k)}{p(\hat{k}_1, \hat{k}_2|\mathbf{\Sigma}_k)}.$$

Suppose we have consistently estimated the parameters of the prior distribution **S1** following the method described in **Supplementary Text S6**. We additionally assume the influence of $k_1$ and $k_2$ on $\mathbf{\Sigma}_k$ is ignorable, and $p(k_1, k_2|\mathbf{\Sigma}_k)$ can thus be directly replaced with $p(k_1, k_2)$. We can then draw random samples from the estimated **S1** and implement the acceptance-rejection procedure to reconstruct these samples with $p\left(\hat{k}_1, \hat{k}_2|k_1, k_2, \mathbf{\Sigma}_k\right)$ used as weights. The resultant samples can be viewed as those drawn according to $p\left(k_1, k_2|\hat{k}_1, \hat{k}_2, \mathbf{\Sigma}_k, \boldsymbol{O}\right)$. This procedure will be repeated a certain number of times to reach the desired sample size $n_{\text{post}}$.

In real data analyses, there may be SNPs that have outlier effect sizes, and it will possibly take a large number of repeats to derive the required posterior samples. To lower the computational burden, MR-PROLLIM, by default, first limits the prior samples (originally following **S1**) to be located at the zero point (0,0), which corresponds to zero-effect SNPs, plus an elliptic region centered at $\left(\hat{k}_1, \hat{k}_2\right)$. The shape of this ellipse is determined by $p_{\text{cover}}$ (0.9999 is the default) and

$\mathbf{\Sigma}_k$ to obtain the expected coverage probability under the assumption of $N\left(\left(\hat{k}_1, \hat{k}_2\right)^T, \mathbf{\Sigma}_k\right)$. Once the posterior samples from $p\left(k_1, k_2 | \hat{k}_1, \hat{k}_2, \mathbf{\Sigma}_k, \boldsymbol{O}\right)$ for each SNP have been generated, they will be kept unchanged and used by MR-PROLLIM to approach the following density throughout the MLE procedure:

$$p\left(\hat{m}_1, \hat{m}_2 | \beta_1, \mathbf{\Omega}, \hat{k}_1, \hat{k}_2, \mathbf{\Sigma}_m, \mathbf{\Sigma}_{mk}, \mathbf{\Sigma}_k\right)$$
$$= \operatorname*{plim}_{n\to\infty} \frac{p_1 p_2}{n} \sum_{i=1}^{n} dN\left(\begin{pmatrix} (\beta_1+\mu_1)k_{1i}+\mu_2 \\ (\beta_1+\mu_1)k_{2i}+2\mu_2 \end{pmatrix} + \mathbf{k}_{ci}, \mathbf{\Sigma}_h + \mathbf{\Sigma}_c\right)$$
$$+ \operatorname*{plim}_{n\to\infty} \frac{p_1(1-p_2)}{n} \sum_{i=1}^{n} dN\left(\begin{pmatrix} \beta_1 k_{1i}+\mu_2 \\ \beta_1 k_{2i}+2\mu_2 \end{pmatrix} + \mathbf{k}_{ci}, \mathbf{\Sigma}_h + \mathbf{\Sigma}_c\right)$$
$$+ \operatorname*{plim}_{n\to\infty} \frac{(1-p_1)}{n} \sum_{i=1}^{n} dN\left(\begin{pmatrix} \beta_1 k_{1i} \\ \beta_1 k_{2i} \end{pmatrix} + \mathbf{k}_{ci}, \mathbf{\Sigma}_c\right),$$

where $k_{1i}$ and $k_{2i}$ are posterior samples, $\mathbf{k}_{ci} = \mathbf{\Sigma}_{mk}\mathbf{\Sigma}_k^{-1}\left(\hat{k}_1 - k_{1i}, \hat{k}_2 - k_{2i}\right)^T$, and other notations are consistent with **Formulas 7** and **8** in **Materials and Methods**.

For binary exposures, we can derive the following distribution, which is an analog to **Formula 8**:

$$\begin{pmatrix} \hat{m}_1 \\ \hat{m}_2 \end{pmatrix} | \beta_1, \mathbf{\Omega}, k_1, k_2, \tilde{p}, \hat{k}_1, \hat{k}_2, \hat{\tilde{p}}, \mathbf{\Sigma}_m, \mathbf{\Sigma}_{mk\tilde{p}}, \mathbf{\Sigma}_{k\tilde{p}}$$
$$\sim p_1 p_2 N\left(\begin{pmatrix} \mu_1 k_1 + \mu_2 + g(k_1, \beta_1, \tilde{p}) \\ \mu_1 k_2 + 2\mu_2 + g(k_2, \beta_1, \tilde{p}) \end{pmatrix} + \mathbf{k}_c, \mathbf{\Sigma}_h + \mathbf{\Sigma}_c\right)$$
$$+ p_1(1-p_2) N\left(\begin{pmatrix} \mu_2 + g(k_1, \beta_1, \tilde{p}) \\ 2\mu_2 + g(k_2, \beta_1, \tilde{p}) \end{pmatrix} + \mathbf{k}_c, \mathbf{\Sigma}_h + \mathbf{\Sigma}_c\right)$$
$$+ (1-p_1) N\left(\begin{pmatrix} g(k_1, \beta_1, \tilde{p}) \\ g(k_2, \beta_1, \tilde{p}) \end{pmatrix} + \mathbf{k}_c, \mathbf{\Sigma}_c\right),$$

where $g(k_i, \beta_1, \tilde{p}) = \ln\left((\exp(k_i) - 1)(1 - \exp(-\beta_1))\tilde{p} + 1\right), i = 1,2$, $\mathbf{k}_c = \mathbf{\Sigma}_{mk\tilde{p}}\mathbf{\Sigma}_{k\tilde{p}}^{-1}\left(\hat{k}_1 - k_1, \hat{k}_2 - k_2, \hat{\tilde{p}} - \tilde{p}\right)^T$, $\mathbf{\Sigma}_c = \mathbf{\Sigma}_m - \mathbf{\Sigma}_{mk\tilde{p}}\mathbf{\Sigma}_{k\tilde{p}}^{-1}\mathbf{\Sigma}_{mk\tilde{p}}^T$, and $\mathbf{\Sigma}_m$, $\mathbf{\Sigma}_{mk\tilde{p}}$, and $\mathbf{\Sigma}_{k\tilde{p}}$ are components of the covariance matrix for $\left(\hat{m}_1, \hat{m}_2, \hat{k}_1, \hat{k}_2, \hat{\tilde{p}}\right)$. We additionally assume $\tilde{p}$ follows the prior distribution **S2** and is independent of $k_1$ and $k_2$. A similar sampling procedure is carried out to relax the NOME assumption for random-effects MR-PROLLIM under the double log-linear model.

Note that the resultant log-likelihood function is generally not globally concave in the parameter space. We adopt the "genoud" optimizer, which combines evolutionary algorithms and quasi-Newton methods to search for the global optimum (8). The asymptotic variances can be estimated by inverting the negative Hessian matrix or using the "sandwich" method. As random-effects MR-PROLLIM has a built-in model selection procedure (see **Materials and Methods**) and the final model may deviate from the true model, the inverse Hessian method may thus fail to produce reliable estimates in some cases. Although the "sandwich" method is robust to model misspecifications, it tends to underestimate the variance if the sample size (i.e., SNP number) is small (9). To obtain a more conservative variance estimate, MR-PROLLIM, by default, calculates both estimators and selects the one with a larger value for $\beta_1$. We found MR-PROLLIM with this strategy performed well in simulations (**Figs. 3** and **5C** and **Fig. S2**).

### Section S8. *Post hoc* diagnoses for random-effects MR-PROLLIM with the full or reduced model

Random-effects MR-PROLLIM has a built-in model selection (degeneration) procedure, which runs automatically to handle the cases where some of the estimates reach the boundary values. This procedure works fine asymptotically but may fail to select the true model with finite SNPs. For example, when all SNPs are valid IVs, random-effects MR-PROLLIM sometimes overfits the data using the full or reduced model with a relatively large $\hat{s}_1$. Although the resultant causal estimator may not exhibit a clear median bias, the corresponding $\hat{p}_1\hat{p}_2$, which measures the marginal probability of correlated pleiotropy and the degree of information loss, can be large, thus leading to an inflated variance for $\hat{\beta}_1$. To help avoid the problem of overfitting, we propose a *post hoc* diagnostic routine for random-effects MR-PROLLIM with the full or reduced model (see **Fig. S8**).

As we have discussed in **Supplementary Text S3**, the classification plots of a full model should contain three different groups of points. Similarly, the classification plots of a reduced model should contain two groups (the black and red points; **Fig. S5**). If the three (or two) groups are well differentiated, users can consider the current model appropriate. Otherwise, we recommend users first implement the MR-PROLLIM classical algorithms (i.e., extensions of classical MR methods; **Materials and Methods**) to obtain the heterogeneity $P$. If the result is insignificant, users can fit the intercept model or select between the intercept and Egger models (see **Supplementary Text S3**). If a significant heterogeneity is detected, users can then fit the reduced model and see whether $\mu_1$ deviates significantly from zero. Note that the $P$ value for $\mu_1 \neq 0$ given by the full model is not precise. If there is no evidence in support of a non-zero $\mu_1$, users may fit the zero-correlation model (i.e., zero correlated pleiotropy; $p_2 = 0$ and $\mu_1 = 0$) to increase statistical power. The above diagnostic procedure is a generic one, and users can adjust it according to their own needs, such as swapping the order of classification plots and heterogeneity tests. A similar procedure for the intercept and Egger models has been provided in **Fig. S6**.

In our real data applications, we found seven trait pairs were analyzed with the reduced model and accompanied by large estimates of $p_1 p_2$ (> 0.5; **Tables S1** and **S2**). *Post hoc* diagnoses identified that five of them did not show significant heterogeneity. For one special trait pair (hypertension to heart failure), random-effects MR-PROLLIM originally yielded an RR of 0.10 (95% CI, 0.00–11.03) with $\hat{p}_1 \hat{p}_2 = 0.89$. The $P$ value for heterogeneity among individual effect estimators assuming no horizontal pleiotropy was 0.15, suggesting the assumption might be appropriate. We therefore let MR-PROLLIM fit the intercept model instead and got an RR of 5.72 (95% CI, 4.71–6.73). The other four cases also showed narrowed confidence intervals after this treatment, but there were no changes of significance.

**Section S9. Derivation of the core equations for the double log-linear model**

All MR-PROLLIM methods assuming the double log-linear model depend largely on **Eqs. 12** and **13** in **Materials and Methods**. We additionally assume the support (i.e., the set of possible values) of $G$ does not change with $\mathbf{C}$. The derivation process of these two equations is presented below:

$\because \ln\left(E(X|G_1, G_2, \mathbf{C}, U)\right) = k_1 G_1 + k_2 G_2 + \mathbf{k}_3^T \mathbf{C} + f_3(U)$,

$\therefore E(X|G_1, G_2, \mathbf{C}, U) = P(X = 1|G_1, G_2, \mathbf{C}, U) = \exp\left(k_1 G_1 + k_2 G_2 + \mathbf{k}_3^T \mathbf{C} + f_3(U)\right)$.

$\because \ln\left(E(Y|X, G_1, G_2, \mathbf{C}, U)\right) = \beta_1 X + h_1 G_1 + h_2 G_2 + \boldsymbol{\beta}_2^T \mathbf{C} + f_1(U)$,

$\therefore E(Y|X, G_1, G_2, \mathbf{C}, U) = \exp\left(\beta_1 X + h_1 G_1 + h_2 G_2 + \boldsymbol{\beta}_2^T \mathbf{C} + f_1(U)\right) \Rightarrow$

$P(Y = 1|X = 1, G_1, G_2, \mathbf{C}, U) = \exp\left(\beta_1 + h_1 G_1 + h_2 G_2 + \boldsymbol{\beta}_2^T \mathbf{C} + f_1(U)\right)$.

$\because E(XY|G_1, G_2, \mathbf{C}, U) = P(X = 1, Y = 1|G_1, G_2, \mathbf{C}, U)$

$= \exp\left(\beta_1 + (h_1 + k_1)G_1 + (h_2 + k_2)G_2 + (\boldsymbol{\beta}_2^T + \mathbf{k}_3^T)\mathbf{C} + f_1(U) + f_3(U)\right)$,

$\therefore E(XY|G_1, G_2, \mathbf{C}) = \exp(\beta_1 + (h_1 + k_1)G_1 + (h_2 + k_2)G_2 + (\boldsymbol{\beta}_2^T + \mathbf{k}_3^T)\mathbf{C})\, E\left(\exp\left(f_1(U) + f_3(U)\right)|\mathbf{C}\right)$,

$E(XY|G = 1, \mathbf{C})/E(XY|G = 0, \mathbf{C}) = \exp(h_1 + k_1)$,

$E(XY|G = 2, \mathbf{C})/E(XY|G = 0, \mathbf{C}) = \exp(h_2 + k_2)$.

$\because E(Y|G_1, G_2, \mathbf{C}, U) = \exp\left(h_1 G_1 + h_2 G_2 + \boldsymbol{\beta}_2^T \mathbf{C} + f_1(U)\right) E(\exp(\beta_1 X)|G_1, G_2, \mathbf{C}, U)$

$= \exp\left(h_1 G_1 + h_2 G_2 + \boldsymbol{\beta}_2^T \mathbf{C} + f_1(U)\right)\left((\exp(\beta_1) - 1)\exp\left(k_1 G_1 + k_2 G_2 + \mathbf{k}_3^T \mathbf{C} + f_3(U)\right) + 1\right)$

$= (\exp(\beta_1) - 1)\exp\left((h_1 + k_1)G_1 + (h_2 + k_2)G_2 + (\boldsymbol{\beta}_2^T + \mathbf{k}_3^T)\mathbf{C} + f_1(U) + f_3(U)\right)$

$+ \exp\left(h_1 G_1 + h_2 G_2 + \boldsymbol{\beta}_2^T \mathbf{C} + f_1(U)\right)$,

$\therefore E(Y|G_1, G_2, \mathbf{C}) =$

$$(\exp(\beta_1)-1)\exp\left((h_1+k_1)G_1+(h_2+k_2)G_2+(\boldsymbol{\beta}_2^T+\mathbf{k}_3^T)\mathbf{C}\right)E\left(\exp\left(f_1(U)+f_3(U)\right)|\mathbf{C}\right)$$
$$+\exp(h_1G_1+h_2G_2+\boldsymbol{\beta}_2^T\mathbf{C})\,E\left(\exp\left(f_1(U)\right)|\mathbf{C}\right).$$

Substituting $E(XY|G_1,G_2,\mathbf{C})$ into $E(Y|G_1,G_2,\mathbf{C})$, we can get:

$$E(Y|G_1,G_2,\mathbf{C})=(1-\exp(-\beta_1))E(XY|G_1,G_2,\mathbf{C})+\exp(h_1G_1+h_2G_2+\boldsymbol{\beta}_2^T\mathbf{C})\,E\left(\exp\left(f_1(U)\right)|\mathbf{C}\right).$$
$$\therefore E(Y|G=0,\mathbf{C})=(1-\exp(-\beta_1))E(XY|G=0,\mathbf{C})+\exp(\boldsymbol{\beta}_2^T\mathbf{C})\,E\left(\exp\left(f_1(U)\right)|\mathbf{C}\right),$$
$$E(Y|G=1,\mathbf{C})=(1-\exp(-\beta_1))E(XY|G=1,\mathbf{C})+\exp(h_1+\boldsymbol{\beta}_2^T\mathbf{C})\,E\left(\exp\left(f_1(U)\right)|\mathbf{C}\right),$$
$$E(Y|G=2,\mathbf{C})=(1-\exp(-\beta_1))E(XY|G=2,\mathbf{C})+\exp(h_2+\boldsymbol{\beta}_2^T\mathbf{C})\,E\left(\exp\left(f_1(U)\right)|\mathbf{C}\right).$$

$$\therefore E(Y|G=1,\mathbf{C})=\left(E(Y|G=0,\mathbf{C})-(1-\exp(-\beta_1))E(XY|G=0,\mathbf{C})\right)\exp(h_1)$$
$$+(1-\exp(-\beta_1))E(XY|G=1,\mathbf{C})$$
$$=\left(E(Y|G=0,\mathbf{C})-(1-\exp(-\beta_1))E(XY|G=0,\mathbf{C})\right)\exp(h_1)$$
$$+(1-\exp(-\beta_1))E(XY|G=0,\mathbf{C})\exp(k_1)\exp(h_1)$$
$$=\left((\exp(k_1)-1)(1-\exp(-\beta_1))E(XY|G=0,\mathbf{C})+E(Y|G=0,\mathbf{C})\right)\exp(h_1).$$

Similarly, the following equation holds:

$$E(Y|G=2,\mathbf{C})=\left((\exp(k_2)-1)(1-\exp(-\beta_1))E(XY|G=0,\mathbf{C})+E(Y|G=0,\mathbf{C})\right)\exp(h_2).$$

Finally, integrating out $\mathbf{C}$ given $G=0$, we can get:

$$\ln\left(\frac{E(E(Y|G=1,\mathbf{C})|G=0)}{E(Y|G=0)}\right)=\ln\left((\exp(k_1)-1)(1-\exp(-\beta_1))\frac{E(XY|G=0)}{E(Y|G=0)}+1\right)+h_1,$$
$$\ln\left(\frac{E(E(Y|G=2,\mathbf{C})|G=0)}{E(Y|G=0)}\right)=\ln\left((\exp(k_2)-1)(1-\exp(-\beta_1))\frac{E(XY|G=0)}{E(Y|G=0)}+1\right)+h_2. \qquad \square$$

**Section S10. Consistent estimators of the iterative expectations**

Consistent estimators of two iterative expectations, $E(E(Y|G=1,\mathbf{C})|G=0)$ and $E(E(Y|G=2,\mathbf{C})|G=0)$, are required to validate the MR-PROLLIM methods assuming the double log-linear model. For simplicity, we first consider there is only one continuous control variable $C$. Suppose there is an equidistant partition of the real number axis for $C$, which results in $n$ intervals that have the same length $\Delta C$ and median values $C_i, i=1,\dots,n$. We randomly draw $N_1$ individuals (type A) with $G=1$ and $N_0$ individuals (type B) with $G=0$. According to $C$, a total of $n_{1i}$ type A individuals and $n_{0i}$ type B individuals are grouped in the $i$th interval. The following equation holds:

$$E(E(Y|G=1,C)|G=0)$$
$$=\int_{-\infty}^{\infty}E(Y|G=1,C)p(C|G=0)\,dC$$
$$=\lim_{n\to\infty}\sum_{i=1}^{n}E(Y|G=1,C_i)\,p(C_i|G=0)\Delta C, \qquad \text{(S3)}$$

where $p(C_i|G=0)$ is the density of $C$ evaluated at $C_i$ given $G=0$. We define a random weight $W_{i,j}=\frac{S_{i,j}/N_0}{1/N_1}=\frac{S_{i,j}N_1}{N_0}\geq 0$, where $S_{i,j}$ is a random variable that satisfies $\sum_{j=1}^{n_{1i}} S_{i,j}=n_{0i}$ and is independent of $Y_{i,j,G=1}$ (i.e., the outcome value of the $j$th type A individual in the $i$th interval) when $i$ is held constant. Then, the following equation holds:

$$E(Y|G=1,C_i)=\underset{n,n_{1i}\to\infty}{\operatorname{plim}}\frac{1}{n_{1i}}\sum_{j=1}^{n_{1i}}\frac{S_{i,j}n_{1i}}{n_{0i}}Y_{i,j,G=1}$$

$$=\underset{n,n_{1i}\to\infty}{\operatorname{plim}}\sum_{j=1}^{n_{1i}}\frac{N_0W_{i,j}}{n_{0i}N_1}Y_{i,j,G=1}. \quad \text{(S4)}$$

Substituting **Eq. S4** and $p(C_i|G=0)=\underset{n,n_{0i}\to\infty}{\operatorname{plim}}\frac{n_{0i}}{N_0\Delta C}$ into **Eq. S3**, we can get:

$$E(E(Y|G=1,C)|G=0)=\underset{n,n_{1i},n_{0i}\to\infty}{\operatorname{plim}}\sum_{i=1}^{n}\sum_{j=1}^{n_{1i}}\frac{N_0W_{i,j}}{n_{0i}N_1}Y_{i,j,G=1}\frac{n_{0i}}{N_0}$$

$$=\underset{n,n_{1i},n_{0i}\to\infty}{\operatorname{plim}}\frac{1}{N_1}\sum_{i=1}^{n}\sum_{j=1}^{n_{1i}}Y_{i,j,G=1}W_{i,j}$$

$$=\underset{N_0,N_1\to\infty}{\operatorname{plim}}\frac{1}{N_1}\sum_{l=1}^{N_1}Y_{l,G=1}W_l. \quad \text{(S5)}$$

MR-PROLLIM calculates $W_l$ according to the following clustering procedure:

i. Set $C_l$ ($l=1,\ldots,N_1$) of each type A individual as the cluster center.
ii. Allocate type B individuals to these cluster centers according to Euclidean distances. If $\geq$ 2 centers (e.g., $n_e$ such centers) share the shortest distance, then $\frac{1}{n_e}$ will be given to all of them.
iii. The weight $W_l$ is calculated as the number of type B individuals allocated to the $l$th center (i.e., $S_l$) multiplied by $\frac{N_1}{N_0}$. Note that $S_l$ may be a fraction due to shared shortest distances.

After determining the weights, MR-PROLLIM calculates the estimator of $E(E(Y|G=1,C)|G=0)$ according to **Eq. S5**. The consistency can be more easily seen if $C$ is a categorical variable with $n_c$ levels. In this case, the estimator is equivalent to $\sum_{i=1}^{n_c}\hat{p}(Y=1|G=1,C=i)\hat{p}(C=i|G=0)$, where $\hat{p}$ denotes the corresponding moment estimator. If there are more control variables, MR-PROLLIM first standardizes all continuous variables (to mean 0 and variance 1) and then carries out the aforementioned clustering procedures within strata of categorical control variables. The estimator of $E(E(Y|G=2,C)|G=0)$ can be derived similarly.

MR-PROLLIM computes the estimate of the required covariance matrix based on first-order Taylor expansions. **Eq. S5** can be further transformed to:

$$\underset{N_0,N_1\to\infty}{\operatorname{plim}}\frac{1}{N_1}\sum_{l=1}^{N_1}Y_{l,G=1}W_l=\underset{N_0,N_1\to\infty}{\operatorname{plim}}\frac{1}{N_1}\sum_{l=1}^{N_1}Y_{l,G=1}\frac{S_lN_1}{N_0}=\underset{N_0,N_1\to\infty}{\operatorname{plim}}\sum_{l=1}^{N_1}Y_{l,G=1}\frac{S_l}{N_0}. \quad \text{(S6)}$$

Note that $\sum_{l=1}^{N_1} \frac{S_l}{N_0} = 1$, and $S_l$ and $S_{l+1}$ are thus correlated with each other. We replace $\frac{S_l}{N_0}$ with $\frac{S_l'}{\sum_{l=1}^{N_1} S_l'}$, where $\{S_1', \ldots, S_{N_1}'\}$ is a sequence of independent and identically distributed random variables. Then, $\sum_{l=1}^{N_1} Y_{l,G=1} \frac{S_l}{N_0}$ can be rewritten as:

$$\sum_{l=1}^{N_1} Y_{l,G=1} \frac{S_l}{N_0} = \sum_{l=1}^{N_1} Y_{l,G=1} \frac{S_l'}{\sum_{l=1}^{N_1} S_l'} = \frac{\frac{1}{N_1}\sum_{l=1}^{N_1} Y_{l,G=1} S_l'}{\frac{1}{N_1}\sum_{l=1}^{N_1} S_l'}$$

$$\approx q_0 + \frac{q_1}{N_1}\sum_{l=1}^{N_1} Y_{l,G=1} S_l' + \frac{q_2}{N_1}\sum_{l=1}^{N_1} S_l'$$

$$= \frac{1}{N_1}\sum_{l=1}^{N_1}\left(q_0 + q_1 Y_{l,G=1} S_l' + q_2 S_l'\right)$$

$$= \frac{1}{N_1}\sum_{l=1}^{N_1} Y_{l,G=1}'$$

$$= \frac{\frac{1}{N}\sum_{i=1}^{N} Y_i'' G_{1,i}}{\frac{N_1}{N}}, \tag{S7}$$

where $q_0 = \frac{\frac{1}{N_1}\sum_{l=1}^{N_1} y_{l,G=1} s_l'}{\frac{1}{N_1}\sum_{l=1}^{N_1} s_l'}$, $q_1 = \frac{1}{\frac{1}{N_1}\sum_{l=1}^{N_1} s_l'}$, $q_2 = -q_0 q_1$, $N$ is the total sample size (individuals with $G \in \{0, 1, 2\}$), and $Y_i''$ is an expansion of $Y_{l,G=1}' = q_0 + q_1 Y_{l,G=1} S_l' + q_2 S_l'$, which has the same value as $Y_{l,G=1}'$ for the same individual with $G = 1$ but is also defined for individuals with $G \in \{0, 2\}$. Again, using the first-order Taylor expansion, we can approximate $\frac{\frac{1}{N}\sum_{i=1}^{N} Y_i'' G_{1,i}}{\frac{N_1}{N}}$ with $\frac{1}{N}\sum_{i=1}^{N}\left(q_0' + q_1' Y_i'' G_{1,i} + q_2' G_{1,i}\right)$, where $q_0' = \frac{1}{N_1}\sum_{l=1}^{N_1} y_{l,G=1}'$, $q_1' = \frac{N}{N_1}$, and $q_2' = -q_0' q_1'$.
Therefore, the following equation holds:

$$E(E(Y|G = 1, C)|G = 0) = \operatorname*{plim}_{N\to\infty} \frac{1}{N}\sum_{i=1}^{N} q_0' + q_1' Y_i'' G_{1,i} + q_2' G_{1,i}. \tag{S8}$$

However, $S_l'$ is a hypothetical variable that is not actually observed. If the sample size $N$ is relatively large, the correlation between $S_l$ and $S_{l+1}$ is small, and $S_l'$ can be replaced with $S_l$ for variance-covariance estimations. The above expectation and variance estimators have been examined with simulations.

Note that MR-PROLLIM adopts first-order Taylor expansions and expansions of random variables defined in subgroups to the whole sample (if necessary) to generate moment estimators as approximations throughout the variance-covariance estimating procedures, not only for the procedure described in this section.

**Section S11. Empirical analyses partially supporting the PAHP assumption**

Ordinary fixed-effects MR-PROLLIM requires the PAHP assumption to ensure consistency. We have shown in **Table S8** that most SNPs appear to have perfectly additive effects on traits, suggesting the pleiotropic effects may also mainly exhibit additive patterns. However, the fixed-effects MR-PROLLIM estimating framework requires an additional SNP selection [required for the noncollinearity condition (continuous exposures) or the root selection procedure (binary exposures); **Materials and Methods**], which will lead to deviations of the SNP-exposure effects from additive patterns among suitable SNPs. We thus conducted several empirical analyses to account for this SNP selection.

As we have discussed in the main text, HDL-C is considered to exert no effect on MI or T2D, and the trait pair of birth weight to SBP is also classified as "non-causal". If no adjustment is made, traditional MR methods will easily obtain significant results for these three trait pairs, indicating possibly strong horizontal pleiotropy. Therefore, with these trait pairs, we can explore the potential connections between the SNP-exposure effect patterns and the pleiotropic effect patterns.

We conducted two types of analyses. The first one was to investigate the potential associations between $\hat{R}_k = \frac{\hat{k}_2}{\hat{k}_1}$ and $\hat{R}_h = \frac{\hat{h}_2}{\hat{h}_1}$ (estimated by SNP-outcome effect ratios among non-causal trait pairs). And the second one was to see whether nominal deviations of SNP-exposure effects from the additive pattern would lead to similar deviations of horizontal pleiotropy. Nominal deviations were judged according to the $Z$ statistic described in **Supplementary Text S4**. We additionally included the trait pair of BMI to SBP as a positive control because we expected there was a strong causal effect which would link the SNP-exposure and SNP-outcome effect ratios. The results are presented in **Fig. S9** and **Table S9**, respectively.

**Section S12. Derivation of the disproportion assumption for continuous exposures**

Suppose all assumptions in **Materials and Methods, Fixed-effects MR-PROLLIM with a continuous exposure** hold. Then the log-linear regression of $Y$ on $\hat{\hat{G}}$, $G$, and **C** according to **Eq. 15** is equivalent to solving the following equations:

$$\hat{\beta}_1 \hat{k}_1 + \hat{h} = \hat{m}_1, \tag{S9}$$

$$\hat{\beta}_1 \hat{k}_2 + 2\hat{h} = \hat{m}_2, \tag{S10}$$

where $\hat{k}_1$ and $\hat{k}_2$ are consistent estimates of $k_1$ and $k_2$, respectively, $\hat{k}_2$ is not equal to $2\hat{k}_1$ (the noncollinearity condition), and $\hat{m}_1$ and $\hat{m}_2$ are derived through a log-linear regression of $Y$ on $G_1$, $G_2$, and **C**. The solutions to **Eqs. S9** and **S10** are:

$$\hat{h} = \frac{\widehat{m}_2\hat{k}_1 - \widehat{m}_1\hat{k}_2}{2\hat{k}_1 - \hat{k}_2}, \tag{S11}$$

$$\hat{\beta}_1 = (2\widehat{m}_1 - \widehat{m}_2)/(2\hat{k}_1 - \hat{k}_2). \tag{S12}$$

If the PAHP assumption is violated but the other assumptions remain valid, then $m_1$ equals $\beta_1 k_1 + h_1$ and $m_2$ equals $\beta_1 k_2 + h_2$, where $h_2$ does not have to be $2h_1$. In this case, $\hat{h}$ converges in probability to zero if and only if $m_2 k_1$ equals $m_1 k_2$, which is equivalent to $h_2 k_1 = h_1 k_2$. Therefore, if $h_2 k_1 \neq h_1 k_2$, $\hat{h}$ can be used to test the existence of horizontal pleiotropy.

**Section S13. Root selection strategy of fixed-effects MR-PROLLIM for binary exposures**

The root selection is a unique procedure required by fixed-effects MR-PROLLIM for binary exposures. For each significant and independent SNP, **Eq. 16**, combined with **Eq. 12**, generally outputs two sets of roots as follows:

$$\hat{\hat{\beta}}_1 = \frac{\hat{\hat{m}}\frac{\hat{\hat{k}}_2}{\hat{\hat{k}}_1} - 2 \pm \sqrt{\left(\hat{\hat{m}}\frac{\hat{\hat{k}}_2}{\hat{\hat{k}}_1} - 2\right)^2 - 4\left(1 - \hat{\hat{m}}\right)}}{2\hat{\hat{k}}_1}, \tag{S13}$$

$$\exp\left(\hat{h}\right) = \frac{\exp(\widehat{m}_1)}{\hat{\hat{k}}_1\hat{\hat{\beta}}_1 + 1}, \tag{S14}$$

or sometimes no real root if $\Delta = \left(\hat{\hat{m}}\frac{\hat{\hat{k}}_2}{\hat{\hat{k}}_1} - 2\right)^2 - 4\left(1 - \hat{\hat{m}}\right) < 0$. To reduce the median biases of the above estimators, MR-PROLLIM applies the following SNP inclusion criteria by default:

i. The Bonferroni-corrected $P$ value of testing the null hypothesis that $\tilde{k}_1 = 0$ is less than 0.1.
ii. The simulated probability for $\Delta < 0$ by parametric bootstrap is less than 0.01.

The resultant suitable SNPs will be put into the root selection procedure according to $Q$ statistics. This procedure aims to find the root combination that exhibits the lowest heterogeneity on $\hat{\hat{\beta}}_1$. However, traversing all possible root combinations may be time-consuming if the number of suitable SNPs is large. We thus adopt a randomized algorithm where SNPs are randomly split into a number of approximately even subgroups, within which root selections are conducted, and all subgroup optimal results are afterward combined together. This procedure is repeated a certain number of times (denoted as $n_{\mathrm{rep}}$), and the combination with the lowest $Q$ is the final output. By default, MR-PROLLIM sets the maximum number of combinations in each subgroup to $2^{16}$ (i.e., 16 SNPs in a subgroup) and limits the probability that two SNPs are not tested together to be < 1E-5. For simplicity, this probability is calculated as follows:

$$\left(\frac{n_{\mathrm{snp}} - m}{n_{\mathrm{snp}} - 1}\right)^{n_{\mathrm{rep}}},$$

where $n_{\mathrm{snp}}$ denotes the total number of suitable SNPs, and $m$ is the number of SNPs in each subgroup (the minimum number will be used if subgroup SNP numbers are not equal). Note that the parameter $\tilde{\beta}_1 = 1 - \exp(-\beta_1)$ is < 1 and $\exp(h)$ is > 0. For a certain set of roots, if the corresponding estimates significantly (Bonferroni-corrected one-sided $P$ value < 0.05) violate either of these two conditions, this set will not be included in the above root selection procedure.

**Section S14. Derivation of the disproportion assumption for binary exposures**

Just like the case of dealing with a continuous exposure, the robust fixed-effects MR-PROLLIM for binary exposures also requires a core assumption to keep $\hat{h}$ as a valid measure of horizontal pleiotropy under model misspecifications. We only consider violations of the PAHP assumption. According to **Eqs. S13** and **S14**, $\exp(\hat{h})$ can be rewritten as follows:

$$\exp(\hat{h}) = \frac{2\exp(\hat{m}_1)}{\hat{\hat{m}}\frac{\hat{\hat{k}}_2}{\hat{\hat{k}}_1} \pm \sqrt{\left(\hat{\hat{m}}\frac{\hat{\hat{k}}_2}{\hat{\hat{k}}_1} - 2\right)^2 - 4(1-\hat{\hat{m}})}}. \quad \text{(S15)}$$

Note that we assume there exist two real roots for $\exp(\hat{h})$. If at least one root equals 1, the following equations hold:

$$2\exp(\hat{m}_1) = \hat{\hat{m}}\frac{\hat{\hat{k}}_2}{\hat{\hat{k}}_1} \pm \sqrt{\left(\hat{\hat{m}}\frac{\hat{\hat{k}}_2}{\hat{\hat{k}}_1} - 2\right)^2 - 4(1-\hat{\hat{m}})} \Rightarrow$$

$$2 = \exp(\hat{m}_1 - \hat{m}_2)\frac{\hat{\hat{k}}_2}{\hat{\hat{k}}_1} \pm \sqrt{\left(\exp(\hat{m}_1 - \hat{m}_2)\frac{\hat{\hat{k}}_2}{\hat{\hat{k}}_1}\right)^2 - 4\exp(-\hat{m}_2)\frac{\hat{\hat{k}}_2}{\hat{\hat{k}}_1} + 4\exp(-\hat{m}_2)} \Rightarrow$$

$$\left(\exp(\hat{m}_1 - \hat{m}_2)\frac{\hat{\hat{k}}_2}{\hat{\hat{k}}_1} - 2\right)^2 = \left(\exp(\hat{m}_1 - \hat{m}_2)\frac{\hat{\hat{k}}_2}{\hat{\hat{k}}_1}\right)^2 - 4\exp(-\hat{m}_2)\frac{\hat{\hat{k}}_2}{\hat{\hat{k}}_1} + 4\exp(-\hat{m}_2) \Rightarrow$$

$$\hat{\hat{k}}_1(\exp(\hat{m}_2) - 1) = \hat{\hat{k}}_2(\exp(\hat{m}_1) - 1) \Rightarrow$$

$$\left(\exp(\hat{k}_1) - 1\right)(\exp(\hat{m}_2) - 1) = \left(\exp(\hat{k}_2) - 1\right)(\exp(\hat{m}_1) - 1).$$

It can be seen from the above derivation that if $\left(\exp(\hat{k}_1) - 1\right)(\exp(\hat{m}_2) - 1)$ equals $\left(\exp(\hat{k}_2) - 1\right)(\exp(\hat{m}_1) - 1)$, at least one of the roots for $\exp(\hat{h})$ equals 1, which means the two propositions are equivalent. Considering $\hat{k}_1$, $\hat{k}_2$, $\hat{m}_1$, and $\hat{m}_2$ are consistent estimators if other assumptions of fixed-effects MR-PROLLIM still hold (see **Materials and Methods, Fixed-effects MR-PROLLIM with a binary exposure**), we conclude that $(\exp(k_1) - 1)(\exp(m_2) - 1) \neq (\exp(k_2) - 1)(\exp(m_1) - 1)$, a sufficient and necessary condition, ensures both roots for $\hat{h}$ converge in probability to non-zero values.

**Section S15. Correlations among the optimal linear combination estimators**

Classical outlier-robust methods that assume the double log-linear model require initial individual estimators under the assumption of no horizontal pleiotropy. MR-PROLLIM, by default, adopts the optimal linear combination estimators given by **Eq. 18**. We have found in simulations that these estimators exhibit a non-zero but generally very weak correlation with each other. Consider the following example:

$$U_1, U_2 \overset{\text{i.i.d.}}{\sim} U(0,1),$$
$$P_X = \exp(0.5G_1 + 0.5G_2 + U_1 - 3),$$
$$P_Y = \exp(X + U_1 + U_2 - 3),$$

where $G_1$ and $G_2$ are two independent SNPs with minor allele frequencies of 0.3 and 0.5, respectively. The corresponding simulation results are provided in **Table S10**. We detected a significant linear correlation between $\hat{\hat{\beta}}_{1,\mathrm{opt},1}$ and $\hat{\hat{\beta}}_{1,\mathrm{opt},2}$, and this correlation did not seem to shrink as the sample size grew. On the contrary, we obtained no significant finding in the simulated case for a continuous exposure. The data-generating process was as follows:

$$U_1, U_2 \overset{\text{i.i.d.}}{\sim} U(0,1),$$
$$X = 0.1G_1 + 0.1G_2 + U_1,$$
$$P_Y = \exp(X + U_1 + U_2 - 3.4).$$

This phenomenon indicates that such correlations should exist specifically in the double log-linear model. Although the correlation among individual estimators does not affect the consistency, it may introduce bias to the confidence interval. We therefore additionally compute the asymptotic covariance matrix for these individual estimators according to first-order Taylor expansions and adjust the estimation procedures as follows:

i. Simple IVW: individual estimators from different SNPs are combined according to **Eq. 18**.
ii. Weighted mode and median: the estimated covariance matrix is used in the bootstrap procedure to generate correlated individual estimates; the weights are not changed.
iii. $Q$ statistic-based outlier removal: we recalculate the statistic according to the Wald test as follows: $Q' = \left(\mathbf{R}\hat{\hat{\boldsymbol{\beta}}}_{1,\mathrm{opt}}\right)^T (\mathbf{R}\boldsymbol{\Sigma}\mathbf{R}^T)^{-1}\mathbf{R}\hat{\hat{\boldsymbol{\beta}}}_{1,\mathrm{opt}}$, where $\mathbf{R} = (\mathbf{I}_{m\times 1}, -\mathbf{I}_{m\times m})$, $\mathbf{I}_{m\times 1}$ is the $m$-dimensional column vector of ones, $\mathbf{I}_{m\times m}$ is the $m$-dimensional identity matrix, $m$ is the SNP number minus 1, and $\boldsymbol{\Sigma}$ denotes the estimated covariance matrix. $Q'$ asymptotically follows $\chi^2(m)$ under the null hypothesis that there is no heterogeneity.

As the random-effects model incorporates an additional between-SNP variance [i.e., $\tau^2$ in **Formula 17**], which will further dilute the weak correlations, we do not carry out analogous adjustments for the additive random-effects combination. Note that the problem of correlations among individual estimators also exists in fixed-effects MR-PROLLIM. We therefore add similar adjustment steps to this method as well.

### Section S16. Setting cutoff *P* values for MR-PROLLIM

MR-PROLLIM currently requires individual-level data as the input but is not designed to perform fast *de novo* GWAS. To reduce the time consumption, we have recommended the two-step SNP selection procedure in **Materials and Methods, SNP selection for MR-PROLLIM**. Briefly, the SNP selection consists of two statistical tests in series, of which the first one is already performed or will be performed using fast algorithms. In this section, we focus on two issues that arise from sequential testing. The first one is the influence on prior estimations described in **Supplementary Text S6**. The second one is the additional steps required for controlling the type I error. Both issues are closely related to $P$ thresholds for MR-PROLLIM.

***Influence on the prior estimation***

The method proposed in **Supplementary Text S6** is primarily designed for one-step SNP selection according to Wald $P$ values. It remains valid If the first-step and second-step tests are equivalent. However, if these two tests are independent or just correlated with each other, the prior distribution of $k_1$ and $k_2$ after the first-step selection will deviate from **Formula S1** and turn to a mixture of the zero point $(0,0)$ and a volcano-like distribution. We consider our prior estimation method is relatively robust to this deviation because:

i. The crater of the volcano-like distribution can be partially filled with the zero points to imitate a normal distribution. This indicates a weight transfer from $(0,0)$ to the volcano-like distribution but also a stronger similarity to **Formula S1**.
ii. If the second-step $P$ is strict relative to the first-step $P$, both the filled volcano-like and the normal distributions will yield similar posterior distributions. This is because most SNPs located at the original concave region will be ruled out, and differences in this region are unimportant.

Therefore, it is recommended to adopt a relatively strict second-step cutoff $P$ value. However, figuring out how strict it should be is a complicated task. As a reference, we have generally used $P_{c2} \leq P_{c1,\mathrm{adj}}/100$ for two independent tests in our simulations and $P_{c2} \leq P_{c1,\mathrm{adj}}/10$ for correlated tests in the real data analyses (we refer to these inequalities as Constraint 1). $P_{c2}$ here denotes the second-step $P$ threshold, and $P_{c1,\mathrm{adj}}$ is the sample size-adjusted first-step $P$ threshold calculated as follows:

$$P_{c1,\mathrm{adj}} = 1 - \mathrm{pchisq}\left(\frac{n_2}{n_1}\mathrm{qchisq}(1 - P_{c1}, 2), 2\right),$$

where pchisq and qchisq denote distribution and quantile functions of a chi-square distribution, respectively, $P_{c1}$ is the actually used first-step $P$ threshold, $n_1$ is the first-step sample size, and $n_2$ is the sample size for MR-PROLLIM.

Note that Constraint 1 is recommended only for random-effects MR-RPOLLIM without the NOME assumption but not for other MR-PROLLIM methods. If $n_2$ is sufficiently large, the normal posterior distribution or the NOME assumption may approximately hold. In this case, Constraint 1 is considered unnecessary.

***Steps for controlling the type I error***

The widely used threshold 5E-8 was proposed mainly for controlling the inflated type I error due to multiple testing among approximately one million independent common SNPs. In MR-PROLLIM practice, the probability that one zero-effect SNP finally passes the selection (denoted as $P_{\text{fp}}$, with fp meaning "false positive") is $\min(P_{c1}, P_{c2})$ if the two tests are equivalent and $P_{c1}P_{c2}$ if the two tests are independent. However, as we have encountered in the real data applications, the two tests may just correlate with each other due to imperfect sample overlap and differences in statistical models.

We therefore propose a parametric bootstrap-based method to estimate $P_{\text{fp}}$ with summary data from the first test and individual data from the second test. As $P_{c2}$ is generally lower than $P_{c1}$, we rewrite $P_{\text{fp}}$ as $P(P_{t1} < P_{c1}|P_{t2} < P_{c2})P_{c2}$, where $P_{t1}$ and $P_{t2}$ denote the $P$ values of the first-step and second-step tests, respectively. If we know the joint distribution of both test statistics, we can draw random samples to get a simulated estimate of $P(P_{t1} < P_{c1}|P_{t2} < P_{c2})$ and thus calculate $\hat{P}_{\text{fp}} = \hat{P}(P_{t1} < P_{c1}|P_{t2} < P_{c2})P_{c2}$. To do this, we first obtain a random sample of zero-effect SNPs, as well as their regression coefficients ($\hat{k}_{0i}$) and corresponding standard errors ($\text{se}_{0i}$) from the first-step tests. We next perform linear or log-linear regressions (the second-step tests) to get $\hat{k}_{1i,s} = \hat{k}_{1i}/\text{se}_{1i}$, $\hat{k}_{2i,s} = \hat{k}_{2i}/\text{se}_{2i}$, and the correlation matrix $\mathbf{\Sigma}_{ki,s}$ ($s$ denotes "scaled"). We assume:

$$\left(\hat{k}_{0,s}, \hat{k}_{1,s}, \hat{k}_{2,s}\right)^T \sim N\left((0,0,0)^T, \begin{pmatrix} 1 & \mathbf{\Sigma}_{0k,s} \\ \mathbf{\Sigma}_{0k,s}^T & \mathbf{\Sigma}_{k,s} \end{pmatrix}\right), \tag{S16}$$

where $\hat{k}_{0,s} = \hat{k}_0/\text{se}_0$, and $\mathbf{\Sigma}_{0k,s}$ denotes the covariance of $\hat{k}_{0,s}$ with $\hat{k}_{1,s}$ and $\hat{k}_{2,s}$. We estimate $\mathbf{\Sigma}_{0k,s}$ using the sample correlation matrix of $\left(\hat{k}_{0,s}, \hat{k}_{1,s}, \hat{k}_{2,s}\right)$ and $\mathbf{\Sigma}_{k,s}$ using the mean value of $\left\{\mathbf{\Sigma}_{ki,s}\right\}$. Afterward, we first simulate random samples from the region where $P_{t2} < P_{c2}$ holds (an elliptic ring that lies within the complementary region of the ellipse and has a default coverage of 0.9999), then calculate the probability that $P_{t1} < P_{c1}$ for each sample according to the conditional form of **Formula S16**, and eventually average these probabilities to derive $\hat{P}(P_{t1} < P_{c1}|P_{t2} < P_{c2})$ and $\hat{P}_{\text{fp}}$.

As a general rule, $\hat{P}_{\text{fp}}$ should be less than $\alpha/n_{\text{ind}}$ (Constraint 2), where $\alpha$ is the desired significance level (e.g., 0.05 or 0.1), and $n_{\text{ind}}$ denotes the number of independent SNPs. Note that $n_{\text{ind}}$ may derivate from one million, depending on the SNP inclusion criteria. Therefore, it is recommended to check $n_{\text{ind}}$ with the full list of SNPs for the first-step tests before making a decision. Users may carry out LD clumping procedures to estimate $n_{\text{ind}}$.

In addition to the aforementioned constraints, there are several other things worth attention for setting $P_{c1}$ and $P_{c2}$, such as the number of candidate SNPs and $\chi^2$ of the second-step test. We recommend a lax $P_{c1}$ (e.g., we generally used 1E-3 for UKB GWAS summary data) to include several thousand candidate SNPs and a relatively strict $P_{c2}$ to make $\chi^2 > 20$, which is similar to $F$ statistic > 10 and equivalent to $P_{c2} < 4.54\text{E-}5$. However, as random-effects MR-PROLLIM without the NOME assumption can account for zero-effect SNPs via incorporating the prior distribution **S1**. The threshold $P_{c2}$ may be set at a higher level than that used for other MR-PROLLIM methods to enlarge the final SNP number. We suggest more than 50 SNPs should be included for random-effects MR-PROLLIM, as we have observed fine performances of this method in simulations with 50–130 SNPs (**Fig. 3**). A summary of the recommended restrictions on the second-step $P$ threshold is given in **Table S11**.

### Section S17. Data-generating procedure for the main simulations

Following (5), we first randomly collected a group of 10,000 individuals from the UKB participants and obtained the genotype data containing 19,020 HapMap SNPs [https://ftp.ncbi.nlm.nih.gov/hapmap/genotypes/hapmap3_r3; minor allele frequency (MAF) > 0.05 and missing rate < 0.05] on chromosome 19. We then repeated this data batch 30 times to create a dataset of 570,600 SNPs. To ensure independence among the between-batch SNPs, genotype data were generated randomly for each batch and eventually combined together.

Control variables are generally required in GWAS. For simplicity, we only simulated two control variables. The first one, $C_1 \sim N(0,1)$, was a continuous variable, which contributed to the variations of $X$ and $Y$ but was independent of the genotypes. Controlling for $C_1$ could help reduce standard errors of the SNP effect estimates, especially for continuous traits. The second one, $C_2$, was a categorical variable, which measured the genetic backgrounds of individuals. To define this variable, we applied k-means clustering to the first ten genetic principal components provided by UKB and extracted three different "ancestral groups" ("kmeans" function in the "stats" R package; 3 clusters; 100 random initiation points; version 4.2.1). To keep the correlation pattern between "ancestral groups" and genotypes, we first simulated the control variable $C_2$ and then drew random genotype samples within $C_2$ subgroups. Note that all sampling processes were conducted with replacement and the sample size could be larger than 10,000.

After preparing the genotype data and control variables, we created $X$ and $Y$ according to the following procedures. The procedure for a continuous $X$ with sample size $n$ is as follows: (a) randomly select independent locations (LD $r^2$ < 0.05) for $n_e$ (200 or 1,000) effect SNPs; (b) generate the effect sizes according to $\begin{pmatrix} k_{1i} \\ k_{2i} \end{pmatrix} \overset{\text{i.i.d.}}{\sim} N\left(\begin{pmatrix} u_k \\ 2u_k \end{pmatrix}, \begin{pmatrix} s_k^2 & 2s_k^2\rho_k \\ 2s_k^2\rho_k & 4s_k^2 \end{pmatrix}\right), i = 1, \ldots, n_e$, where $u_k$ = 0, and $\rho_k$ is fixed to 0.9; (c) select an appropriate $s_k^2$, ensuring the sum of the SNP and control variable effects has an expected variance of 0.32 (calculated based on simulations; the variance of the control variable effect term is set to 0.09, with 0.03 assigned to $C_1$ and 0.06 assigned to $C_2$); (d) generate the residual $\varepsilon_x$ from $N(0,0.68)$; (e) sum the SNP effect, control variable effect, and residual together. Through the above procedure, we got an expected $R^2$ of 0.32 for the aggregate effect of SNPs and control variables. For a binary $X$, we simulated the linear part of **Eq. 11** similarly. We let the sum of the SNP and control variable effects have an expected variance of 0.73, the residual $\varepsilon_x$ follow $N(\mu_x, 0.09)$, and the sum of these two parts follow $N(-2.2,0.82)$ approximately. In this setting, about 0.8% of the probabilities were > 1. After forcing these values to be 1, we got an average Nagelkerke pseudo $R^2$ of 0.23 for the aggregate effect of SNPs and control variables.

The procedure for simulating $Y$ is as follows: (a) simulate pleiotropic effects according to **Formula 5** for the effect SNPs ($\rho = \rho_k$ = 0.9); (b) generate the control variable effect with the variance fixed to 0.09; (c) set $\beta_1$ and generate the exposure effect; (d) simulate the residual $k_x\varepsilon_x + \varepsilon_y$, where $\varepsilon_y \sim N(\mu_y, \sigma_{\varepsilon y}^2)$ is an independent component, and $k_x$, $\mu_y$, and $\sigma_{\varepsilon y}$ are selected to keep the linear part of **Eq. 2** approximately following $N(-2.5,0.78)$ (the variance of $k_x\varepsilon_x$ is set to 0.05; about 0.2% of the probabilities are > 1); (e) force the probabilities to be ≤ 1 and draw random samples for binary $Y$. Note that both the associations for binary $X$ and $Y$ may deviate from the log-linear model because a proportion of the simulated probabilities are > 1. Nevertheless, as these cases are rare, such deviations will exert little influence on individual effect estimators. We thus consider the log-linear model as a reasonable approximation.

As is mentioned previously, there are two major ways to obtain candidate SNPs for MR-PROLLIM. To lower the computational burden, we designed another procedure to imitate the first case in our simulations. For each replicate, we first simulated data of a reference GWAS. We then collected the effect SNPs and other SNPs in LD with them (LD $r^2$ > 0.5 for the case of

200 effect SNPs and > 0.9 for the case of 1,000 SNPs; LD matrix was calculated with the "snpgdsLDMat" function in R package "SNPRelate"; version 1.28.0 (10)). We ran linear (continuous $X$) or log-linear (binary $X$) regressions on these SNPs (plus control variables) and retained those having Wald $P$ < 1E-3 and selected by the LD clumping procedure (LD $r^2$ threshold = 0.05; clumping window = 10,000 kb; PLINK 1.90) as candidates. These candidate SNPs were checked again using the independent data that we simulated for formal MR analysis, with Wald $P$ = 1E-5 (random-effects MR-PROLLIM) or 1E-6 (other methods) as the threshold (i.e., the second-step $P$ threshold). The sample sizes for the reference GWAS and formal analysis were kept consistent. We did not take zero-effect SNPs into account because they could hardly pass two independent tests in series. We also neglected the SNPs weakly or moderately correlated with the effect SNPs under the consideration that they were unlikely to be selected as the representative SNPs during LD clumping.

**Section S18. Simplified simulation procedure for Case B**

It is possible that in MR practice, there may be several prominently strong SNPs (i.e., having relatively large effects on the exposure, while others, the majority, have small effects) that largely affect the final estimate, or only a few strong SNPs are eventually used.

A simplified simulation procedure was adopted for Case B. We generated SNP-exposure effects based on $\begin{pmatrix} k_{1,i} \\ k_{2,i} \end{pmatrix} \overset{\text{i.i.d.}}{\sim} N\left(\begin{pmatrix} 0 \\ 0 \end{pmatrix}, \begin{pmatrix} s_k^2 & 2s_k^2\rho_k \\ 2s_k^2\rho_k & 4s_k^2 \end{pmatrix}\right), i = 1, \dots, 20$ for three different sets of SNPs (sequentially, for a continuous exposure, $N_{\text{snp}} = 10, 5, 5$, $s_k = 0.11, 0.015, 0.015$, and $\rho_k = 0.7, 0.7, 1$; for a binary exposure, $N_{\text{snp}} = 10, 5, 5$, $s_k = 0.38, 0.04, 0.04$, and $\rho_k = 0.7, 0.7, 1$). We simulated according to the following data-generating process:

$$C \sim \text{Bernoulli}(0.5),$$

$$u_{10}, u_{20} \overset{\text{i.i.d.}}{\sim} N(0,1),$$

$$u_1 = \sqrt{0.96}u_{10} + 0.4C,$$

$$u_2 = \sqrt{0.5}u_{20} + \sqrt{0.5}u_1,$$

$$\begin{pmatrix} h_{1,i} \\ h_{2,i} \end{pmatrix} \overset{\text{i.i.d.}}{\sim} 0.5p_1 N\left(\begin{pmatrix} k_{1,i} \\ 2k_{1,i} \end{pmatrix}, \begin{pmatrix} s_h^2 & 2s_h^2\rho_h \\ 2s_h^2\rho_h & 4s_h^2 \end{pmatrix}\right) + 0.5p_1 N\left(\begin{pmatrix} 0 \\ 0 \end{pmatrix}, \begin{pmatrix} s_h^2 & 2s_h^2\rho_h \\ 2s_h^2\rho_h & 4s_h^2 \end{pmatrix}\right) + (1 - p_1)\begin{pmatrix} 0 \\ 0 \end{pmatrix},$$

$$s_h = 0.03 \text{ (continuous exposure) and } s_h = 0.06 \text{ (binary exposure)},$$

$$X = \mathbf{k}_1^T\mathbf{G}_1 + \mathbf{k}_2^T\mathbf{G}_2 + 0.2C + 0.7u_1 \text{ (continuous exposure)},$$

$$P_X = F(\exp(S(\mathbf{k}_1^T\mathbf{G}_1 + \mathbf{k}_2^T\mathbf{G}_2 + 0.2C + 0.5u_1) - 2.5)) \text{ (binary exposure)},$$

$$P_Y = F(\exp(S(\beta_1 X + \mathbf{h}_1^T\mathbf{G}_1 + \mathbf{h}_2^T\mathbf{G}_2 + 0.2C + 0.5u_2) - 2.3)),$$

where $\mathbf{G}_1$ and $\mathbf{G}_2$ are column vectors denoting the dummy variables for 20 SNPs, $S$ is a function that returns a centralized version of the input numbers, and $F$ denotes the function that forces the abnormal probabilities (i.e., values > 1) to be 1. We first fixed $\rho_h$ at 1 to support the PAHP assumption and generated 20 mutually independent SNPs for $n$ (60,000 for continuous exposures and 80,000 for binary exposures) individuals with the minor allele frequencies calculated as follows:

$$p_{0,i} \overset{\text{i.i.d.}}{\sim} U(0.15,0.35), k_{0,i} \overset{\text{i.i.d.}}{\sim} U(-0.1,0.1), i = 1,2,\dots,20,$$

$$p_{\text{MAF},i,j} = p_{0,i} + k_{0,i}(C_j - 0.5), j = 1,2,\dots,n.$$

We next removed the PAHP assumption by fixing $\rho_h$ at 0.7. To reduce the influence of abnormal probabilities, we additionally required the proportion of such probabilities to be less than 0.01. We repeated 500 times for each parameter setting. The simulation results are shown in **Fig. S3**.

**Section S19. Preparing and analyzing the UK Biobank data**

We first collected candidate SNPs for each trait according to GWAS summary data published by the Neale lab (GWAS round 2; http://www.nealelab.is/uk-biobank). The Neale lab had imposed an initial quality control on reported variants. We further required the candidates to satisfy the following criteria: (i) $P$ value < 1E-3 (for several traits, the threshold is 1E-5); (ii) MAF > 0.05; (iii) the SNP is located on autosomes; (iv) the SNP is recorded in the "snp151Common" database (http://hgdownload.cse.ucsc.edu/goldenPath/hg19/database/); (v) the SNPs are approximately independent (LD $r^2$ threshold = 0.1; clumping window = 10,000 kb; reference population of European ancestry from the 1000 Genomes Project; PLINK 1.90). This filtration generally resulted in several thousand candidate SNPs for each trait.

We restricted our analyses to Caucasian individuals and removed those judged as "outliers for heterozygosity or missing rate" or tagged with "sex chromosome aneuploidy". It has been reported that there exists a relatively high proportion of having close relatives among the UKB participants. Following the strategy by Bycroft *et al.* (11), we selected a largest independent set from these related individuals using the R function "largest_ivs" in package "igraph" (version 1.3.2) (12). Independence here was defined as kinship estimate < 0.0442, the default threshold recommended by KING to rule out third-degree or closer relatives (13). We next determined the exposures, outcomes, and control variables for the final sample. All disease traits were judged based on self-reported data, ICD-9, and ICD-10 codes. See **Table S12** for details.

Prior to a formal MR-PROLLIM analysis, we need to decide the second-step cutoff $P$ value for a further SNP filtration, as the first-step candidates may not always show strong SNP-exposure effects in our samples. Another key purpose of the second-step threshold is to control for the inflated type I error induced by multiple testing. We have proposed a parametric bootstrap method to estimate the probability of a zero-effect SNP passing two correlated tests in series (**Supplementary Text S16**). The results, together with the selected thresholds, are listed in **Table S13**. Note that we also implemented additional LD clumping procedures to apply a stricter requirement of independence (LD $r^2$ < 0.05) for formal analyses.

To make a more comprehensive comparison, we conducted summary statistic-based MR analyses using CAUSE. The GWAS summary data were mainly collected from the Neale lab (**Table S12**). Since the sample size and statistical model were not the same as ours, a direct comparison should be inappropriate. We thus added several other summary statistic-based methods as references, including the weighted median, weighted mode, and simple IVW. There was an approximately perfect sample overlap between the exposure and outcome GWAS datasets. We adopted $\hat{\rho}$ (i.e., estimate of the correlation coefficient) given by CAUSE to adjust for such overlaps, with standard errors of the individual effect estimators calculated as follows:

$$\sqrt{\frac{\mathrm{se}(\hat{\beta}_{YG})^2}{\hat{\beta}_{XG}^2}+\frac{\hat{\beta}_{YG}^2\mathrm{se}(\hat{\beta}_{XG})^2}{\hat{\beta}_{XG}^4}-\frac{2\hat{\rho}\mathrm{se}(\hat{\beta}_{XG})\mathrm{se}(\hat{\beta}_{YG})\hat{\beta}_{YG}}{\hat{\beta}_{XG}^3}}.$$

We used 1E-3 as the $P$ threshold for CAUSE and 1E-7 for other summary statistic-based methods.

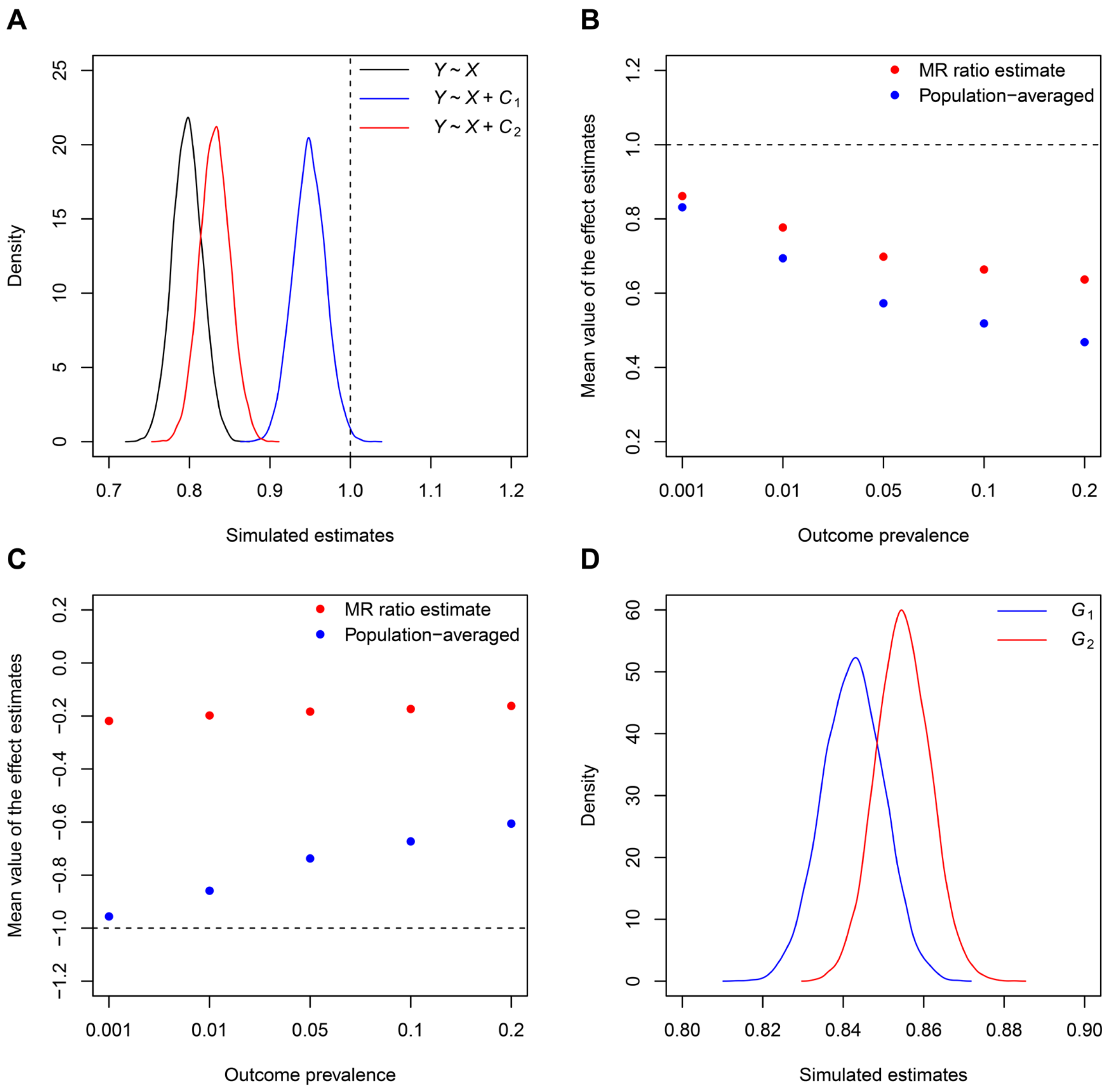


**Fig. S1. Simulation results illustrating the non-collapsibility of OR and its influence on MR.** (**A**) Simulation results for Example 1. Logistic regressions of $Y$ on $X$ and different control variables were conducted. The sample size was 50,000, and each case was repeated 10,000 times. The results indicate that the marginal logarithmic OR for $X$ varies with the distribution of the unmeasured explanatory variables even if no confounding effect exists. (**B**) Simulation results for Example 2. To reduce uncertainty, we set the sample size to one million. We repeated 10,000 times for the case of prevalence ≈ 0.001 (estimated based on simulations) and 1,000 times for other cases. The sampling error of each mean value is considered negligible (half width of the 95% confidence interval < 0.01). When the prevalence is about 1%, the simulated relative biases are -22.3% (relative to the conditional OR) and 11.9% (relative to the averaged OR), respectively. (**C**) Simulation results for Example 3. (**D**) Simulation results for Example 4. The sample size was set to 200,000, and each case was repeated 10,000 times. More details about these examples are provided in **Supplementary Text S1**.

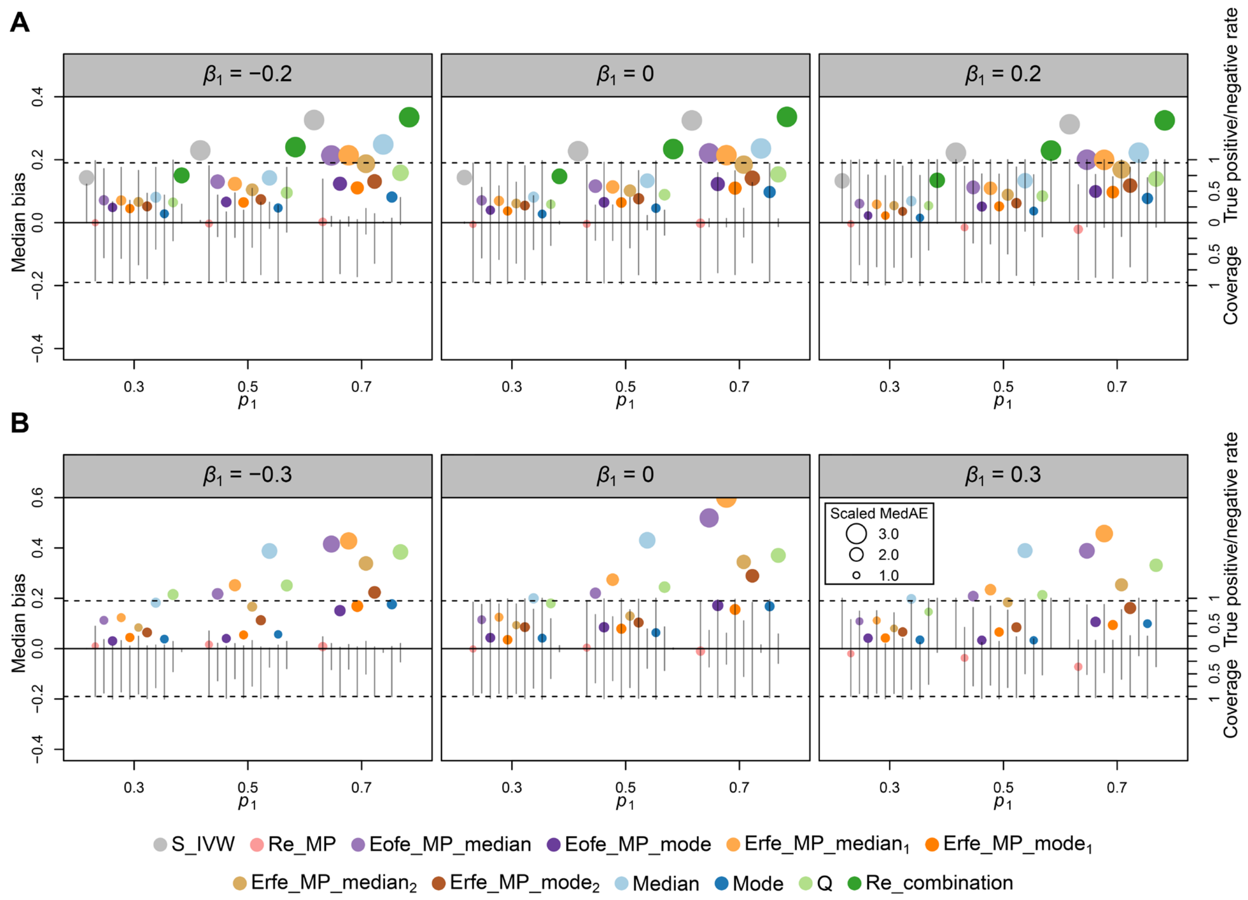


**Fig. S2. Statistical performances of different MR-PROLLIM methods in simulations with 1,000 effect SNPs.** (**A**) Statistical performance of MR-PROLLIM for continuous exposures. (**B**) Statistical performance of MR-PROLLIM for binary exposures. This figure shows the summary data of simulations conducted by generating 1,000 effect SNPs with the expected heritability held constant. The sample size was set to 80,000 for continuous exposures and 100,000 for binary exposures. See the caption of **Fig. 3** for more details.

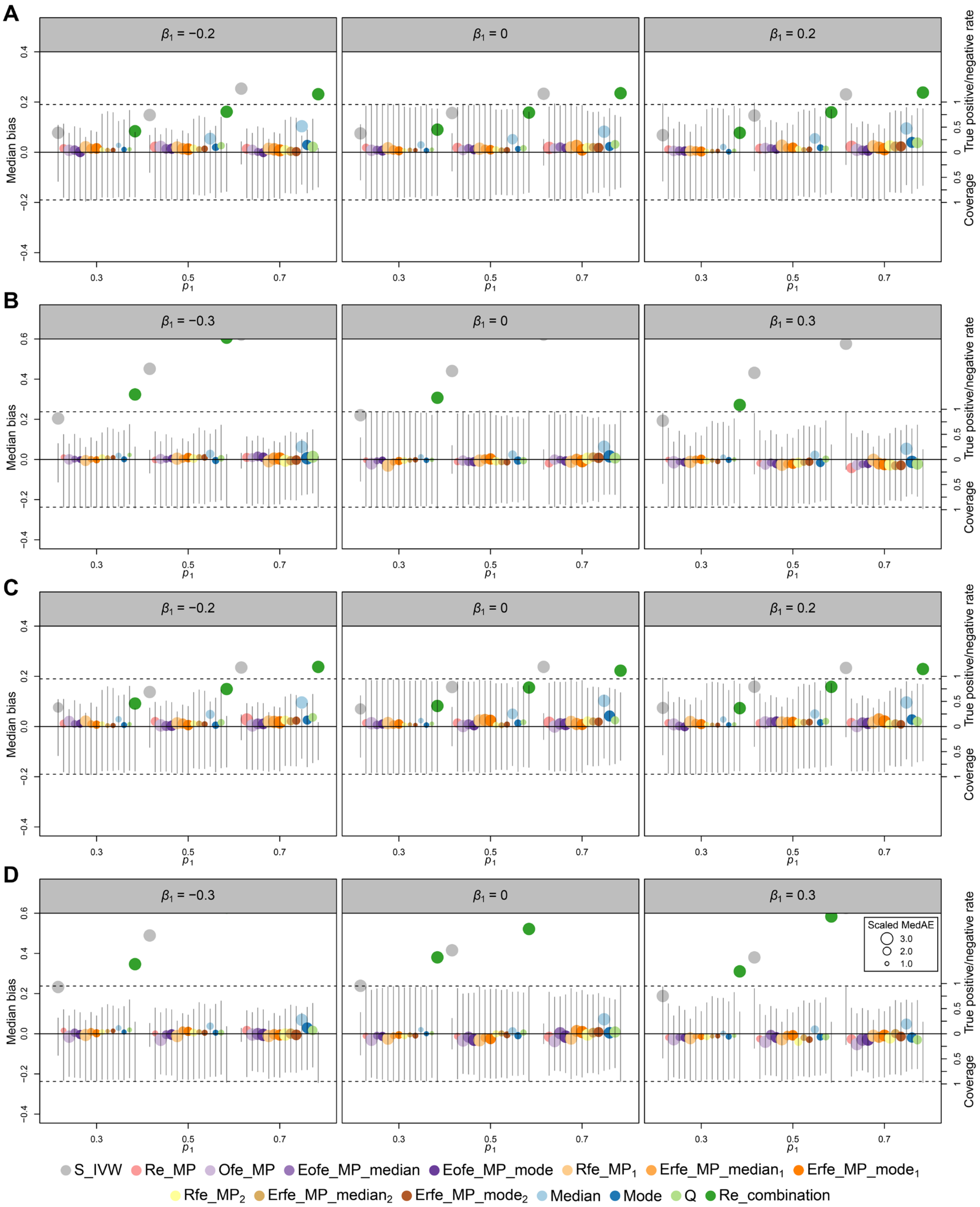


**Fig. S3. Statistical performances of different MR-PROLLIM methods in simulations with 20 effect SNPs.** (**A**) MR-PROLLIM for continuous exposures with the PAHP assumption. (**B**) MR-PROLLIM for binary exposures with the PAHP assumption. (**C**) MR-PROLLIM for continuous exposures without the PAHP assumption. (**D**) MR-PROLLIM for binary exposures without the PAHP assumption. See the caption of **Fig. 3** for more details. Ordinary fixed-effects MR-PROLLIM (Ofe_MP) and robust fixed-effects MR-PROLLIM (Rfe_MP) estimators, which do not suffer from false-negative judgments on the horizontal pleiotropy of unsuitable SNPs (**Materials and Methods**), are also included. These figures indicate that the ordinary fixed-effects MR-PROLLIM estimators (purple points) are relatively sensitive to the violation of PAHP, while the robust versions are less affected. Due to lack of SNPs, random-effects MR-PROLLIM tends to produce biased confidence intervals with actual coverage probabilities of around 90%.

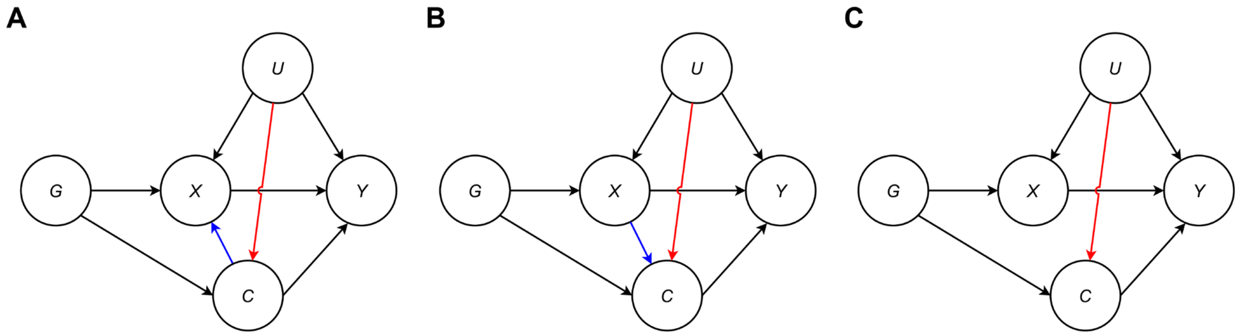


**Fig. S4. Directed acyclic graphs illustrating three subtypes of the control variables.** (**A**) Directed acyclic graph for Subtype A control variables. (**B**) Directed acyclic graph for Subtype B control variables. (**C**) Directed acyclic graph for Subtype C control variables.

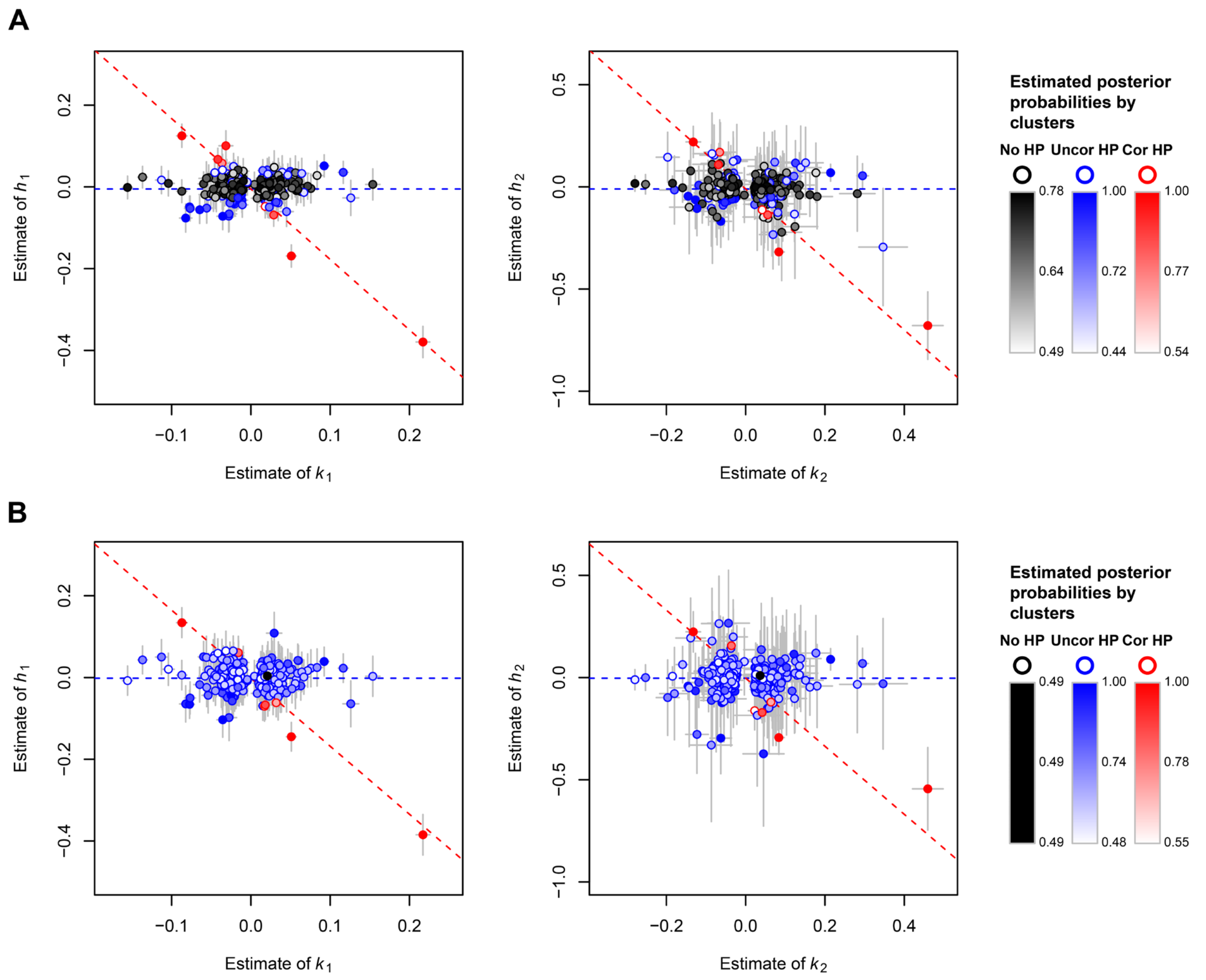


**Fig. S5. SNP classifications for the associations of HDL-C with angina and MI.** (**A**) SNP classifications for angina. (**B**) SNP classifications for MI. SNPs are classified according to maximum estimated posterior probabilities within the random-effects MR-PROLLIM framework and are distinguished by colors. For each trait pair, two subfigures are presented as MR-PROLLIM allows non-additive SNP effects. The variable $h_i$ $(i = 1, 2)$, which denotes a pleiotropic effect, equals $m_i - \beta_1 k_i$ for continuous exposures and $m_i - \ln\big((\exp(k_i) - 1)(1 - \exp(-\beta_1))\tilde{p} + 1\big)$ for binary exposures (**Materials and Methods**). The gray crosses indicate the 95% confidence intervals. The confidence intervals for $h_i$ are computed under the assumption that $\beta_1$ is estimated without error. The blue horizontal dashed lines mark the value of $\hat{\mu}_2$ (left) or $2\hat{\mu}_2$ (right), and the red dashed lines have a slope of $\hat{\mu}_1$ and an intercept of $\hat{\mu}_2$ (left) or $2\hat{\mu}_2$ (right). The constituent ratio of the inferred clusters is closely related to $\hat{p}_1$ and $\hat{p}_2$, but they do match exactly. For example, $1 - \hat{p}_1$ for MI is 0.30, but only one SNP (rs3996350) is classified into the cluster of no horizontal pleiotropy. Four SNPs (rs1065853, rs646776, rs429358, and rs11671872) are inferred to have correlated pleiotropy in both trait pairs. All of them were previously reported to have significant effects on LDL-C, TG, or other lipid traits according to GWAS Catalog records [https://www.ebi.ac.uk/gwas/; including records for SNPs in LD ($r^2 > 0.8$) with them]. And except for rs11671872, the other three were also linked to non-lipid traits, such as inflammatory, liver function, and cognitive measurements.

HP: horizontal pleiotropy; Uncor: uncorrelated; Cor: correlated.

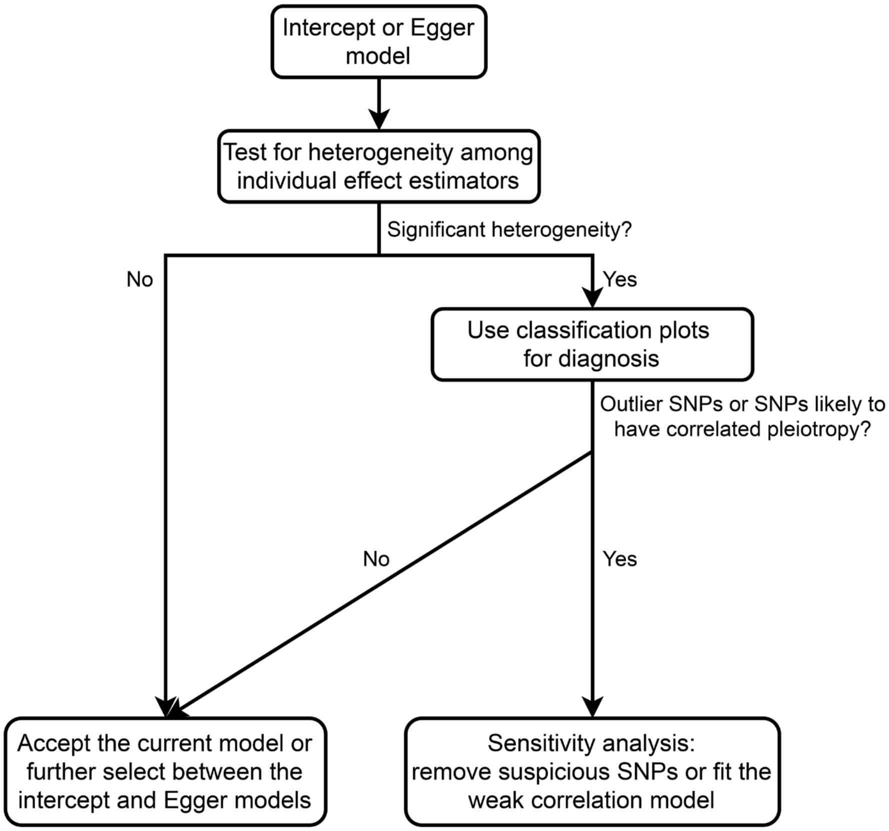


**Fig. S6. Procedure of *post hoc* analyses for the intercept and Egger models.** See **Supplementary Text S3** for details.

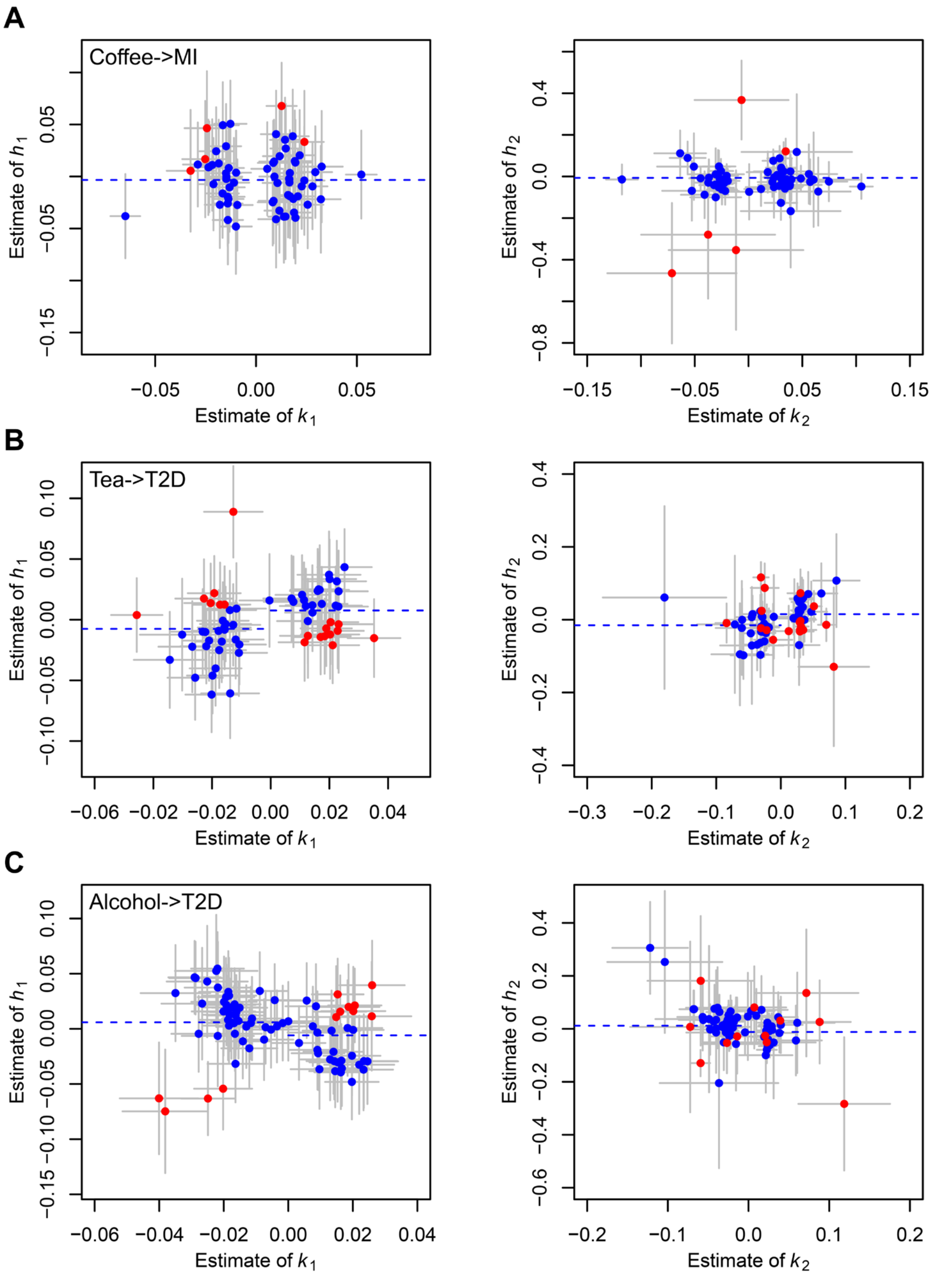


**Fig. S7. Classification plots for trait pairs with relatively obvious abnormalities.** (**A**) Classification plots for coffee and MI. (**B**) Classification plots for tea and T2D. (**C**) Classification plots for alcohol and T2D. Red points denote suspicious SNPs. Coffee and tea are inversely coded (i.e., $k_i > 0$ means less consumption; **Materials and Methods**). See **Fig. S5** for more details about the classification plots.

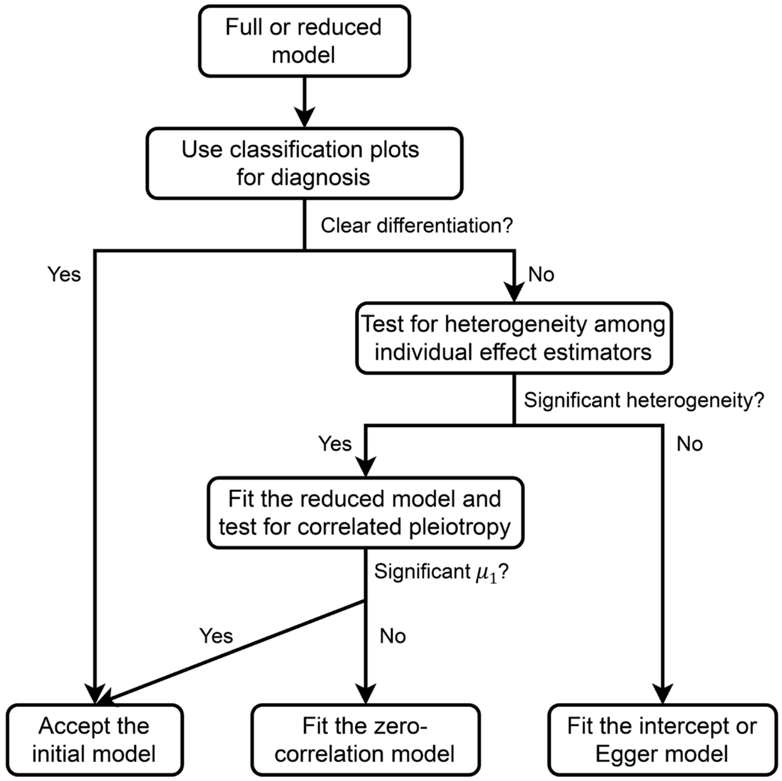


**Fig. S8. Procedure of *post hoc* diagnoses for the full and reduced models.** See **Supplementary Text S8** for details.

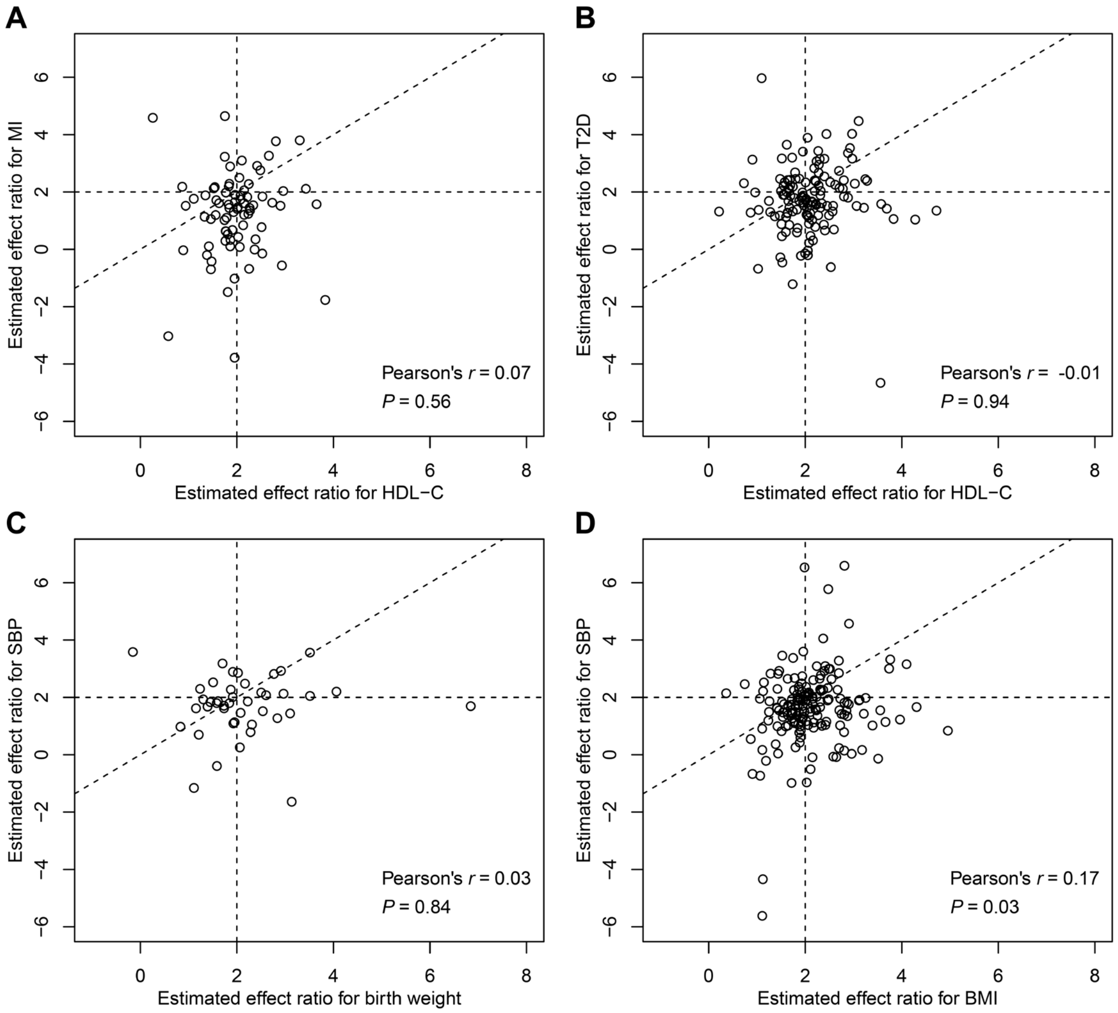


**Fig. S9. Associations between SNP-exposure and SNP-outcome effect ratios.** (**A–C**) Scatter plots for the considered non-causal trait pairs. Linear or log-linear regressions adjusted for sex, baseline age, and the first ten genetic principal components were conducted to derive the effect ratios. To improve robustness, we applied the following inclusion criteria: Wald $P$ values for SNP-exposure effects < 5E-6 (HDL-C) or < 5E-5 (birth weight); $P$ values for $\hat{k}_1 < 0.05$; Wald $P$ values for SNP-outcome effects < 0.05; and $P$ values for $\hat{h}_1 < 0.05$. We did not adopt the FDR-adjusted $P$ value because it would result in insufficient SNP numbers. Pearson's correlation coefficients (denoted as $r$) were used to measure the potential linear associations. The oblique dashed line in each plot has a slope of 1. (**D**) Scatter plot for the positive control. The same SNP inclusion criteria as for HDL-C were followed. The effect ratio correlation for this trait pair might be diluted by pleiotropic effects and measurement errors to some extent.

**Table S3. Sensitivity analyses for trait pairs with potential abnormalities detected by classification plots**

| Trait pair | Original results | | Removal of suspicious SNPs | | Weak correlation model | | Number of suspicious SNPs (%) |
|---|---|---|---|---|---|---|---|
| | **RR (95% CI)** | ***P* value** | **RR (95% CI)** | ***P* value** | **RR (95% CI)** | ***P* value** | |
| Coffee->MI | 0.70 (0.10, 1.30)[I] | 3.32E-01 | 0.80 (0.25, 1.36)[I] | 4.82E-01 | NA | NA | 5 (7.2) |
| Coffee->Heart failure | 1.55 (0.97, 2.14)[I] | 6.25E-02 | 1.43 (0.88, 1.99)[I] | 1.28E-01 | NA | NA | 4 (5.8) |
| Tea->T2D | 1.81 (0.00, 3.84)[E] | 4.31E-01 | 0.00 (0.00, 2.26)[E] | 2.94E-01 | 0.50 (0.00, 1.24) | 1.87E-01 | 18 (28.6) |
| Alcohol->T2D | 1.87 (0.61, 5.69)[E] | 2.70E-01 | 0.58 (0.30, 1.12)[E] | 1.06E-01 | 0.73 (0.50, 1.07) | 1.05E-01 | 12 (15.0) |
| Birth weight->Angina | 1.09 (0.60, 1.97)[E] | 7.86E-01 | 1.16 (0.69, 1.95)[E] | 5.84E-01 | NA | NA | 4 (2.7) |
| Birth weight->MI | 1.45 (0.68, 3.11)[E] | 3.37E-01 | 1.47 (0.74, 2.92)[E] | 2.67E-01 | NA | NA | 3 (2.0) |
| HDL-C->Arrhythmia | 0.85 (0.70, 1.03)[E] | 9.82E-02 | 0.96 (0.89, 1.03)[E] | 2.24E-01 | NA | NA | 2 (0.7) |
| HDL-C->Heart failure | 0.81 (0.56, 1.16)[E] | 2.51E-01 | 0.98 (0.85, 1.12)[E] | 7.46E-01 | NA | NA | 1 (0.3) |

See **Supplementary Text S3** for details.

**Table S4. Relative biases and significant detection rate of the simple IVW method by correlated pleiotropy status**

| Index | Trait pairs analyzed with the simple IVW method | | | *P* value |
|---|---|---|---|---|
| | **All (*N* = 83)** | **Significant correlated pleiotropy (*N* = 22)** | **Insignificant correlated pleiotropy (*N* = 61)** | |
| Relative bias (absolute value), median (range) | 0.223 (0.014, 1.780) | 0.292 (0.047, 1.780) | 0.179 (0.014, 0.570) | 0.03 |
| Significant detection, count (%) | 20/44 (45.5) | 9/11 (81.8) | 11/33 (33.3) | 0.01 |

Random-effects MR-PROLLIM was used to infer whether a trait pair had a non-zero effect (39 significant and 44 insignificant pairs; FDR-adjusted $P$ threshold = 0.1) and whether a trait pair was affected by correlated pleiotropy (22 significant and 61 insignificant pairs; unadjusted $P$ threshold = 0.05). Relative biases were calculated with the random-effects MR-PROLLIM results taken as the true values among the 39 trait pairs. "Significant detection" was defined as the unadjusted $P$ value given by the simple IVW method < 0.05 among the 44 trait pairs. These detections might contain false-positive ones. The absolute values of relative biases were compared using the Wilcoxon rank sum test. The significant detection rates were compared using Fisher's exact test.

**Table S5. Nominally significant detections of all MR methods by causal categories**

| Method | Causal category | Number of trait pairs | Number of significant detections | Proportion |
|---|---|---|---|---|
| Re_MP | | 31 | 29 | 0.935 |
| Eofe_MP_median | | 31 | 24 | 0.774 |
| Eofe_MP_mode | | 31 | 17 | 0.548 |
| Erfe_MP_median$_1$ | | 29 | 22 | 0.759 |
| Erfe_MP_mode$_1$ | | 29 | 17 | 0.586 |
| Erfe_MP_median$_2$ | | 29 | 21 | 0.724 |
| Erfe_MP_mode$_2$ | | 29 | 15 | 0.517 |
| Median$_1$ | Considered causal | 31 | 24 | 0.774 |
| Mode$_1$ | | 31 | 16 | 0.516 |
| Q | | 31 | 31 | 1.000 |
| Re_combination | | 31 | 27 | 0.871 |
| S_IVW$_1$ | | 31 | 31 | 1.000 |
| CAUSE | | 31 | 15 | 0.484 |
| Median$_2$ | | 31 | 22 | 0.710 |
| Mode$_2$ | | 31 | 16 | 0.516 |
| S_IVW$_2$ | | 31 | 26 | 0.839 |
| Re_MP | | 76 | 40 | 0.526 |
| Eofe_MP_median | | 74 | 38 | 0.514 |
| Eofe_MP_mode | | 74 | 26 | 0.351 |
| Erfe_MP_median$_1$ | | 72 | 35 | 0.486 |
| Erfe_MP_mode$_1$ | | 72 | 25 | 0.347 |
| Erfe_MP_median$_2$ | | 72 | 29 | 0.403 |
| Erfe_MP_mode$_2$ | | 72 | 23 | 0.319 |
| Median$_1$ | Considered causal and inconclusive | 76 | 35 | 0.461 |
| Mode$_1$ | | 76 | 24 | 0.316 |
| Q | | 76 | 50 | 0.658 |
| Re_combination | | 76 | 45 | 0.592 |
| S_IVW$_1$ | | 76 | 49 | 0.645 |
| CAUSE | | 76 | 17 | 0.224 |
| Median$_2$ | | 76 | 31 | 0.408 |
| Mode$_2$ | | 76 | 22 | 0.289 |
| S_IVW$_2$ | | 76 | 42 | 0.553 |
| Re_MP | | 22 | 0 | 0.000 |
| Eofe_MP_median | | 20 | 4 | 0.200 |
| Eofe_MP_mode | | 20 | 2 | 0.100 |
| Erfe_MP_median$_1$ | | 19 | 4 | 0.211 |
| Erfe_MP_mode$_1$ | | 19 | 2 | 0.105 |
| Erfe_MP_median$_2$ | | 19 | 2 | 0.105 |
| Erfe_MP_mode$_2$ | | 19 | 2 | 0.105 |
| Median$_1$ | Considered non-causal | 22 | 3 | 0.136 |
| Mode$_1$ | | 22 | 2 | 0.091 |
| Q | | 22 | 8 | 0.364 |
| Re_combination | | 22 | 11 | 0.500 |
| S_IVW$_1$ | | 22 | 12 | 0.545 |
| CAUSE | | 22 | 5 | 0.227 |
| Median$_2$ | | 22 | 5 | 0.227 |
| Mode$_2$ | | 22 | 4 | 0.182 |
| S_IVW$_2$ | | 22 | 14 | 0.636 |

Re_MP: Random-effects MR-PROLLIM;
Eofe_MP_median: Extended ordinary fixed-effects MR-PROLLIM median estimator;
Eofe_MP_mode: Extended ordinary fixed-effects MR-PROLLIM mode estimator;
Erfe_MP_median$_1$: Extended robust fixed-effects MR-PROLLIM median estimator (without first-stage simplification);
Erfe_MP_mode$_1$: Extended robust fixed-effects MR-PROLLIM mode estimator (without first-stage simplification);
Erfe_MP_median$_2$: Extended robust fixed-effects MR-PROLLIM median estimator (with first-stage simplification);
Erfe_MP_mode$_2$: Extended robust fixed-effects MR-PROLLIM mode estimator (with first-stage simplification);
Median$_1$: Classical weighted median estimator within MR-PROLLIM framework;
Mode$_1$: Classical weighted mode estimator within MR-PROLLIM framework;
Q: $Q$ statistic-based outlier removal within MR-PROLLIM framework;
Re_combination: Additive random-effects combination within MR-PROLLIM framework;
S_IVW$_1$: Simple inverse-variance weighted combination within MR-PROLLIM framework;
Median$_2$: Classical weighted median estimator using summary data;
Mode$_2$: Classical weighted mode estimator using summary data;
S_IVW$_2$: Simple inverse-variance weighted combination using summary data.

**Table S6. Comparison of the simple IVW results derived using MR-PROLLIM and the MR ratio method**

| Trait pair | Simple IVW (MR-PROLLIM) | | Simple IVW (traditional MR ratio) | | Ratio of log OR to log RR |
|---|---|---|---|---|---|
| | Log RR, 95% CI | *P* value | Log OR, 95% CI | *P* value | |
| Coffee->Arrhythmia | 0.132 (-0.180, 0.369) | 3.66E-01 | 0.035 (-0.038, 0.108) | 3.42E-01 | 0.268 |
| Coffee->Angina | 0.151 (-0.221, 0.422) | 3.75E-01 | 0.063 (-0.020, 0.147) | 1.36E-01 | 0.419 |
| Coffee->MI | -0.268 (-1.226, 0.212) | 3.28E-01 | -0.053 (-0.156, 0.050) | 3.15E-01 | 0.197 |
| Coffee->Heart failure | 0.554 (0.186, 0.823) | 6.73E-03 | 0.148 (0.031, 0.264) | 1.29E-02 | 0.267 |
| Coffee->Stroke | 0.165 (-0.408, 0.526) | 4.95E-01 | 0.052 (-0.057, 0.162) | 3.50E-01 | 0.319 |
| Coffee->T2D | 0.736 (0.539, 0.901) | 1.17E-08 | 0.263 (0.180, 0.346) | 4.74E-10 | 0.357 |
| Tea->Arrhythmia | -0.320 (-0.592, 0.056) | 8.67E-02 | -0.103 (-0.207, 0.001) | 5.12E-02 | 0.323 |
| Tea->MI | -0.444 (-0.794, 0.100) | 9.40E-02 | -0.089 (-0.235, 0.058) | 2.35E-01 | 0.200 |
| Tea->Stroke | -0.286 (-3.804, 0.392) | 5.04E-01 | -0.084 (-0.239, 0.072) | 2.91E-01 | 0.293 |
| Tea->T2D | -0.230 (-1.152, 0.241) | 3.99E-01 | -0.043 (-0.158, 0.071) | 4.61E-01 | 0.187 |
| Smoking->Arrhythmia | 0.291 (0.014, 0.508) | 4.08E-02 | 0.107 (0.042, 0.171) | 1.20E-03 | 0.366 |
| Smoking->Angina | 0.457 (0.149, 0.693) | 6.71E-03 | 0.136 (0.063, 0.210) | 2.90E-04 | 0.298 |
| Smoking->MI | 0.629 (0.256, 0.900) | 3.31E-03 | 0.214 (0.124, 0.304) | 3.47E-06 | 0.340 |
| Smoking->Heart failure | 0.361 (-0.230, 0.730) | 1.83E-01 | 0.148 (0.046, 0.251) | 4.66E-03 | 0.411 |
| Smoking->Stroke | 0.443 (0.009, 0.745) | 4.63E-02 | 0.172 (0.074, 0.270) | 5.75E-04 | 0.388 |
| Smoking->T2D | 0.502 (0.231, 0.715) | 1.12E-03 | 0.172 (0.101, 0.243) | 1.92E-06 | 0.343 |
| Nap->Arrhythmia | 0.217 (-0.108, 0.461) | 1.68E-01 | 0.010 (-0.053, 0.073) | 7.62E-01 | 0.045 |
| Nap->Angina | 0.580 (0.300, 0.799) | 4.11E-04 | 0.136 (0.065, 0.207) | 1.81E-04 | 0.234 |
| Nap->MI | 0.594 (0.225, 0.863) | 4.42E-03 | 0.092 (0.003, 0.181) | 4.36E-02 | 0.155 |
| Nap->Heart failure | 0.364 (-0.289, 0.756) | 2.12E-01 | 0.036 (-0.065, 0.137) | 4.87E-01 | 0.099 |
| Nap->Stroke | 0.488 (0.053, 0.790) | 3.19E-02 | 0.062 (-0.034, 0.159) | 2.03E-01 | 0.128 |
| Nap->T2D | 0.787 (0.549, 0.979) | 4.65E-07 | 0.186 (0.116, 0.256) | 1.79E-07 | 0.237 |
| Hypertension->Arrhythmia | 1.135 (1.033, 1.227) | 4.62E-43 | 0.265 (0.231, 0.299) | 1.44E-53 | 0.234 |
| Hypertension->Angina | 1.767 (1.664, 1.861) | 1.95E-61 | 0.426 (0.388, 0.464) | 1.98E-106 | 0.241 |
| Hypertension->MI | 1.677 (1.548, 1.792) | 2.81E-39 | 0.442 (0.394, 0.491) | 4.20E-72 | 0.264 |
| Hypertension->Heart failure | 1.682 (1.503, 1.834) | 2.14E-22 | 0.379 (0.325, 0.433) | 1.35E-42 | 0.225 |
| Hypertension->Stroke | 1.490 (1.350, 1.613) | 3.15E-31 | 0.398 (0.346, 0.450) | 8.51E-51 | 0.267 |
| Hypertension->T2D | 1.278 (1.112, 1.420) | 1.84E-20 | 0.237 (0.199, 0.274) | 2.60E-35 | 0.185 |

Coffee and tea consumptions were dichotomized according to the median values. Two trait pairs (Tea->Angina and Tea->Heart failure) were removed because the effect estimates given by these two methods were different in signs.

**Table S8. SNPs with non-additive effects on traits**

| Trait | Number of significant SNPs | Number of nominally non-additive SNPs | Proportion for nominally non-additive effects (95% CI) | Estimated proportion for non-additive effects (95% CI) |
|---|---|---|---|---|
| **Binary** | | | | 0.106 (0.073, 0.144) |
| Coffee | 78 | 11 | 0.141 (0.073, 0.238) | |
| Tea | 47 | 10 | 0.213 (0.107, 0.357) | |
| Smoking | 170 | 35 | 0.206 (0.148, 0.275) | |
| Nap | 169 | 14 | 0.083 (0.046, 0.135) | |
| **Continuous** | | | | 0.039 (0.031, 0.047) |
| Alcohol | 49 | 9 | 0.184 (0.088, 0.320) | |
| Birth weight | 206 | 30 | 0.146 (0.100, 0.201) | |
| BMI | 1180 | 84 | 0.071 (0.057, 0.087) | |
| SBP | 501 | 47 | 0.094 (0.070, 0.123) | |
| TG | 806 | 73 | 0.091 (0.072, 0.113) | |
| LDL-C | 576 | 64 | 0.111 (0.087, 0.140) | |
| HDL-C | 1010 | 74 | 0.073 (0.058, 0.091) | |
| Urate | 797 | 65 | 0.082 (0.064, 0.103) | |

Linear or log-linear regressions additionally adjusted for sex, baseline age, and the first ten genetic principal components were used to derive the data for this table. The proportions for non-additive effects are conservative estimates. See **Supplementary Text S4** for more details.

**Table S9. Associations between nominal classifications of the SNP-exposure and SNP-outcome effect ratios**

| SNP-exposure effect ratio | SNP-outcome effect ratio, ***N*** (%) | | | Total, ***N*** (%) | Fisher's ***P*** value |
|---|---|---|---|---|---|
| | $\hat{R} > 2$ | $\hat{R} = 2$ | $\hat{R} < 2$ | | |
| HDL-C->MI | | | | | 0.84 |
| $\hat{R} > 2$ | 0 (0.0) | 3 (100.0) | 0 (0.0) | 3 (2.9) | |
| $\hat{R} = 2$ | 1 (1.1) | 74 (78.7) | 19 (20.2) | 94 (90.4) | |
| $\hat{R} < 2$ | 0 (0.0) | 5 (71.4) | 2 (28.6) | 7 (6.7) | |
| HDL-C->T2D | | | | | 0.56 |
| $\hat{R} > 2$ | 0 (0.0) | 7 (87.5) | 1 (12.5) | 8 (5.0) | |
| $\hat{R} = 2$ | 0 (0.0) | 139 (93.3) | 10 (6.7) | 149 (93.1) | |
| $\hat{R} < 2$ | 0 (0.0) | 3 (100.0) | 0 (0.0) | 3 (1.9) | |
| Birth weight->SBP | | | | | 1.00 |
| $\hat{R} > 2$ | 0 (0.0) | 2 (100.0) | 0 (0.0) | 2 (3.5) | |
| $\hat{R} = 2$ | 0 (0.0) | 50 (94.3) | 3 (5.7) | 53 (93.0) | |
| $\hat{R} < 2$ | 0 (0.0) | 2 (100.0) | 0 (0.0) | 2 (3.5) | |
| BMI->SBP | | | | | 0.008 |
| $\hat{R} > 2$ | 2 (20.0) | 8 (80.0) | 0 (0.0) | 10 (4.5) | |
| $\hat{R} = 2$ | 4 (1.9) | 179 (85.2) | 27 (12.9) | 210 (94.2) | |
| $\hat{R} < 2$ | 0 (0.0) | 1 (33.3) | 2 (66.7) | 3 (1.3) | |

This table is analogous to **Fig. S9**. We classified the estimated effect ratios (denoted as $\hat{R}$) into three categories according to two-sided $P$ values (0.05 was adopted as the threshold). We additionally relaxed the $P$ thresholds for $\hat{k}_1$ and $\hat{h}_1$ from 0.05 to 0.2 to enlarge the SNP numbers. Without this step, the original Fisher's $P$ value for the positive control was 0.12, indicating the system might be underpowered. See **Supplementary Text S11** for more details**.**

**Table S10. Simulation results for correlations among individual effect estimators assuming no horizontal pleiotropy**

| Type of exposure | Sample size $n_1$ = 20,000 | | Sample size $n_2$ = 100,000 | |
|---|---|---|---|---|
| | Estimated $r$ (95% CI) | *P* value | Estimated $r$ (95% CI) | *P* value |
| Binary | -0.016 (-0.022, -0.010) | 4.29E-07 | -0.018 (-0.025, -0.012) | 5.33E-09 |
| Continuous | -0.000 (-0.007, 0.006) | 0.88 | -0.002 (-0.008, 0.004) | 0.55 |

A total of 100,000 random replicates were conducted to obtain the results for each case. Pearson's correlation coefficients (denoted as $r$) were adopted.

**Table S11. Summary of the recommended restrictions on the second-step *P* threshold**

| Recommended restrictions on the second-step threshold | Applicability | | Reasons |
|---|---|---|---|
| | Random-effects MP | Other methods | |
| $P_{c2} < \frac{P_{c1,\text{adj}}}{10}, P_{c1,\text{adj}} = 1 - \text{pchisq}\left(\frac{n_2}{n_1}\text{qchisq}(1 - P_{c1}, 2), 2\right)$ | Yes | No | Reduce the influence of two-step SNP selection on the posterior samples. |
| $P_{\text{fp}} < \frac{\alpha}{n_{\text{ind}}}$ | Yes* | Yes | Control for inflated type I error due to multiple testing. |
| $P_{c2} < 4.54\text{E} - 5$ | Yes* | Yes | Select nominally strong instrumental variables. |
| $n_{\text{SNP}} > 50$ | Yes | No | Obtain enough SNPs for random-effects MR-PROLLIM. |

* Random-effects MR-PROLLIM without the NOME assumption can account for zero-effect SNPs. Therefore, these restrictions may be relaxed.

**Table S13. Thresholds of $P$ values for two-step SNP selections**

| Trait | First-step $P$ threshold | Second-step $P$ threshold for re_MP | P_false_positive for re_MP | Second-step $P$ threshold for other methods | P_false_positive for other methods |
|---|---|---|---|---|---|
| Coffee | 1.00E-03 | 5.00E-05 | 7.85E-06 | 2.00E-06 | 5.28E-07 |
| Tea | 1.00E-03 | 5.00E-05 | 5.64E-06 | 5.00E-06 | 8.37E-07 |
| Smoking | 1.00E-03 | 1.00E-05 | 4.51E-06 | 2.00E-06 | 9.91E-07 |
| Alcohol | 1.00E-03 | 5.00E-04 | 3.66E-06 | 5.00E-05 | 5.21E-07 |
| Nap | 1.00E-03 | 1.00E-05 | 4.30E-06 | 1.00E-06 | 4.84E-07 |
| Birth weight | 1.00E-03 | 5.00E-06 | 2.44E-06 | 1.00E-06 | 5.28E-07 |
| BMI | 1.00E-05 | 1.00E-07 | 3.56E-08 | 1.00E-07 | 3.56E-08 |
| Hypertension | 1.00E-03 | 1.00E-07 | 3.60E-08 | 1.00E-07 | 3.60E-08 |
| TG | 1.00E-03 | 1.00E-05 | 4.47E-06 | 1.00E-06 | 5.06E-07 |
| LDL-C | 1.00E-03 | 1.00E-05 | 2.83E-06 | 2.00E-06 | 6.82E-07 |
| HDL-C | 1.00E-05 | 1.00E-07 | 1.28E-08 | 1.00E-07 | 1.28E-08 |
| Urate | 1.00E-05 | 1.00E-07 | 3.76E-08 | 1.00E-07 | 3.76E-08 |

The probability of a false positive detection by two tests in series (P_false_positive) was estimated with the proposed parametric bootstrap method [3,000 random SNPs; SNPs that show significant (FDR-adjusted $P < 0.1$) associations with the exposure are removed]. Following our SNP filtration criteria, we could at most obtain approximately 118 thousand independent SNPs from the GWAS summary data published by the Neale lab. This number was estimated based on LD clumping using PLINK (LD $r^2$ threshold = 0.05; clumping window = 10,000 kb; reference population of European ancestry from the 1000 Genomes Project; PLINK 1.90) with all $P$ values fixed to 0. To control for the inflated type I error induced by multiple testing, we generally need to keep the P_false_positive for other methods < 8.47E-07 ($\alpha$ = 0.1). Considering that most of the methods are robust to outliers, and the P_false_positive may be somewhat overestimated due to non-zero-effect SNPs, we do not set $\alpha$ here to 0.05.

**Legends for Tables S1, S2, S7, and S12**

Table S1. Main MR results and trait pair classifications

Table S2. Details and raw results for all trait pairs analyzed with MR-PROLLIM

Table S7. MR results for trait pairs with birth weight as the outcome

Table S12. Traits included in the real data MR analyses

### References


1. P. Ranganathan, R. Aggarwal, C. S. Pramesh, Common pitfalls in statistical analysis: Odds versus risk. *Perspect. Clin. Res.* **6**, 222-224 (2015).
2. E. C. Norton, B. E. Dowd, M. L. Maciejewski, Odds ratios-Current best practice and use. *JAMA* **320**, 84-85 (2018).
3. P. Cummings, The relative merits of risk ratios and odds ratios. *Arch. Pediatr. Adolesc. Med.* **163**, 438-445 (2009).
4. S. Vansteelandt, J. Bowden, M. Babanezhad, E. Goetghebeur, On instrumental variables estimation of causal odds ratios. *Stat. Sci.* **26**, 403-422 (2011).
5. J. Morrison, N. Knoblauch, J. H. Marcus, M. Stephens, X. He, Mendelian randomization accounting for correlated and uncorrelated pleiotropic effects using genome-wide summary statistics. *Nat. Genet.* **52**, 740-747 (2020).
6. D. A. Freedman, On the so-called "Huber Sandwich Estimator" and "Robust Standard Errors". *Am. Stat.* **60**, 299-302 (2006).
7. R. W. Farebrother, The distribution of a positive linear combination of chi-square random variables. *J. R. Stat. Soc. Ser. C-Appl. Stat.* **33**, 332-339 (1984).
8. W. R. Mebane, Jr., J. S. Sekhon, Genetic optimization using derivatives: The rgenoud package for R. *J. Stat. Softw.* **42**, 1-26 (2011).
9. M. P. Fay, B. I. Graubard, Small-sample adjustments for Wald-type tests using sandwich estimators. *Biometrics* **57**, 1198-1206 (2001).
10. X. Zheng *et al.*, A high-performance computing toolset for relatedness and principal component analysis of SNP data. *Bioinformatics* **28**, 3326-3328 (2012).
11. C. Bycroft *et al.*, The UK Biobank resource with deep phenotyping and genomic data. *Nature* **562**, 203-209 (2018).
12. G. Csardi, T. Nepusz, The igraph software package for complex network research. *InterJ. Compl. Syst.* **1695**, 1-9 (2006).
13. A. Manichaikul *et al.*, Robust relationship inference in genome-wide association studies. *Bioinformatics* **26**, 2867-2873 (2010).
